# Many but not all deep neural network audio models capture brain responses and exhibit correspondence between model stages and brain regions

**DOI:** 10.1101/2022.09.06.506680

**Authors:** Greta Tuckute, Jenelle Feather, Dana Boebinger, Josh H. McDermott

## Abstract

Models that predict brain responses to stimuli provide one measure of understanding of a sensory system, and have many potential applications in science and engineering. Deep artificial neural networks have emerged as the leading such predictive models of the visual system, but are less explored in audition. Prior work provided examples of audio-trained neural networks that produced good predictions of auditory cortical fMRI responses and exhibited correspondence between model stages and brain regions, but left it unclear whether these results generalize to other neural network models, and thus how to further improve models in this domain. We evaluated model-brain correspondence for publicly available audio neural network models along with in-house models trained on four different tasks. Most tested models out-predicted previous filter-bank models of auditory cortex, and exhibited systematic model-brain correspondence: middle stages best predicted primary auditory cortex while deep stages best predicted non-primary cortex. However, some state-of-the-art models produced substantially worse brain predictions. Models trained to recognize speech in background noise produced better brain predictions than models trained to recognize speech in quiet, potentially because hearing in noise imposes constraints on biological auditory representations. The training task influenced the prediction quality for specific cortical tuning properties, with best overall predictions resulting from models trained on multiple tasks. The results generally support the promise of deep neural networks as models of audition, though they also indicate that current models do not explain auditory cortical responses in their entirety.

## Introduction

An overarching aim of neuroscience is to build quantitatively accurate computational models of sensory systems. Success entails models that take sensory signals as input and reproduce the behavioral judgments mediated by a sensory system as well as its internal representations. A model that can replicate behavior and brain responses for arbitrary stimuli would help validate the theories that underlie the model, but would also have a host of important applications. For instance, such models could guide brain-machine interfaces by specifying patterns of brain stimulation needed to elicit particular percepts or behavioral responses.

One approach to model building is to construct machine systems that solve biologically relevant tasks, based on the hypothesis that task constraints may cause them to reproduce the characteristics of biological systems^1, 2^. Advances in machine learning have stimulated a wave of renewed interest in this model building approach. Specifically, deep artificial neural networks (DNNs) now achieve human-level performance on real-world classification tasks such as object and speech recognition, yielding a new generation of candidate models in vision, audition, language, and other domains^3–8^. DNN models are relatively well explored within vision, where they reproduce some patterns of human behavior^9–12^ and in many cases appear to replicate aspects of the hierarchical organization of the primate ventral stream^13–16^. These and other findings are consistent with the idea that brain representations are constrained by the demands of the tasks organisms must carry out, such that optimizing for ecologically relevant tasks produces better models of the brain in a variety of respects.

These modeling successes have been accompanied by striking examples of model behaviors that deviate from those of humans. For instance, current neural network models are often vulnerable to adversarial perturbations – targeted changes to the input that are imperceptible to humans, but which change the classification decisions of a model^17–20^. Current models also often do not generalize to stimulus manipulations to which human recognition is robust, such as additive noise or translations of the input^12, 21–24^. Models also typically exhibit invariances that humans lack, such that model metamers – stimuli that produce very similar responses in a model – are typically not recognizable as the same object class to humans^25–27^. And efforts to compare models to classical perceptual effects exhibit a mixture of successes and failures, with some human perceptual phenomena missing from the models^28, 29^. The causes and significance of these model failures remain an active area of investigation and debate^30^.

Alongside the wave of interest within human vision, deep neural network models have also stimulated research in audition. Comparisons of human and model behavioral characteristics have found that audio-trained neural networks often reproduce patterns of human behavior when optimized for naturalistic tasks and stimulus sets^31–35^. Several studies have also compared audio-trained neural networks to brain responses within the auditory system^31, 36, 37–43^. The best-known of these prior studies is arguably that of Kell et al., (2018), who found that DNNs jointly optimized for speech and music classification could predict functional magnetic resonance imaging (fMRI) responses to natural sounds in auditory cortex substantially better than a standard model based on spectrotemporal filters. In addition, model stages exhibited correspondence with brain regions, with middle stages best predicting primary auditory cortex and deeper stages best predicting non-primary auditory cortex. However, Kell et al. used only a fixed set of two tasks, investigated a single class of model, and relied exclusively on regression-derived predictions as the metric of model-brain similarity.

Several subsequent studies built on these findings by analyzing models trained on various speech-related tasks, and found they were able to predict cortical responses to speech better than chance, with some evidence that different model stages best predicted different brain regions^40,41,42,43^. Another recent study examined models trained on sound recognition tasks, finding better predictions of brain responses and perceptual dissimilarity ratings when compared to traditional acoustic models^44^. But each of these studies analyzed only a small number of models, and each used a different brain dataset, making it difficult to compare results across studies, and leaving the generality of brain-DNN similarities unclear. Specifically, it has remained unclear whether DNNs trained on other tasks and sounds also produce good predictions of brain responses, whether the correspondence between model stages and brain regions is consistent across models, and whether the training task critically influences the ability to predict responses in particular parts of auditory cortex. These questions are important for substantiating the hierarchical organization of the auditory cortex (by testing whether distinct stages of computational models best map onto different regions of the auditory system), for understanding the role of tasks in shaping cortical representations (by testing whether optimization for particular tasks produces representations that match those of the brain), and for guiding the development of better models of the auditory system (by helping to understand the factors that enable a model to predict brain responses).

To answer these questions, we examined brain-DNN similarities within the auditory cortex for a large set of models. To address the generality of brain-DNN similarities, we tested a large set of publicly available audio-trained neural network models, trained on a wide variety of tasks and spanning many types of models. To address the effect of training task, we supplemented these publicly available models with in-house models trained on four different tasks. We evaluated both the overall quality of the brain predictions as compared to a standard baseline spectrotemporal filter model of the auditory cortex^45^, as well as the correspondence between model stages and brain regions. To ensure that the general conclusions were robust to the choice of model-brain similarity metric, wherever possible we used two different metrics: the variance explained by linear mappings fit from model features to brain responses^46^, and representational similarity analysis^47^ (noting that these two metrics evaluate distinct inferences about what might be similar between two representations^48, 49^). We used two different fMRI datasets to assess the reproducibility and robustness of the results: the original dataset (Norman-Haignere et al., 2015, n=8) used in Kell et al., to facilitate comparisons to those earlier results, as well as a second recent dataset (Boebinger et al., 2021, n=20) with data from a total of 28 unique participants. We analyzed auditory cortical brain responses, as subcortical responses are challenging to measure with the necessary reliability (and hence were not included in the datasets we analyzed).

We found that most DNN models produced better predictions of brain responses than the baseline model of the auditory cortex. In addition, most models exhibited a correspondence between model stages and brain regions, with lateral, anterior, and posterior non-primary auditory cortex being better predicted by deeper model stages. Both of these findings indicate that many such models provide better descriptions of cortical responses than traditional filter-bank models of auditory cortex. However, not all models produced good predictions, suggesting that some training tasks and architectures yield better brain predictions than others. We observed effects of the training data, with models trained to hear in noise producing better brain predictions than those trained exclusively in quiet. We also observed significant effects of the training task on the predictions of speech, music, and pitch-related cortical responses. The best overall predictions were produced by models trained on multiple tasks. The results replicated across both fMRI datasets and with representational similarity analysis. The results indicate that many DNNs replicate aspects of auditory cortical representations, but indicate the important role of training data and tasks in obtaining models that yield accurate brain predictions, in turn consistent with the idea that auditory cortical tuning has been shaped by the demands of having to support auditory behavior.

## Results

### Deep neural network modeling overview

The artificial neural network models considered here take an audio signal as input and transform it via cascades of operations loosely inspired by biology: filtering, pooling, and normalization, among others. Each stage of operations produces a representation of the audio input, typically culminating in an output stage: a set of units whose activations can be interpreted as the probability that the input belongs to a particular class (e.g., a spoken word, or phoneme, or sound category).

A model is defined by its “architecture” – the arrangement of operations within the model – and by the parameters of each operation that may be learned during training. These parameters are typically initialized randomly, and are then optimized via gradient descent to minimize a loss function over a set of training data. The loss function is typically designed to quantify performance of a task. For instance, training data might consist of a set of speech recordings that have been annotated, the model’s output units might correspond to word labels, and the loss function might quantify the accuracy of the model’s word labeling compared to the annotations. The optimization that occurs during training would cause the model’s word labeling to become progressively more accurate.

A model’s performance is a function of both the architecture and the training procedure; training is thus typically conducted alongside a search over the space of model architectures to find an architecture that performs the training task well. Once trained, a model can be applied to any arbitrary stimulus, yielding a decision (if trained to classify its input) that can be compared to the decisions of human observers, along with internal model responses that can be compared to brain responses. Here we focus on the internal model responses, comparing them to fMRI responses in human auditory cortex, with the goal of assessing whether the representations derived from the model reproduce aspects of representations in the auditory cortex as evaluated by two commonly used metrics.

### Model selection

We began by compiling a set of models that we could compare to brain data (see “Candidate models” in Methods for full details and Tables 1 and 2 in Methods for an overview). Two criteria dictated the choice of models. First, we sought to survey a wide range of models to assess the generality with which deep neural networks would be able to model auditory cortical responses. Second, we wanted to explore effects of the training task. The main constraint on the model set was that there were relatively few publicly available audio-trained deep neural network models available at the time of this study (in part because much work on audio engineering is done in industry settings where models and datasets are not made public). We thus included every model we could obtain in a PyTorch implementation that had been trained on some sort of large-scale audio task (i.e., we neglected models trained to classify spoken digits, or other tasks with small numbers of classes, on the grounds that such tasks are unlikely to place strong constraints on the model representations^52, 53^). The PyTorch constraint resulted in the exclusion of 3 models that were otherwise available at the time of the experiments (see Methods). The resulting set of 9 models varied in both their architecture (spanning convolutional neural networks, recurrent neural networks, and transformers) and training task (ranging from automatic speech recognition and speech enhancement to audio captioning and audio source separation).

**Table 1.** External model overview.

| Model name | Brief description | Model input | Model output | Training dataset |
| --- | --- | --- | --- | --- |
| AST (Audio Spectrogram Transformer)<sup>142</sup> | Transformer architecture for audio classification. | Spectrogram | AudioSet label (527) | AudioSet (ImageNet pretraining)<sup>55,124</sup> |
| DCASE2020<sup>143</sup> | Recurrent network trained for automated audio captioning. | Spectrogram | Audio text captions (4,367) | Clotho V1<sup>144</sup> |
| DeepSpeech2<sup>145</sup> | Recurrent architecture for automatic speech recognition. | Spectrogram | Characters (29) | LibriSpeech<sup>146</sup> |
| MetricGAN<sup>147</sup> | Generative adversarial network for speech enhancement. | Spectrogram | Voice-enhanced audio | VoiceBank-DEMAND<sup>148</sup> |
| S2T (Speech-to-Text)<sup>149</sup> | Transformer architecture for automatic speech recognition and speech-to-text translation. | Spectrogram | Words (10,000) | LibriSpeech<sup>146</sup> |
| SepFormer (Separation Transformer)<sup>150</sup> | Transformer architecture for speech separation. | Waveform | Source-separated audio | WHAMR!<sup>151</sup> |
| VGGish<sup>152</sup> | Convolutional architecture for audio classification. | Spectrogram | Video label (30,871) | YouTube-100M<sup>152</sup> |
| VQ-VAE (ZeroSpeech2020)<sup>153</sup> | Convolutional encoder architecture for speech synthesis in a target speaker's voice. | Spectrogram | Audio in target speaker's voice | ZeroSpeech 2019 training dataset<sup>154</sup> |
| Wav2vec2<sup>129</sup> | Transformer architecture for automatic speech recognition. | Waveform | Characters (32) | LibriSpeech<sup>146</sup> |

**Table 2.**
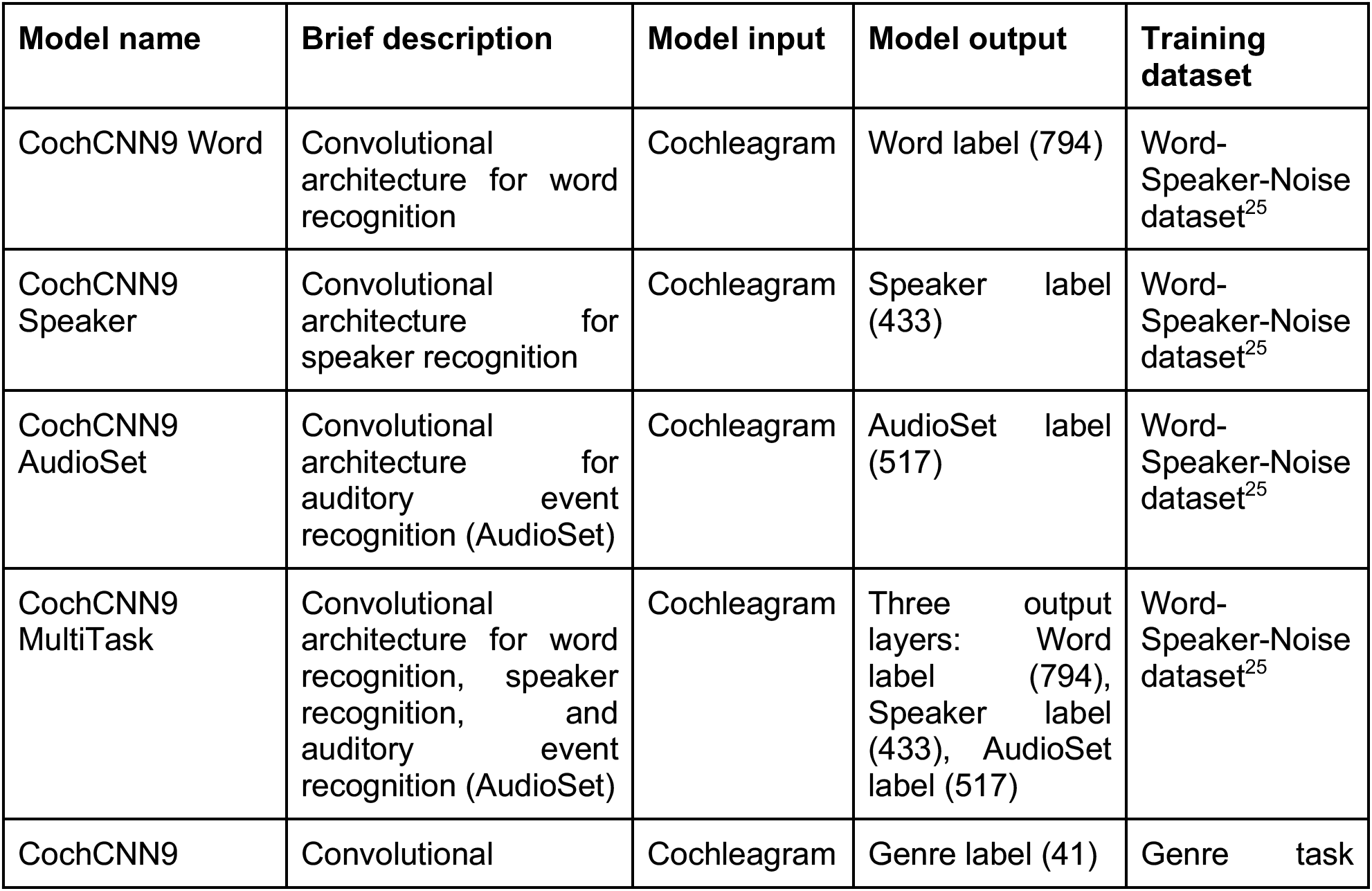
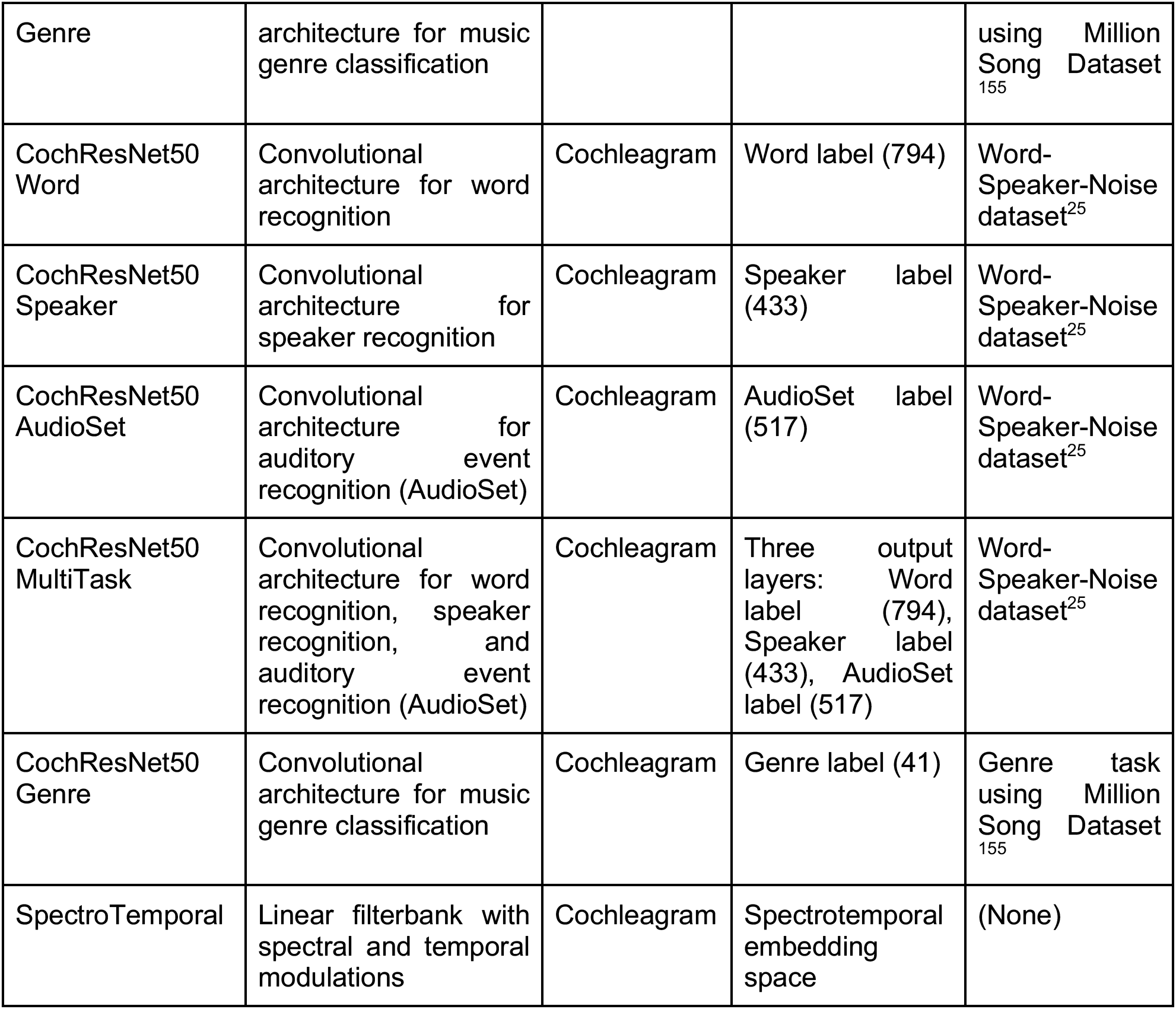
In-house model overview.

To supplement these external models, we trained ten models ourselves: two architectures trained separately on each of four tasks as well as on three of the tasks simultaneously. We used the three tasks that could be implemented using the same dataset (where each sound clip had labels for words, speakers, and audio events). One of the architectures we used was similar to that used in our earlier study (Kell et al., 2018), which identified a candidate architecture from a large search over number of stages, location of pooling, and size of convolutional filters. The model was selected entirely based on performance on the training tasks (i.e., word and music genre recognition). The resulting model performed well on both word and music genre recognition, and was more predictive of brain responses to natural sounds than a set of alternative neural network architectures as well as a baseline model of auditory cortex. This in-house architecture (henceforth CochCNN9) consisted of a sequence of convolutional, normalization, and pooling stages preceded by a hand-designed model of the cochlea (henceforth termed a ‘cochleagram’). The second in-house architecture was a ResNet50^54^ backbone with a cochleagram front end (henceforth CochResNet50). CochResNet50 was a much deeper model than CochCNN9 (50 layers compared to 9 layers) with residual (skip-layer) connections, and although this architecture was not determined via an explicit architecture search for auditory tasks, it was developed for computer vision tasks^54^, and outperformed CochCNN9 on the training tasks (see Methods; Candidate models). We used two architectures to obtain a sense of the consistency of any effects of task that we might observe.

The four in-house training tasks consisted of recognizing words, speakers, audio events (labeled clips from the AudioSet^55^ dataset, consisting of human and animal sounds, excerpts of various musical instruments and genres, and environmental sounds), or musical genres from audio (referred to henceforth as Word, Speaker, AudioSet and Genre, respectively). The multi-task models had three different output layers, one for each included task (Word, Speaker, and AudioSet), connected to the same network. The three tasks for the multi-task network were originally chosen because we could train on all of them simultaneously using a single existing dataset (the Word-Speaker-Noise dataset^25^) in which each clip has three associated labels: a word, a speaker, and a background sound (from AudioSet). For the single-task networks, we used one of these three sets of labels. We additionally trained models with a fourth task – a music-genre classification task originally presented in Kell et al., (2018), that used a distinct training set. As it turned out, the first three tasks individually produced better brain predictions than the fourth, and the multi-task model produced better predictions than any of the models individually, and so we did not explore additional combinations of tasks. These in-house models were intended to allow a controlled analysis of the effect of task, to complement the all-inclusive but uncontrolled set of external models.

We compared each of these models to an untrained baseline model that is commonly used in cognitive neuroscience^45^. The baseline model consisted of a set of spectrotemporal modulation filters applied to a model of the cochlea (henceforth referred to as the SpectoTemporal model). The SpectroTemporal baseline model was explicitly constructed to capture tuning properties observed in the auditory cortex, and previously been found to account for auditory cortical responses to some extent^56^, particularly in primary auditory cortex^57^, and thus provided a strong baseline for model comparison.

### Brain data

To assess the replicability and robustness of the results, we evaluated the models on two independent fMRI datasets (each with three scanning sessions per participant). Each presented the same set of 165 two-second natural sounds to human listeners. One experiment^50^ collected data from 8 participants with moderate amounts of musical experience (henceforth <u>NH2015</u>). This dataset was analyzed in a previous study investigating deep neural network predictions of fMRI responses^31^. The second experiment^51^ collected data from a different set of 20 participants, 10 of whom had almost no musical experience, and 10 of whom had extensive musical training (henceforth <u>B2021</u>). The fMRI experiments measured the blood-oxygen-level-dependent (BOLD) response to each sound in each voxel in the auditory cortex of each participant (including all temporal lobe voxels that responded significantly more to sound than silence, and whose test-retest response reliability exceeded a criterion; see Methods; fMRI data). We note that the natural sounds used in the fMRI experiment, with which we evaluated model-brain correspondence, were not part of the training data for the models, nor were they drawn from the same distribution as the training data.

### General approach to analysis

Because the sounds were short relative to the time constant of the fMRI BOLD signal, we summarized the fMRI response from each voxel as a single scalar value for each sound. The primary similarity metric we used was the variance in these voxel responses that could be explained by linear mappings from the model responses, obtained via regression. This regression analysis has the advantage of being in widespread use^31, 46, 56, 58,59,60^ and hence facilitates comparison of results to related work. We supplemented the regression analysis with a representational similarity analysis^47^, and wherever possible present results from both metrics.

The steps involved in the regression analysis are shown in Figure 1A. Each sound was passed through a neural network model, and the unit activations from each network stage were used to predict the response of individual voxels (after averaging unit activations over time to mimic the slow time constant of the BOLD signal). Predictions were generated with cross-validated ridge regression, using methods similar to those of many previous studies using encoding models of fMRI measurements^31, 46, 56, 58,59,60^. Regression yields a linear mapping that rotates and scales the model responses to best align them to the brain response, as is needed to compare responses in two different systems (model and brain, or two different brains or models). A model that reproduces brain-like representations should yield similar patterns of response variation across stimuli once such a linear transform has been applied (thus “explaining” a large amount of the brain response variation across stimuli).

**Figure 1.**
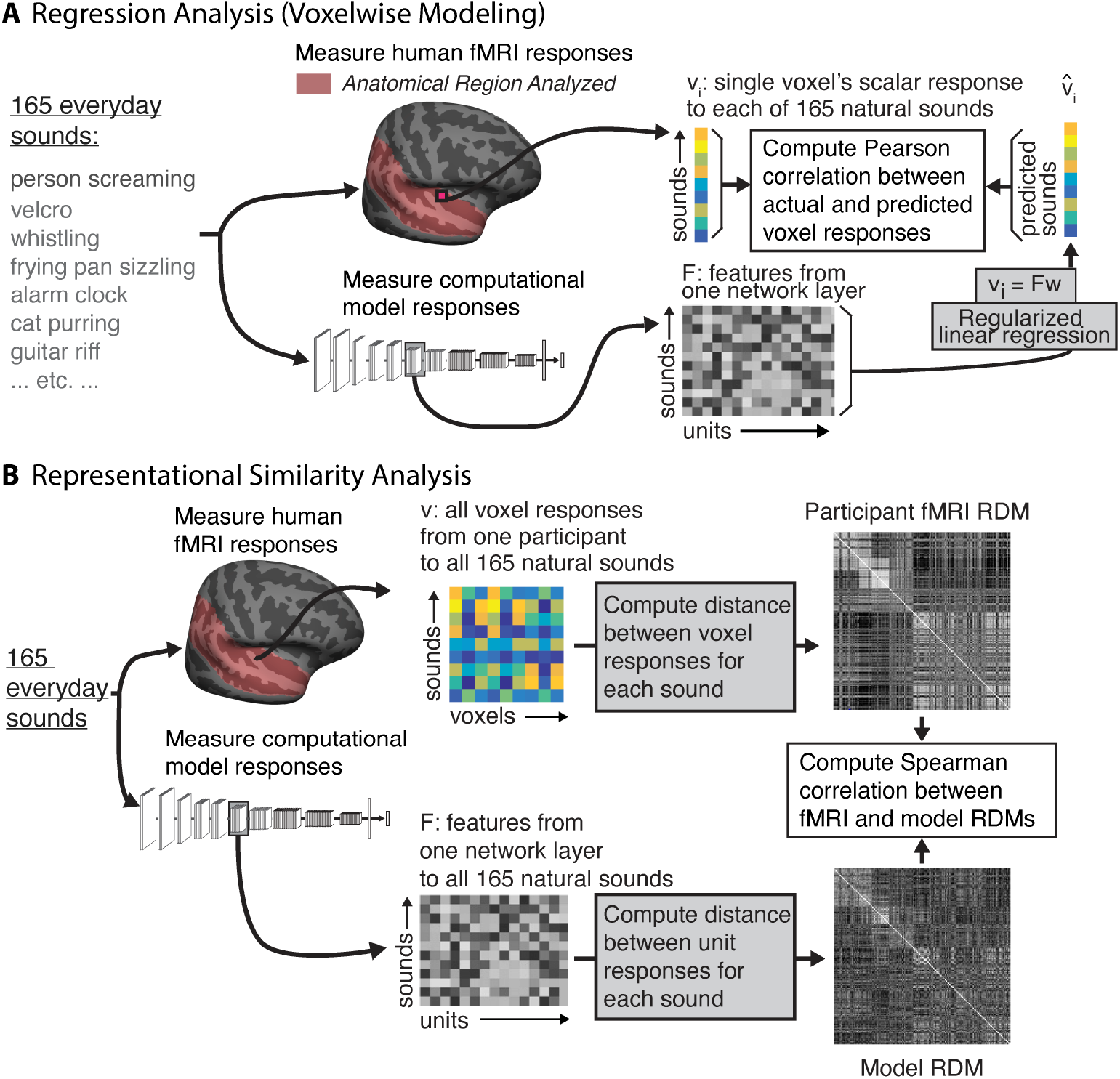
Analysis method. **(A)** Regression analysis (voxelwise modeling). Brain activity of human participants (n=8, n=20) was recorded while they listened to a set of 165 natural sounds in fMRI. Data were taken from two previous publications^50, 51^. We then presented the same set of 165 sounds to each model, measuring the time-averaged unit activations from each model stage in response to each sound. We performed an encoding analysis where voxel activity was predicted by a regularized linear model of the DNN activity. We modeled each voxel as a linear combination of model units from a given model stage, estimating the linear transform with half (n=83) the sounds and measuring the prediction quality by correlating the empirical and predicted response to the left-out sounds (n=82) using the Pearson correlation. We performed this procedure for 10 random splits of the sounds. Figure adapted from Kell et al.^31^. **(B)** Representational Similarity Analysis. We used the set of brain data and model activations described for the voxelwise regression modeling. We constructed a representational dissimilarity matrix (RDM) from the fMRI responses by computing the distance (1-Pearson correlation) between all voxel responses to each pair of sounds. We similarly constructed an RDM from the unit responses from a model stage to each pair of sounds. We measured the Spearman correlation between the fMRI and model RDMs as the metric of model-brain similarity. When reporting this correlation from a best model stage, we used 10 random splits of sounds, choosing the best stage from the training set of 83 sounds and measuring the Spearman correlation for the remaining set of 82 test sounds. The fMRI RDM is the average RDM across all participants for all voxels and all sounds in NH2015. The model RDM is from an example model stage (ResNetBlock_2 of the CochResNet50-MultiTask network).

The specific approach here was modeled after that of Kell et al., 2018: we used 83 of the sounds to fit the linear mapping from model units to a voxel’s response, and then evaluated the predictions on the 82 remaining sounds, taking the median across 10 training/test cross-validation splits, and correcting for both the reliability of the measured voxel response and the reliability of the predicted voxel response^61, 62^. The variance explained by a model stage was taken as a metric of the brain-likeness of the model representations. We asked i) to what extent the models in our set were able to predict brain data, and ii) whether there was a relationship between stages in a model and regions in the human brain. We performed the same analysis on the SpectroTemporal baseline model for comparison.

To assess the robustness of our overall conclusions to the evaluation metric, we also performed Representational Similarity Analysis to compare the representational geometries between brain and model responses (Figure 1B). We first measured representational dissimilarity matrices (RDMs) for a set of voxel responses from the Pearson correlation of all the voxel responses to one sound with that for another sound. These correlations for all pairs of sounds yields a matrix, which is standardly expressed as 1-C, where C is the correlation matrix. When computed from all voxels in the auditory cortex, this matrix is highly structured, with some pairs of sounds producing much more similar responses than others (Supplementary Figure S1). We then analogously measured this matrix from the time-averaged unit responses within a model stage. To assess whether the representational geometry captured by these matrices was similar between a model and the brain, we measured the Spearman correlation between the brain and model RDMs. As in previous work^63, 64^, we did not correct this metric for the reliability of the RDMs, but instead computed a noise ceiling for it. We estimated the noise ceiling as the correlation between a held-out participant’s RDM and the average RDM of the remaining participants.

The two metrics we employed are arguably the two most commonly used for model-brain comparison, and measure distinct things. Regression reveals whether there are linear combinations of model features that can predict brain responses. A model could thus produce high explained variance even if it contained extraneous features that have no correspondence with the brain (because these will get low weight in the linear transform inferred by regression). By comparison, RDMs are computed across all model features, and hence could appear distinct from a brain RDM even if there is a subset of model features that captures the brain’s representational space. Accurate predictions or similar representational geometries also do not necessarily imply that the underlying features are the same in the model and the brain, only that the model features are correlated with brain features across the stimulus set that is used^57, 65^ (typically natural sounds or images). Model-based stimulus generation can help address the latter issue^57^, but ideally require a dedicated neuroscience experiment for each model, which in this context was prohibitive. Although the two metrics we used have limitations, an accurate model of the brain should replicate brain responses according to both metrics, making them a useful starting point for model evaluation.

### Many DNN models outperform traditional models of the auditory cortex

We first assessed the overall accuracy of the brain predictions for each model using regularized regression, aggregating across all voxels in the auditory cortex. For each DNN model, explained variance was measured for each voxel using the single best-predicting stage for that voxel, selected with independent data (see Methods; Voxel response modeling). This approach was motivated by the hypothesis that particular stages of the neural network models might best correspond to particular regions of the cortex. By contrast, the baseline model had a single stage intended to model the auditory cortex (preceded by earlier stages intended to capture cochlear processing), and so we derived predictions from this single “cortical” stage. In each case we then took the median of this explained variance across voxels for a model (averaged across participants).

As shown in Figure 2A, the best-predicting stage of most trained DNN models produced better overall predictions of auditory cortex responses than did the standard SpectroTemporal baseline model^45^ (see Supplementary Figure S2 for predictivity across model stages). This was true for all of the in-house models as well as about half of the external models developed in engineering contexts. However, some models developed in engineering contexts did not produce good predictions, substantially under-predicting the baseline model. The heterogeneous set of external models was intended to test the generality of brain-DNN relations, and sacrificed controlled comparisons between models (because models differed on many dimensions). It is thus difficult to pinpoint the factors that cause some models to produce poor predictions. This finding nonetheless demonstrates that some models that are trained on large amounts of data, and that perform some auditory tasks well, do not accurately predict auditory cortical responses. But the results also show that many models produce better predictions than the classical SpectroTemporal baseline model. As shown in Figure 2A, the results were highly consistent across the two fMRI datasets. In addition, results were fairly consistent for different versions of the in-house models trained from different random seeds (Figure 2B).

**Figure 2.**
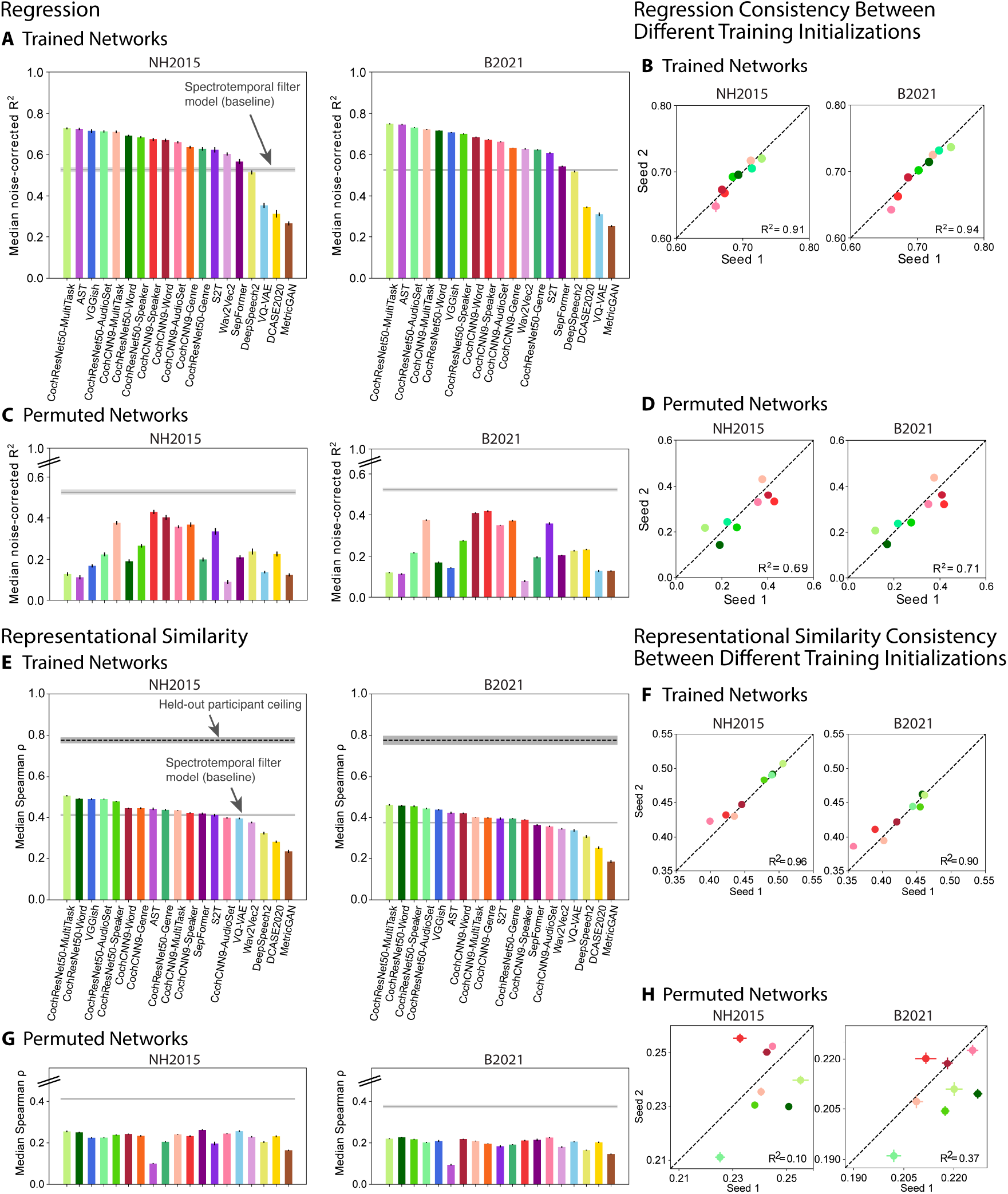
Evaluation of overall model-brain similarity. **(A)** Using regression, explained variance was measured for each voxel and the aggregated median variance explained was obtained for the best-predicting stage for each model, selected using independent data. Grey line shows variance explained by the SpectroTemporal baseline model. Colors indicate the nature of the model architecture: CochCNN9 architectures in shades of red, CochResNet50 architectures in shades of green, Transformer architectures in shades of violet (AST, Wav2vec2, S2T, SepFormer), recurrent architectures in shades of yellow (DCASE2020, DeepSpeech2), other convolutional architectures in shades of blue (VGGish, VQ-VAE), and miscellaneous in brown (MetricGAN). Error bars are within-participant SEM. Error bars are smaller for the B2021 dataset because of the larger number of participants (n=20 vs. n=8). For both datasets, most trained models out-predict the baseline model**. (B)** We trained the in-house models from two different random seeds. The median variance explained for the first and second seed models are plotted on the x- and y-axes, respectively. Each data point represents a model using the same color scheme as in panel A. **(C-D)** Same analysis as in panels A-B but for the control networks with permuted weights. All permuted models produce worse predictions than the baseline. **(E)** Representational similarity between all auditory cortex fMRI responses and the trained computational models. The models and colors are the same as in A. The dashed black line shows the noise ceiling measured by comparing one participant’s RDM with the average of the RDMs from each of the other participants (we plot the noise ceiling rather than noise correcting as in the regression analyses in order to be consistent with what is standard for each analysis). Error bars are within-participant SEM. As in the regression analysis, many of the trained models exhibit RDMs that are more correlated with the human RDM than is the baseline model’s RDM. **(F)** The Spearman correlation between the model and fMRI RDMs for two different seeds of the in-house models. The results for the first and second seeds are plotted on the x- and y-axes, respectively. Each data point represents a model using the same color scheme as in panel E. **(G-H)** Same analysis as in panels E-F but with the control networks with permuted weights. RDMs for all permuted models are less correlated with the human RDM compared to the baseline model’s correlation with the human RDM. Data and code with which to reproduce results are available at https://github.com/gretatuckute/auditory_brain_dnn.

### Brain predictions of DNN models depend critically on task-optimization

To assess whether the improved predictions compared to the SpectroTemporal baseline model could be entirely explained by the DNN architectures, we performed the same analysis with each model’s parameters (e.g., weights, biases) permuted within each model stage (Figure 2C-D). This model manipulation destroyed the parameter structure learned during task-optimization, while preserving the model architecture and the marginal statistics of the model parameters. This was intended as replacement for testing untrained models with randomly initialized weights^31^, the advantage being that it seemed a more conservative test for the external models, for which the initial weight distributions were in some cases unknown.

In all cases, these control models produced worse predictions than the trained models, and in no case did they out-predict the baseline model. This result indicates that task optimization is consistently critical to obtaining good brain predictions. It also provides evidence that the presence of multiple model stages (and selection of the best-predicting stage) is not on its own sufficient to cause a DNN model to out-predict the baseline model. These conclusions are consistent with previously published results^31^ but substantiate them on a much broader set of models and tasks.

### Qualitatively similar conclusions from representational similarity

To ensure that the conclusions from the regression-based analyses were robust to the choice of model-brain similarity metric, we conducted analogous analyses using representational similarity. Analyses of representational similarity gave qualitatively similar results to those with regression. We computed the Spearman correlation between the RDM for all auditory cortex voxels and that for the unit activations of each stage of each model, choosing the model stage that yielded the best match. We used 83 of the sounds to choose the best-matching model stage, and then measured the model-brain RDM correlation for RDMs computed for the remaining 82 sounds. We performed this procedure with 10 different splits of the sounds, averaging the correlation across the 10 splits. This analysis showed that most of the models in our set had RDMs that were more correlated with the human auditory cortex RDM than that of the baseline model (Figure 2E), and the results were consistent across two trained instantiations of the in-house models (Figure 2F). Moreover, the two measures of model-brain similarity (variance explained and correlation of RDMs) were correlated in the trained networks (R^2^=0.75 for NH2015 and R^2^=0.79 for B2021, p<<.001), with models that showed poor matches on one metric tending to show poor matches on the other. The correlations with the human RDM were nonetheless well below the noise ceiling and not much higher than those for the baseline model, indicating that none of the models fully accounts for the fMRI representational similarity. As expected, the RDMs for the permuted models were less similar to that for human auditory cortex, never exceeding the correlation of the baseline model (Figure 2G-H). Overall, these results provide converging evidence for the conclusions of the regression-based analyses.

### Improved predictions of DNN models are most pronounced for pitch, speech, and music-selective responses

To examine the model predictions for specific tuning properties of the auditory cortex, we used a previously derived set of cortical response components. Previous work^50^ found that cortical voxel responses to natural sounds can be explained as a linear combination of six response components (Figure 3A). These six components can be interpreted as capturing the tuning properties of underlying neural populations. Two of these components were well accounted for by audio frequency tuning, and two others were relatively well explained by tuning to spectral and temporal modulation frequencies. One of these latter two components was selective for sounds with salient pitch. The remaining two components were highly selective for speech and music, respectively. The six components had distinct (though overlapping) anatomical distributions, with the components selective for pitch, speech, and music most prominent in different regions of non-primary auditory cortex (Figure 3B; components 4-6, selective for pitch, speech, and music, respectively). These components provide one way to examine whether the improved model predictions seen in Figure 2 are specific to particular aspects of cortical tuning.

**Figure 3.**
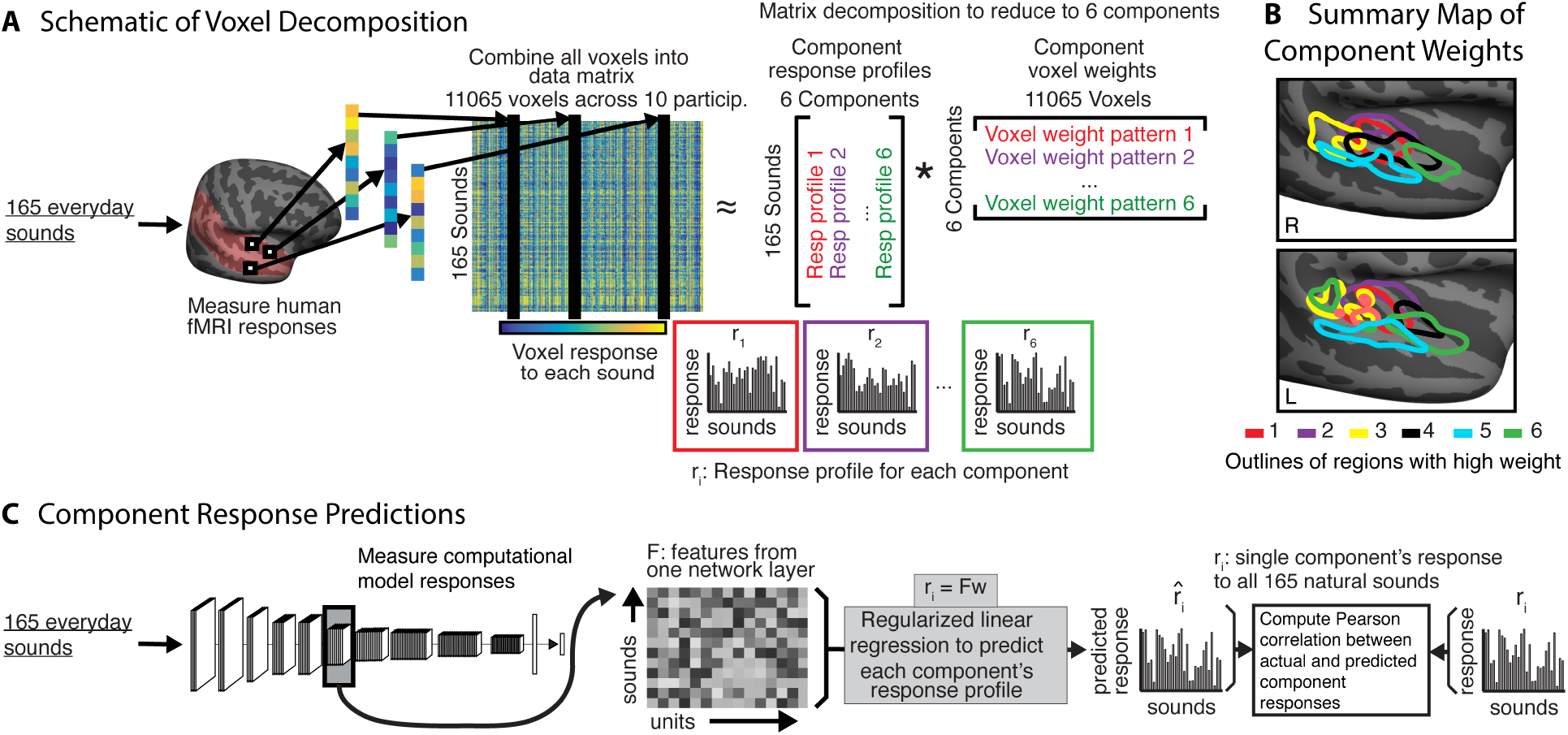
Component decomposition of fMRI responses. **(A)** Voxel component decomposition method. The voxel responses of a set of participants are approximated as a linear combination of a small number of component response profiles. The solution to the resulting matrix factorization problem is constrained to maximize a measure of the non-Gaussianity of the component weights. Voxel responses in auditory cortex to natural sounds are well accounted for by six components. Figure adapted from Norman-Haignere et al.^50^. **(B)** The six components are concentrated in different regions of the auditory cortex. Figure adapted from Norman-Haignere et al.^50^. **(C)** We generated model predictions for each component’s response using the same approach used for voxel responses, in which the model unit responses were combined to best predict the component response, with explained variance measured in held-out sounds (taking the median of the explained variance values obtained across train/test cross-validation splits).

We again used regression to generate model predictions, but this time using the component responses rather than voxel responses (Figure 3C). We fit a linear mapping from the unit activations in a model stage (for a subset of “training” sounds) to the component response, then measured the predictions for left-out “test” sounds, averaging the predictions across test splits. The main difference between the voxel analyses and the component analyses is that we did not noise-correct the estimates of explained component variance. This is because we could not estimate test-retest reliability of the components, as they were derived with all three scans worth of data. We also restricted this analysis to regression-based predictions because representational similarity cannot be measured from single response components.

Figure 4A shows the actual component responses (from the Norman-Haignere et al., 2015 dataset) plotted against the predicted responses for the best-predicting model stage (selected separately for each component) of the multi-task CochResNet50, which gave the best overall voxel response predictions (Figure 2). The model replicates most of the variance in all components (between 61% and 88% of the variance, depending on the component). Given that two of the components are highly selective for particular categories, one might suppose that the good predictions in these cases could be primarily due to predicting higher responses for some categories than others, and the model indeed reproduces the differences in responses to different sound categories (e.g., with high responses to speech in the speech-selective component, and high responses to music in the music-selective component). However, it also replicates some of the response variance within sound categories. For instance, the model predictions explained 51.9% of the variance in responses to speech sounds in the speech-selective component, and 53.5% of the variance in the responses to music sounds in the music-selective component (both of these values are much higher than would be expected by chance; speech: p=.001; music: p<.001). We note that although we could not estimate the reliability of the components in a way that could be used for noise correction, in a previous paper we measured their similarity between different groups of participants, and this was lowest for component 3, followed by component 6^51^. Thus, the differences between components in the overall quality of the model predictions are plausibly related to their reliability.

**Figure 4.**
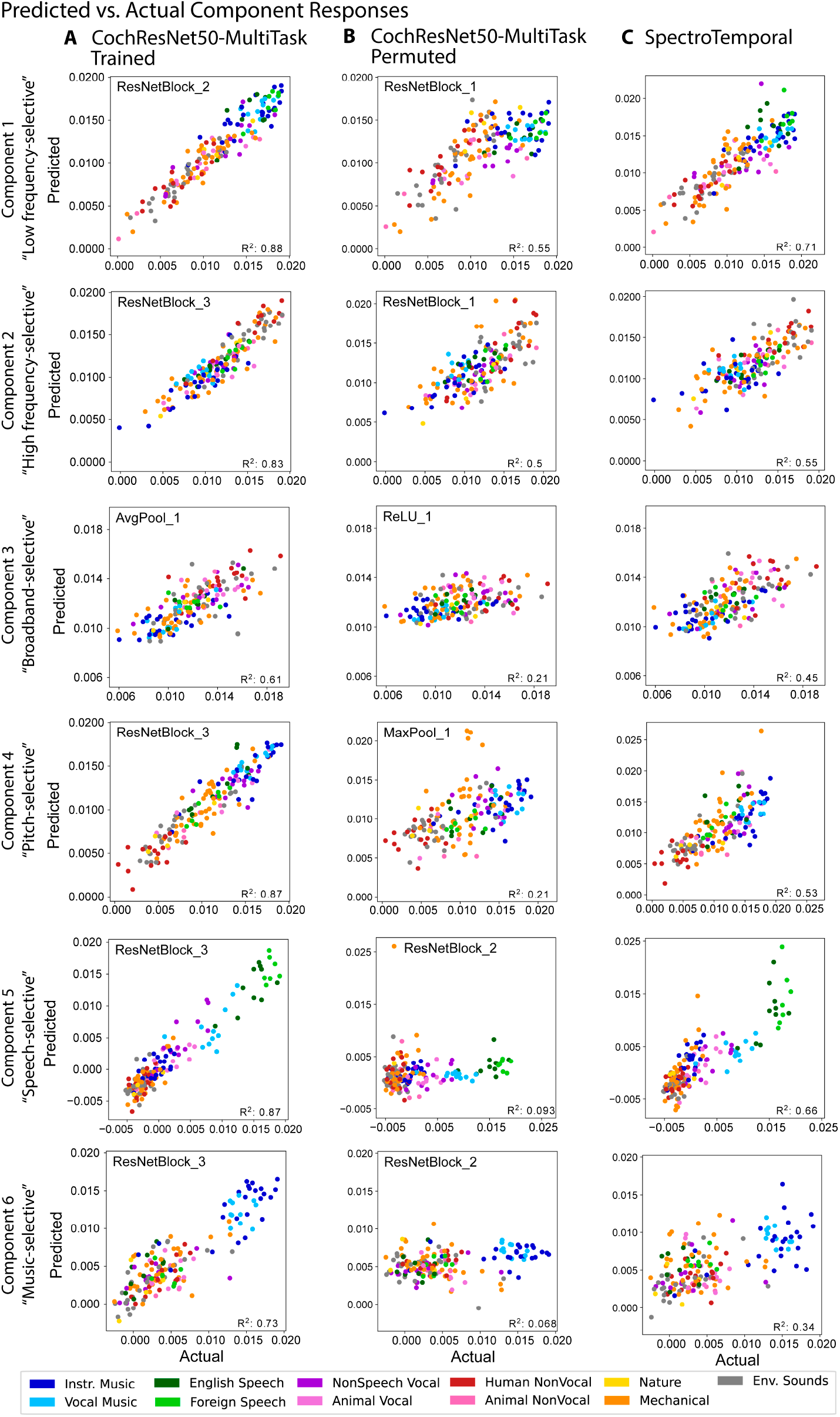
Example model predictions for six components of fMRI responses to natural sounds. **(A)** Predictions of the six components by a trained deep neural network model (CochResNet50-MultiTask). Each data point corresponds to a single natural sound from the set of 165. Data point color denotes the sound’s semantic category. Model predictions were made from the model stage that best predicted a component’s response. The predicted response is the average of the predictions for a sound across the test half of 10 different train-test splits (including each of the splits for which the sound was present in the test half). **(B)** Predictions of the six components by same model used in A but with permuted weights. Predictions are substantially worse than for the trained model, indicating that task optimization is important for obtaining good predictions, especially for components 4-6. **(C)** Predictions of the six components by the SpectroTemporal model. Predictions are substantially worse than for the trained model, particularly for components 4-6. Data and code with which to reproduce results are available at https://github.com/gretatuckute/auditory_brain_dnn.

The component response predictions were much worse for models with permuted weights, as expected given the results of Figure 2 (Figure 4B; results shown for the permuted multi-task CochResNet50; results were similar for other models with permuted weights, though not always as pronounced). The notable exceptions were the first two components, which reflect frequency tuning^50^. This is likely because frequency information is made explicit by a convolutional architecture operating on a cochlear representation, irrespective of the model weights. For comparison we also show the component predictions for the SpectroTemporal baseline model (Figure 4C). These are significantly better than those of the best-predicting stage of the permuted CochResNet50MultiTask model (one-tailed p<<.001; permutation test), but significantly worse than those of the trained CochResNet50MultiTask model for all six components (one-tailed p<<.001; permutation test).

These findings held across most of the neural network models we tested. Most of the trained neural network models produced better predictions than the SpectroTemporal baseline model for most of the components (Figure 5A), with the improvement being specific to the trained models (Figure 5B). However, it is also apparent that the difference between the trained and permuted models is most pronounced for components 4-6 (selective for pitch, speech, and music, respectively; compare Figure 5A to 5B). This result indicates that the improved predictions for task-optimized models are most pronounced for higher-order tuning properties of auditory cortex.

**Figure 5.**
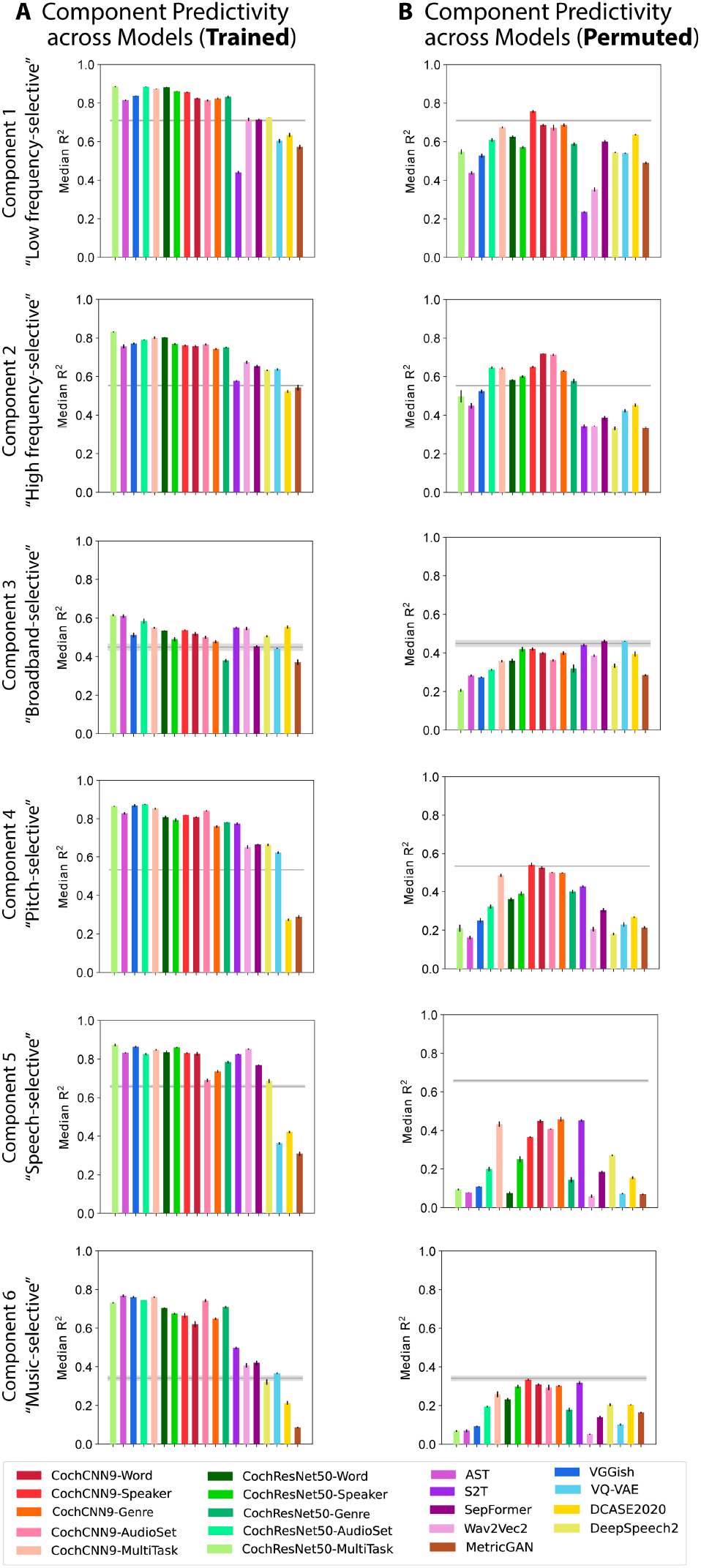
Summary of model predictions of fMRI response components. **(A)** Component response variance explained by each of the trained models. Model ordering is the same as that in Figure 2A for ease of comparison. Variance explained was obtained from the best-predicting stage of each model for each component, selected using independent data. Error bars are SEM over iterations of the model stage selection procedure (see Methods; Component modeling). See Supplementary Figure S3 for a comparison of results for models trained with different random seeds (results were overall similar for different seeds). **(B)** Component response variation explained by each of the permuted models. The trained models (both in-house and external), but not the permuted models, tend to out-predict the SpectroTemporal baseline for all components, but the effect is most pronounced for components 4-6. Data and code with which to reproduce results are available at https://github.com/gretatuckute/auditory_brain_dnn.

### Many DNN models exhibit model-stage-brain-region correspondence with auditory cortex

One of the most intriguing findings from the neuroscience literature on deep neural network models is that the models often exhibit some degree of correspondence with the hierarchical organization of sensory systems^13–16, 31, 66^, with particular model stages providing the best matches to responses in particular brain regions. To explore the generality of this correspondence for audio-trained models, we first examined the best-predicting model stage for each voxel of each participant in the two fMRI datasets, separately for each model. We used regression-based predictions for this analysis as it was based on single voxel responses.

We first plotted the best-predicting stage as a surface map displayed on an inflated brain. The best-predicting model stage for each voxel was expressed as a number between 0 and 1, and we plot the median of this value across participants. In both datasets and for most models, earlier model stages tended to produce the best predictions of primary auditory cortex while deeper model stages produced better predictions of non-primary auditory cortex. We show these maps for the eight best-predicting models in Figure 6A, and provide them for all remaining models in Supplementary Figure S4. There was some variation from model to model, both in the relative stages that yield the best predictions, and in the detailed anatomical layout of the resulting maps, but the differences between primary and non-primary auditory cortex were fairly consistent across models. The stage-region correspondence was specific to the trained models; the models with permuted weights produce relatively uniform maps (Supplementary Figure S5).

**Figure 6.**
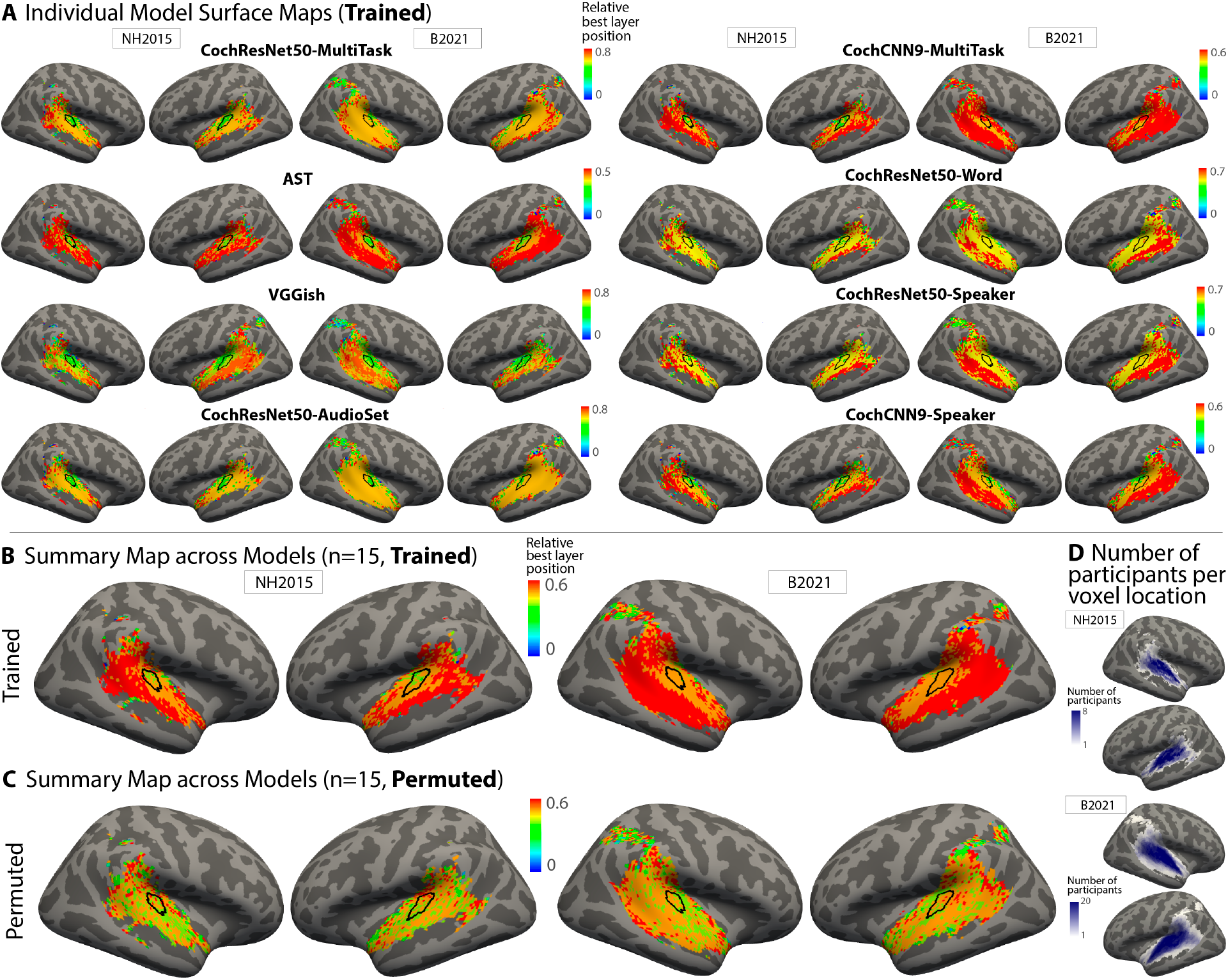
Surface maps of best-predicting model stage. **(A)** To investigate correspondence between model stages and brain regions, we plot the model stage that best predicts each voxel as a surface map (FsAverage) (median best stage across participants). We assigned each model stage a position index between 0 and 1 (using minmax normalization such that the first stage is assigned a value of 0 and the last stage a value of 1). We show this map for the eight best-predicting models as evaluated by the median noise-corrected R^2^ plotted in Figure 2A (see Supplementary Figure S4 for maps from other models). The color scale limits were set to extend from 0 to the stage beyond the most common best stage (across voxels). We found that setting the limits in this way made the variation across voxels in the best stage visible by not wasting dynamic range on the deep model stages, which were almost never the best-predicting stage. Because the relative position of the best-predicting stage varied across models, the color bar scaling varies across models. For both datasets, middle stages best predict primary auditory cortex, while deep stages best predict non-primary cortex. We note that the B2021 dataset contained voxel responses in parietal cortex, some of which passed the reliability screen. We have plotted a best-predicting stage for these voxels in these maps for consistency with voxel inclusion criteria in the original publication^51^, but note that these voxels only passed the reliability screen in a few participants (see panel D), and that the variance explained for these voxels was low, such that the best-predicting stage is not very meaningful. **(B)** Best-stage map averaged across all models that produced better predictions than the baseline SpectroTemporal model. The map plots the median value across models, and thus is composed of discrete color values. The thin black outline plots the borders of an anatomical ROI corresponding to primary auditory cortex. **(C)** Best-stage map for the same models as in (B), but with permuted weights. **(D)** Maps showing the number of participants per voxel location on the FsAverage surface for both datasets (1-8 participants for NH2015; 1-20 participants for B2021). Darker colors denote a larger number of participants per voxel. Because we only analyzed voxels that passed a reliability threshold, some locations only passed the threshold in a few participants. Note also that the regions that were scanned were not identical in the two datasets. Data and code with which to reproduce results are available at https://github.com/gretatuckute/auditory_brain_dnn.

To summarize these maps across models, we computed the median best-stage for each voxel across all 15 models that produced better overall predictions compared to the baseline model (Figure 2A). The resulting map provides a coarse measure of the central tendency of the individual model maps (at the cost of obscuring the variation that is evident across models). If there was no consistency across the maps for different models, this map should be uniform. Instead, the best-stage summary map (Figure 6B) shows a clear gradient, with voxels in and around primary auditory cortex (black outline) best predicted by earlier stages than voxels beyond primary auditory cortex. This correspondence is lost when the weights are permuted, contrary to what would be expected if the model architecture was primarily responsible for the correspondence (Figure 6C).

To quantify the trends that were subjectively evident in the surface maps, we computed the average best stage within four anatomical regions of interest (ROIs): one for primary auditory cortex, along with three ROIs for posterior, lateral, and anterior non-primary auditory cortex. These ROIs were combinations of subsets of ROIs in the Glasser et al. parcellation^67^ (Figure 7A). The ROIs were taken directly from a previous publication^51^, where they were intended to capture the auditory cortical regions exhibiting reliable responses to natural sounds, and were not adapted in any way to the present analysis. We visualized the results of this analysis by plotting the average best stage for the primary ROI vs. that of each of the non-primary ROIs, expressing the stage’s position within each model as a number between 0 and 1 (Figure 7B). In each case, nearly all models lie above the diagonal (Figure 7C), indicating that all three regions of non-primary auditory cortex are consistently better predicted by deeper model stages compared to primary auditory cortex, irrespective of the model. This result was statistically significant in each case (Wilcoxon signed rank test: two-tailed p<.005 for all six comparisons; two datasets x three non-primary ROIs). By comparison, there was no clear evidence for differences between the three non-primary ROIs (two-tailed Wilcoxon signed rank test: after Bonferroni correction for multiple comparisons, none of the six comparisons reached statistical significance at the p<.05 level; two datasets x three comparisons). See Supplementary Figure S6 for the variance explained by each model stage for each model for the four ROIs.

**Figure 7.**
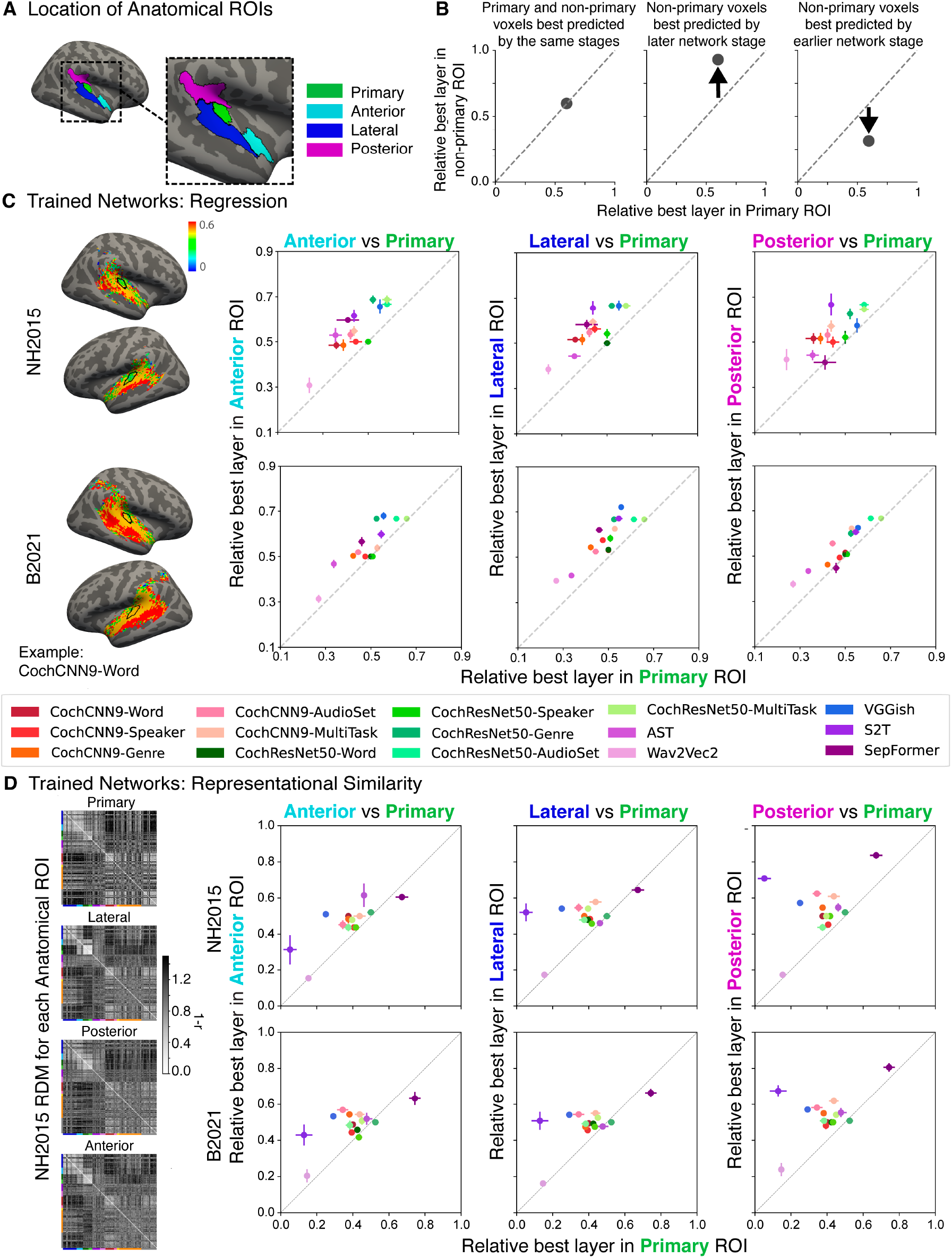
Nearly all DNN models exhibit stage-region correspondence. **(A)** Anatomical ROIs for analysis. ROIs were reproduced from a previous study^51^, in which they were derived by pooling ROIs from the Glasser anatomical parcellation^67^. **(B)** To summarize the model-stage-brain-region correspondence across models, we obtained the median best-predicting stage for each model within the four anatomical ROIs from A: primary auditory cortex (x axis in each plot in C and D) and anterior, lateral, and posterior non-primary regions (y axes in C and D). **(C)** We performed the analysis on each of the two fMRI datasets, including each model that out-predicted the baseline model in Figure 2 (n=15 models). Each data point corresponds to a model, with the same color correspondence as in Figure 2. Error bars are within-participant SEM. The non-primary ROIs are consistently best-predicted by later stages than the primary ROI. **(D)** Same analysis as C but with the best-matching model stage determined by correlations between the model and ROI representational dissimilarity matrices. RDMs for each anatomical ROI (left) are grouped by sound category, indicated by colors on the left and bottom edges of each RDM (same color-category correspondence as in Figure 4). Larger-scale fMRI RDMs for each ROI including the name of each sound is provided in Supplementary Figure S1. Data and code with which to reproduce results are available at https://github.com/gretatuckute/auditory_brain_dnn.

To confirm that these results were not merely the result of the deep neural network architectural structure (for instance, with pooling operations tending to produce larger receptive fields at deeper stages compared to earlier stages), we performed the same analysis on the models with permuted weights. In this case the results showed no evidence for a mapping between model stages and brain regions (Supplementary Figure S7A; no significant difference between primary and non-primary ROIs in any of the six cases; Wilcoxon signed rank tests, two-tailed p>0.16 in all cases). This result is consistent with the surface maps (Figure 6C and Supplementary Figure S5), which tended to be fairly uniform.

We repeated the ROI analysis using representational similarity to determine the best-matching model stage for each ROI, and obtained similar results. The model stages with representational geometry most similar to that of non-primary ROIs were again situated later than the model stage most similar to the primary ROI, in both datasets (Figure 7D; Wilcoxon signed rank test: two-tailed p<.007 for all six comparisons; two datasets x three non-primary ROIs). The model stages that provided the best match to each ROI according to each of the two metrics (regression and RSA) were correlated (R^2^=0.21 for NH2015 and R^2^=0.21 for B2021, measured across the 60 best stage values from 15 trained models for the four ROIs of interest, p<.0005 in both cases). This correlation is highly statistically significant but is nonetheless well below the maximum it could be given the reliability of the best stages (conservatively estimated as the correlation of the best stage between the two fMRI datasets; R^2^=0.87 for regression and R^2^=0.94 for representational similarity). This result suggests that the two metrics capture different aspects of model-brain similarity and that they do not fully align for the models we have at present, even though the general trend for deeper stages to better predict non-primary responses is present in both cases.

Overall, these results are consistent with the stage-region correspondence findings of Kell et al. (2018), but show that they apply fairly generally to a wide range of DNN models, that they replicate across different brain datasets, and are generally consistent across different analysis methods. The results suggest that the different representations learned by early and late stages of DNN models map onto differences between primary and non-primary auditory cortex in a way that is qualitatively consistent across a diverse set of models. This finding provides support for the idea that primary and non-primary human auditory cortex instantiate distinct types of representations that resemble earlier and later stages of a computational hierarchy. However, the specific stages that best align with particular cortical regions vary across models, and depend on the metric used to evaluate alignment. Together with the finding that all model predictions are well below the maximum attainable value given the measurement noise (Figure 2), these results indicate that none of the tested models fully account for the representations in human auditory cortex.

### Presence of noise in training data modulates model predictions

We found in our initial analysis that many models produced good predictions of auditory cortical brain responses, in that they out-predicted the SpectroTemporal baseline model (Figure 2). But some models gave better predictions than others, raising the question of what causes differences in model predictions. To investigate this question, we analyzed the effect of a) the training data and b) the training task the model was optimized for, using the in-house models which consisted of the same two architectures trained on different tasks.

Out of the many manipulations of training data that one could in principle test, the presence of background noise was of particular theoretical interest. Background noise is ubiquitous in real-world auditory environments, and the auditory system is relatively robust to its presence^31, 68–73, 74, 75^, suggesting that it might be important in shaping auditory representations. For this reason, the previous models in Kell et al.^31^, as well as the in-house model extensions shown in Figure 2, were all trained to recognize words and speakers in noise (on the grounds that this is more ecologically valid than training exclusively on “clean” speech). We previously found in the domains of pitch perception^32^ and sound localization^33^ that optimizing models for natural acoustic conditions (including background noise among other factors) was critical to reproducing the behavioral characteristics of human perception, but it was not clear whether model-brain similarity metrics would be analogously affected. To test whether the presence of noise in training influences model-brain similarity, we retrained both in-house model architectures on the word and speaker recognition tasks using just the speech stimuli from the Word-Speaker-Noise dataset (without added noise). We then repeated the analyses of Figures 2 and 5.

As shown in Figure 8, models trained exclusively on clean speech produced significantly worse overall brain predictions compared to those trained on speech in noise. This result held for both the word and speaker recognition tasks, and for both model architectures (regression: p<<.001 via one-tailed bootstrap test for each of the eight comparisons, two datasets x four comparisons; representational similarity: p<<.001 in each case for same comparisons). The result was not specific to speech-selective brain responses, as the training noise boost was seen for each of the pitch-selective, speech-selective, and music-selective response components (Supplementary Figure S8). This training data manipulation is obviously one of many that are in principle possible, and does not provide an exhaustive examination of the effect of training data, but the results are consistent with the notion that optimizing models for sensory signals like those for which biological sensory systems are plausibly optimized increases the match to brain data. They are also consistent with findings that machine speech recognition systems, which are typically trained on clean speech (usually because there are separate systems to handle denoising), do not always reproduce characteristics of human speech perception^76, 77^.

**Figure 8.**
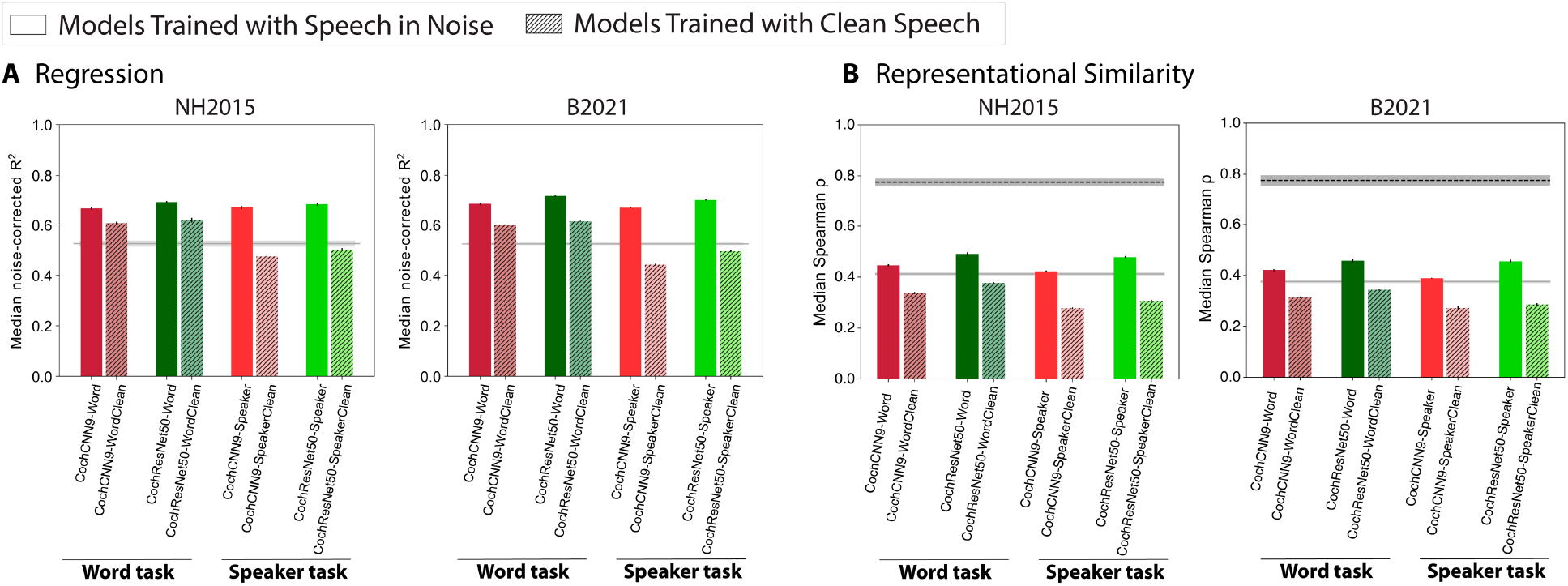
Model predictions of brain responses are better for models trained in background noise. **(A)** Effect of noise in training on model-brain similarity assessed via regression. Using regression, explained variance was measured for each voxel and the aggregated median variance explained was obtained for the best-predicting stage for each model, selected using independent data. Grey line shows variance explained by the SpectroTemporal baseline model. Colors indicate the nature of the model architecture with CochCNN9 architectures in shades of red, and CochResNet50 architectures in shades of green. Models trained in the presence of background noise are shown in the same color scheme as in Figure 2; models trained with clean speech are shown with hashing. Error bars are within-participant SEM. For both datasets, the models trained in the presence of background noise exhibit higher model-brain similarity than the models trained without background noise. **(B)** Effect of noise in training on model-brain representational similarity. Same conventions as A, except that the dashed black line shows the noise ceiling measured by comparing one participant’s RDM with the average of the RDMs from each of the other participants. Error bars are within-participant SEM. Data and code with which to reproduce results are available at https://github.com/gretatuckute/auditory_brain_dnn.

### Training task modulates model predictions

To assess the effect of the training task a model was optimized for, we analyzed the brain predictions of the in-house models, which consisted of the same two architectures trained on different tasks. The results shown in Figure 2 indicate that some of our in-house tasks produced better overall predictions than others, and that the best overall model as evaluated with either metric (regression or RDM similarity) was that trained on three of the tasks (the CochResNet50-MultiTask).

To gain insight into the source of these effects, we examined the in-house model predictions for the six components of auditory cortical responses (Figure 3) that vary across brain regions. The components seemed a logical choice for an analysis of the effect of task on model predictions because they isolate distinct cortical tuning properties. We focused on the pitch-selective, speech-selective, and music-selective components, because these showed the largest effects of model training (components 4-6, Figure 4&5), and because the tasks that we trained on seemed a priori most likely to influence representations of these types of sounds. This analysis was necessarily restricted to the regression-based model predictions because RDMs are not defined for any single component’s response.

A priori it was not clear what to expect. The representations learned by neural networks are a function both of the training stimuli and the task they are optimized for^32, 33^, and in principle either (or both) could be critical to reproducing the tuning found in the brain. For instance, it seemed plausible that speech- and music-selectivity might only become strongly evident in systems that must perform speech- and music-related tasks. However, given the distinct acoustic properties of speech, music and pitch, it also seemed plausible that they might naturally segregate within a distributed neural representation simply from generic representational constraints that might occur for any task, such as the need to represent sounds efficiently^78–80^ (here imposed by the finite number of units in each model stage). Our in-house tasks allowed us to distinguish these possibilities, because the training stimuli were held constant (for three of the tasks, and for the multi-task model; the genre task involved a distinct training set), with the only difference being the labels that were used to compute the training loss. Thus, any differences in predictions between these models reflect changes in the representation due to behavioral constraints rather than the training stimuli.

We note that the AudioSet task consists of classifying sounds within YouTube soundtracks, and the sounds and associated labels are diverse. In particular it includes many labels associated with music – both musical genres, and instruments (67 and 78 classes, respectively, out of 516 total). By comparison, our musical genre classification task contained exclusively genre labels, but only 41 of them. It thus seemed plausible that the AudioSet task might produce music- and pitch-related representations.

Comparisons of the variance explained in each component revealed interpretable effects of the training task (Figure 9). The pitch-selective component was best predicted by the models trained on AudioSet (R^2^ was higher for AudioSet-trained model than for the word-, speaker- or genre-trained models in both the CochCNN9 and CochResNet50 architectures, one-tailed p<.0005 for all six comparisons, permutation test). The speech-selective component was best predicted by the models trained on speech tasks. This was true both for the word recognition task (R^2^ higher for the word-trained model than for the genre or AudioSet-trained models for both architectures, one-tailed p<.0005 for three out of four comparisons; p=0.19 for CochResNet50-Word vs. AudioSet) and for the speaker recognition task (one-tailed p<.005 for all four comparisons). Finally, the music-selective component was best predicted by the models trained on AudioSet (R^2^ higher for the AudioSet-trained model than for the word-, speaker- or genre-trained models for both architectures, p<.0005 for all six comparisons), consistent with the presence of music-related classes in this task. We note also that the component was less well predicted by the models trained to classify musical genre. This latter result may indicate that the genre dataset/task does not fully tap into the features of music that drive cortical responses. For instance, some genres could be distinguished by the presence or absence of speech, which may not influence the music component’s response^50, 51^ (but which could enable the representations learned from the genre task to predict the speech component). Note that the absolute variance explained in the different components cannot be straightforwardly compared, because the values are not noise-corrected (unlike the values for the voxel responses).

**Figure 9.**
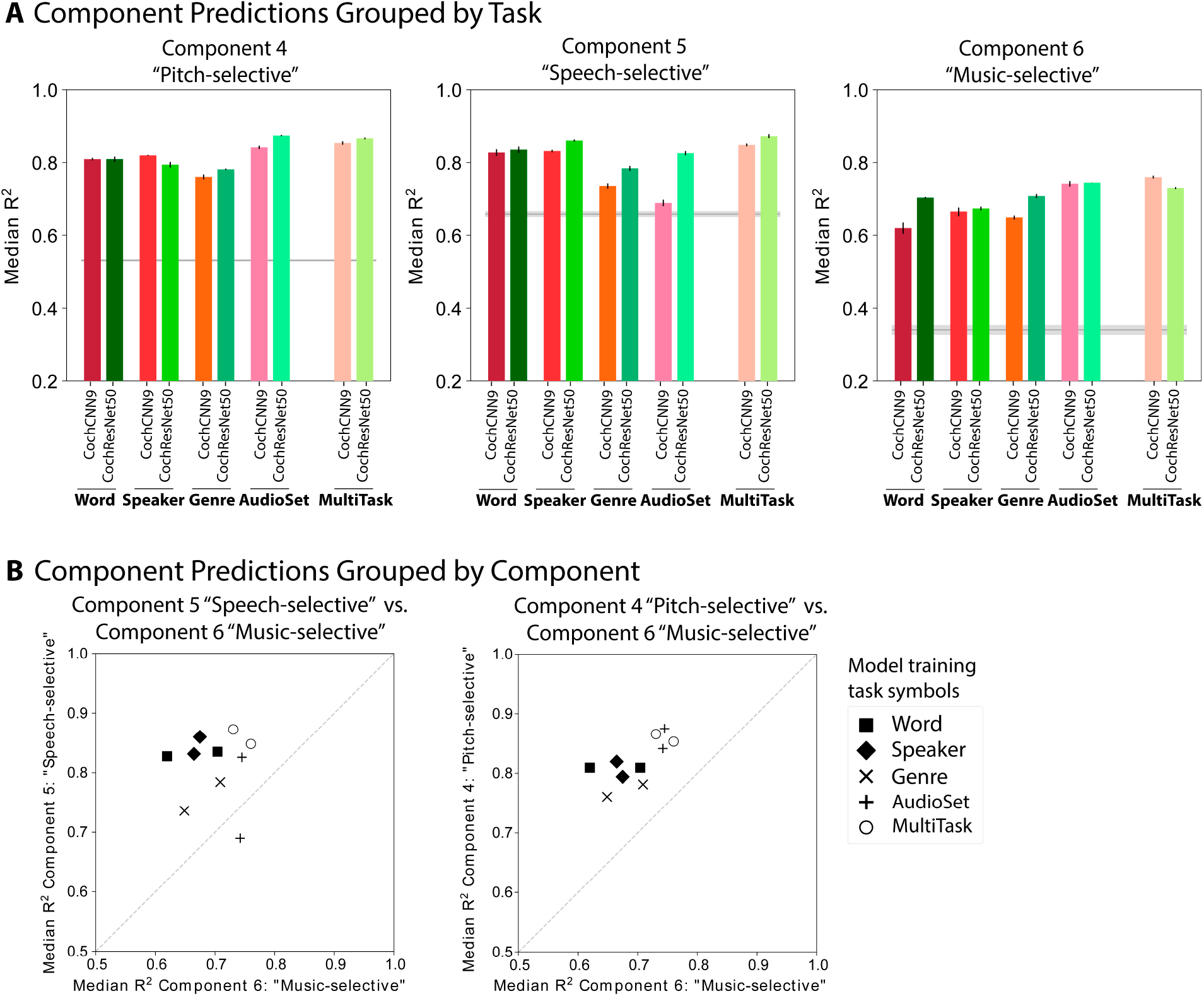
Training task modulates model predictions. **(A)** Component response variance explained by each of the trained in-house models. Predictions are shown for components 4-6 (pitch-selective, speech-selective, and music-selective, respectively). The in-house models were trained separately on each of four tasks as well as on three of the tasks simultaneously, using two different architectures. Explained variance was measured for the best-predicting stage of each model for each component selected using independent data. Error bars are SEM over iterations of the model stage selection procedure (see Methods; Component modeling). Gray line plots the variance explained by the SpectroTemporal baseline model. **(B)** Scatter plots of in-house model predictions for pairs of components. The upper panel shows the variance explained for component 5 (speech-selective) vs. component 6 (music-selective), and the lower panels shows component 6 (music-selective) vs. component 4 (pitch-selective). Symbols denote the training task. In the left panel, the four models trained on speech-related tasks are furthest from the diagonal, indicating good predictions of speech-selective tuning at the expense of those for music-selective tuning. In the right panel, the models trained on AudioSet are set apart from the others in their predictions of both the pitch-selective and music-selective components. Error bars are smaller than the symbol width (and are provided in panel A) and so are omitted for clarity. Data and code with which to reproduce results are available at https://github.com/gretatuckute/auditory_brain_dnn.

The differences between tasks were most evident in scatter plots of the variance explained for pairs of components (Figure 9B). For instance, the speech-trained models are furthest from the diagonal when the variance explained in the speech and music components are compared. And the AudioSet-trained models, along with the multi-task models, are well separated from the other models when the pitch- and music-selective components are compared. Given that these models were all trained on the same sounds, the differences in their ability to replicate human cortical tuning for pitch, music and speech suggests that these tuning properties emerge in the models from the demands of supporting of specific behaviors. The results do not exclude the possibility that these tuning properties could also emerge through some form of unsupervised learning, or from some other combination of tasks. But they nonetheless provide a demonstration that the distinct forms of tuning in the auditory cortex could be a consequence of specialization for domain-specific auditory abilities.

We found that in each component and architecture, the multi-task models predicted component responses about as well as the best single-task model. It was not obvious a priori that a model trained on multiple tasks would capture the benefits of each single-task model – one might alternatively suppose that the demands of supporting multiple tasks with a single representation would muddy the ability to predict domain-specific brain responses. Indeed, the multi-task models achieved slightly lower task performance than the single-task models on each of the tasks (see Methods; Training CochResNet50 and CochCNN9 models – Word, Speaker, and AudioSet tasks). This result is consistent with the results of Kell et al. that dual-task performance was impaired in models that were forced to share representations across tasks^31^. However, this effect evidently did not prevent the multi-task model representations from capturing speech- and music-specific response properties. This result indicates that multi-task training is a promising path toward better models of the brain, in that the resulting models appear to combine the advantages of individual tasks.

### Representation dimensionality correlates with model predictivity but does not explain it

Although the task manipulation showed a benefit of multiple tasks in our in-house models, the task alone does not obviously explain the large variance across external models in the measures of model-brain similarity that we used. Motivated by recent findings that the dimensionality of a model’s representation tends to correlate with regression-based brain predictions of ventral visual cortex^81^, we examined whether a model’s effective dimensionality could account for some of the differences we observed between models (Supplementary Figure S9).

The effective dimensionality is intended to summarize the number of dimensions over which a model’s activations vary for a stimulus set, and is estimated from the eigenvalues of the covariance matrix of the model activations to a stimulus set (see Methods; Effective dimensionality). Effective dimensionality cannot be greater than the minimum value of either the number of stimuli in the stimulus set or the model’s ambient dimensionality (i.e., the number of unit activations), but is typically lower than both of these because the activations of different units in a model can be correlated. Effective dimensionality must limit predictivity when a model’s dimensionality is lower than the dimensionality of the underlying neural response, because a low dimensional model response could not account for all of the variance in a high dimensional brain response.

We measured effective dimensionality for each stage of each evaluated model (Supplementary Figure S9). We pre-processed the model activations to match the pre-processing used for the model-brain comparisons. The effective dimensionality for model stages ranged from ∼1 to ∼65 for our stimulus set (using the regression analysis pre-processing). By comparison, the effective dimensionality of the fMRI responses was 8.75 (for NH2015) and 5.32 (for B2021). Effective dimensionality tended to be higher in trained than in permuted models, and tended to increase from one model stage to the next in trained models. The effective dimensionality of a model stage was modestly correlated with the stage’s explained variance (R^2^=0.19 and 0.20 for NH2015 and B2021, respectively; Supplementary Figure S9, panel Aii), and with the model-brain RDM similarity (R^2^=0.15 and 0.18 for NH2015 and B2021, respectively; Supplementary Figure S9, panel Bii). However, this correlation was much lower than the reliability of the explained variance measure (R^2^=0.98, measured across the two fMRI datasets for trained networks; Supplementary Figure S9, panel Ai), and the reliability of the model-brain RDM similarity (R^2^=0.96; Supplementary Figure S9, panel Bi). Effective dimensionality thus does not explain the majority of the variance across models – there was wide variation in the dimensionality of models with good predictivity, and also wide variation in predictivity of models with similar dimensionality.

Intuitively, dimensionality could be viewed as a confound for regression-based brain predictions. High-dimensional model representations might be more likely to produce better regression scores by chance, on the grounds that the regression can pick out a small number of dimensions that approximate the function underlying the brain response, while ignoring other dimensions that are not brain-like. But because the RDM is a function of all of a representation’s dimensions, it is not obvious why high-dimensionality on its own should lead to higher RDM similarity. The comparable relationship between RDM similarity and dimensionality thus helps to rule out dimensionality as a confound in the regression analyses. In addition, both relationships were quite modest. Overall, the results show that there is a weak relationship between dimensionality and model-brain similarity, but that it cannot explain most of the variation we saw across models.

## Discussion

We examined similarities between representations learned by contemporary deep neural network models and those in the human auditory cortex, using regression and representational similarity analyses to compare model and brain responses to natural sounds. We used two different brain datasets to evaluate a large set of models trained to perform audio tasks. Most of the models we evaluated produced more accurate brain predictions than a standard spectrotemporal filter model of the auditory cortex^45^. Predictions were consistently much worse for models with permuted weights, indicating a dependence on task-optimized features. The improvement in predictions with model optimization was particularly pronounced for cortical responses in non-primary auditory cortex selective for pitch, speech, or music. Predictions were worse for models trained without background noise. We also observed task-specific prediction improvements for particular brain responses, e.g., with speech tasks producing the best predictions of speech-selective brain responses. Accordingly, the best overall predictions (aggregating across all voxels) were obtained with models trained on multiple tasks. We also found that most models exhibited some degree of correspondence with the presumptive auditory cortical hierarchy, with primary auditory voxels being best predicted by model stages that were consistently earlier than the best-predicting model stages for non-primary voxels. The training-dependent model-brain similarity and model-stage-brain-region correspondence was evident both with regression and representational similarity analyses. The results indicate that more often than not, deep neural network models optimized for audio tasks learn representations that capture aspects of human auditory cortical responses and organization.

Our general strategy was to test as many models as we could, and the model set included every audio model with an implementation in PyTorch that was publicly available at the time of our experiments. The motivation for this “kitchen sink” approach was to provide a strong test of the generality of brain-DNN correspondence. The cost is that the resulting model comparisons were uncontrolled – the external models varied in architecture, training task, and training data, such that there is no way to attribute differences between model results to any one of these variables. To better distinguish the role of the training data and task, we complemented the external models with a set of models built in our lab that enabled a controlled manipulation of task, and some manipulations of the training data. These models had identical architectures, and for three of the tasks had the same training data, being distinguished only by which of three types of labels the model was asked to predict.

### Insights into the auditory system

What do our results reveal about the auditory system? The main immediate biological contribution lies in providing further evidence and context for functional differentiation between regions of human auditory cortex. Discussions of auditory cortical functional organization commonly revolve around two proposed principles. The first is that the cortex is organized hierarchically into a sequence of stages corresponding to cortical regions^31, 82–84^. Much of the evidence for hierarchy is associated with speech processing, in that speech-specific responses only emerge outside of primary cortical areas^50, 59, 85–90, 91^. Other evidence for hierarchical organization comes from analyses of responses to natural sounds, which show selective responses to music and song in non-primary auditory cortex^50, 51, 92^. These non-primary responses occur with longer latencies and longer integration windows^93^ than primary cortical responses. In addition, stimuli that are matched in audio and modulation frequency content to natural sounds drive responses in primary, but not non-primary, auditory cortex^57^. Non-primary areas also show greater invariance to real-world background noise^74^. The present results provide a distinct additional type of evidence for a broad distinction between the computational description of primary and non-primary auditory cortex, with primary and non-primary voxels being consistently best-predicted by earlier and later stages of hierarchical models, respectively. We note that these results do not speak to the anatomical connections between regions, only to their stimulus selectivity and correspondence to hierarchical computational models. The present results in particular do not necessarily imply that the observed regional differences reflect strictly sequential stages of processing^94^. But they do show that the qualitative relationship to earlier and later model stages is fairly consistent across datasets and models.

The second commonly articulated principle of functional organization is that of domain specificity – the idea that different regions are specialized for different auditory functions. Previous evidence for this idea comes from findings that selectivity for particular stimulus attributes is localized to distinct regions of auditory cortex. In particular, speech selectivity is typically found to be localized to the superior temporal gyrus^50, 59, 85–90, 91^, music-selective responses are localized anterior and posterior from primary auditory cortex^50, 51, 92, 95, 96^, and location-specific responses to the planum temporale^97–101^. The present results provide additional evidence for domain-specific responses, in that particular tasks produced model representations that best predicted particular response components. This was true even though the models in question were trained on identical sound sets. The results indicate that the way sound is used to perform tasks can shape representations in ways that cannot be entirely explained by the distribution of sound features a system is optimized for. These results were not a foregone conclusion: a priori it seemed possible that the particular classification tasks we used would not be sufficient to differentially account for auditory cortical tuning. It also seemed possible that domain-specific responses might arise irrespective of the training task, based on the acoustically distinct properties of speech, music, and pitch. The results lend plausibility to the idea that the response selectivity found in the human auditory cortex (to pitch, music, and speech) could arise from optimization for specific tasks, though it does not prove this possibility (because it remains possible that similar tuning properties could emerge from the right type of unsupervised learning).

### How to build a model of auditory cortex?

What do our results reveal about how to build a good model of human auditory cortex? First, they provide broad additional support for the idea that training a hierarchical model to perform tasks on natural audio signals produces representations that exhibit some alignment with the cortex as measured by two commonly used metrics (better alignment than was obtainable by previous generations of models). The fact that many models produce relatively good predictions suggests that these models contain audio features that typically align to some extent with brain representations, at least for the fMRI measurements we are working with, and in the sense of producing responses that are correlated for natural sounds. And the fact that results were consistently worse for models with permuted weights suggests that training is critical to obtaining these features. Second, some models built for engineering purposes produce poor brain predictions. Although the heterogeneity of the models limits our ability to diagnose the factors that underlie these model-brain discrepancies, the result suggests that we should not expect every DNN model to produce strong alignment with the brain. Third, background noise in the training data consistently improved the predictions of models trained on speech tasks. The improvement held for speech-selective responses in addition to responses not specifically related to speech. Thus, even the representation of speech seems to be made more brain-like by training in noise. Fourth, multiple tasks seem to improve predictions. The results suggest that particular tasks produce representations that align well with particular brain responses, such that a model trained on multiple tasks gets the best of all worlds (Figure 9). We note also that the two external models that produced model-brain similarity on par with the in-house models were trained on one of the in-house model tasks (AudioSet), further consistent with the idea that the task is important. Fifth, models with higher-dimensional representations are somewhat more likely to produce good matches with the brain. At present it is not clear what drives this effect, but there was a modest but consistent effect evident with both metrics we used.

A majority of the models we tested produced overall predictivity between 60% and 75% of the explainable variance (for regression-based predictions). These values were well above that of the SpectroTemporal baseline model but well below the noise ceiling. Moreover, the RDM similarity was far below the noise ceiling for all models. This result thus indicates that all current models are inadequate as complete descriptions of auditory fMRI responses. Our results provide some suggestions for how to improve the models, with the main avenue being training on more realistic data and on a more diverse set of tasks, but it remains to be seen whether incremental extensions will be enough to bridge the gap that is evident in our results.

We note that none of the DNN models we tested were designed or tuned in any way to be able to predict brain responses. One of the in-house models was the result of an architecture search, but that search was constrained only to achieve good task performance. Thus, the models were optimized only to be able to carry out particular auditory functions. The advent of publicly available brain/behavior benchmarks^102^ raises the possibility that models could be “hacked” to score well on model-brain comparisons, but such benchmarks are not yet in widespread use in audition, and played no role in our model development. By contrast, the baseline SpectroTemporal filter model was explicitly designed to replicate auditory cortical tuning seen experimentally in spectrotemporal receptive fields^45^. For instance, the filters are logarithmically spaced, consistent with neurophysiological^103^ and psychophysical^104, 105^ observations, and tuned to both spectral and temporal modulation. It is thus expected that the baseline model would be able to predict cortical responses to some extent, particularly in primary auditory cortex^56, 57^, and indeed its predictions were much better than those of the permuted models. The fact that task-optimized models tend to produce better matches to the cortex than the baseline model thus seems nontrivial.

Another difference between the tested DNN models and the baseline SpectroTemporal model is the number of stages: the DNN models have multiple stages, while the baseline model has a single stage intended to model the auditory cortex. The multiple stages in DNN models require choosing which model stages contribute to model predictions. In principle, all stages of a DNN model could be used concurrently to model the brain response of interest. Instead, our model comparisons used the best-predicting model stage (selected using data distinct from that used to measure the explained variance or RDM similarity). Although this allows an additional hyperparameter when fitting multi-stage models, this analysis choice could be argued to disadvantage multi-stage models – if there was no correspondence between model stages and brain regions, or if the correspondence was sufficiently poor, better predictions might result by using features combined across multiple stages. We also found that the DNN models with permuted weights in all cases produced worse matches to the cortex than the single-stage baseline model, providing further evidence that the analysis procedure does not intrinsically favor multi-stage models. These observations again suggest that the better-then-baseline matches of task-optimized DNN models to the cortex are nontrivial.

The poor performance of some of the models built for engineering purposes suggests a cautionary note. Training neural network models at scale requires substantial compute resources and expertise, and it is thus tempting to obtain models that have been developed in industry labs for other purposes and attempt to use them as models of the brain^41, 43, 106^. Our results suggest that one should not assume that such a model will necessarily produce good brain predictions. In particular, the three speech recognition models (S2T, wav2vec2, and DeepSpeech2) all produced worse predictions than all of our main in-house models (Figure 2). This result is also consistent with the fact that speech recognition systems derived in engineering contexts do not reproduce some characteristics of human speech perception^76, 77^. These model shortcomings could relate to these models having been trained on clean speech (as is common for such systems given that they are typically combined with separate denoising systems) – when we explicitly manipulated background noise during training, we found that training without noise caused the in-house models to produce worse brain predictions. The inclusion of an initial cochlear stage may also help to produce representations and behavior like those of biological auditory systems^32^, though we did not manipulate that here.

### Relation to prior work

The best-known prior study along these lines is that of Kell et al., (2018), and the results here qualitatively replicate the key results of that study. One contribution of the present study thus lies in showing that these earlier results hold for many different auditory models. In particular, most trained models produce better predictions than the SpectroTemporal baseline model, and most exhibit a correspondence between model stages and brain regions. The consistently worse predictions obtained from models with random/permuted weights also replicates prior work, providing more evidence that optimizing representations for tasks tends to bring them in closer alignment with biological sensory systems. In addition, we substantiated these main conclusions using representational similarity analyses in addition to regression-based predictions, providing converging evidence for model-brain matches. Overall, the results indicate a qualitatively similar set of results to those obtained in the ventral visual pathway, where many different trained models produce overall good predictions^64^.

The Kell et al. study used a model trained on two tasks, but did not test the extent to which the multiple tasks improved the overall match to human brain responses. Here we compared model-brain similarity for models trained on single tasks and models trained on multiple tasks, and saw advantages for multiple tasks. We note that it is not always straightforward to train models to perform multiple tasks, and indeed that the Kell et al. study found that task performance was optimized when the representations subserving the two tasks were partially segregated. This representational segregation could potentially interact with the extent to which the model representations match to human brain responses. But for the tasks we considered here, it was not necessary to explicitly force representational segregation in order to achieve good task performance, or good predictions of human brain responses.

Beyond the study by Kell et al., there have been relatively few other efforts to compare neural network models to auditory cortical responses. One study compared representational similarity of fMRI responses to music to the responses of a neural network trained on music annotations, but did not compare to standard baseline models of auditory cortex^36^. Another study optimized a network for a small-scale (10 digit) word recognition task, and reported seeing some neurophysiological properties of the auditory system^38^. Koumura et al.^37, 107^ trained networks on environmental sound or speech classification, and observed tuning to amplitude modulation, similar to that found in peripheral and mid-level stages of biological auditory systems, but did not investigate the putative hierarchy of cortical regions. Giordano et al. found that three deep neural network models predicted non-primary voxel responses better than standard acoustic features, generally consistent with the results shown here^44^. Millet et al.^41^ used a self-supervised speech model to predict brain responses to naturalistic speech, and found a stage-region correspondence similar to that in Kell et al. and the present work. However, they used a model that we found to produce poor predictivity compared to others that we tested, and the overall variance explained was relatively low. Similarly, Vaidya et al.^43^ demonstrated that certain self-supervised speech models capture distinct stages of speech processing. Our results complement these findings in showing that they apply to a large set of models and to responses to natural sounds more generally.

### Limitations of our approach and results

The analyses presented here are intrinsically limited by the coarseness of fMRI data in space and time. Voxels contain many thousands of neurons, and the slow time constant of the BOLD signal averages the underlying neuronal responses on a timescale of several seconds. It remains possible that responses of single neurons would be harder to relate to the responses of the sorts of models tested here, particularly when temporal dynamics are examined. Our analyses are also limited by the number of stimuli that can feasibly be presented in an fMRI experiment (less than 200 given our current methods and reliability standards). It is possible that larger stimulus sets would better differentiate the models we tested.

The conclusions here are also limited by the two metrics of model-brain similarity that we used. The regression-based metric of explained variance is based on the assumption that representational similarity can be meaningfully assessed using a linear mapping between responses to natural stimuli^46, 108, 109^. This assumption is common in systems neuroscience, but could obscure aspects of a model representation that deviate markedly from those of the brain, because the linear mapping picks out only the model features that are predictive of brain responses. There is ample evidence that deep neural network models tend to rely partially on different features than humans^18, 19^, and have partially distinct invariances^25, 27^ for reasons that remain unclear. Encoding model analyses may mask the presence of such discrepant model properties. They also leave open the possibility that the features that are picked out by the linear mapping are not the same as those in the brain – they only need to be correlated with the brain features to produce good brain predictions on a finite dataset^57, 65^. We note that accurate predictions of brain responses may be useful in a variety of applied contexts, and so have value independent of the extent to which they capture intuitive notions of similarity. In addition, accurate predictive models might be scientifically useful in helping to better understand what is represented in a brain response (e.g., by generating predictions of stimuli that yield high or low responses, that can then be tested experimentally^110^). But there are nonetheless limitations when relying exclusively on regression to evaluate whether a model replicates brain representations.

Representational dissimilarity matrices complement regression-based metrics, but have their own limitations. RDMs are computed from the entirety of a representation, and so reflect all of its dimensions, but conversely are not invariant to linear transformations. Scaling some dimensions up and others down can alter an RDM even if it does not alter the information that can be extracted from the underlying representation. Further, RDMs rely on choosing a distance measure between model responses to construct the RDM and a distance measure between two RDMs, and the most commonly used measures do not obey formal properties of metric spaces^111^. Although the RDM comparisons we employ are in widespread use, the measurement of representational similarity is an active area of research, with alternative metrics under development^111^. Moreover, RDMs must be computed from sets of voxel responses, and so are sensitive to the (potentially ad hoc) choice of which voxels to pool together. For instance, our first analysis (Figure 2) pooled voxels across all of auditory cortex, and this may have limited the similarity observed with individual model stages (potentially explaining in part the modest advantage relative to the baseline model). By contrast, regression metrics can be evaluated on individual voxels.

The fact that the regression and RDM analyses yielded similar qualitative conclusions is reassuring, but they are but two of a large space of possible model-brain similarity metrics^57, 111,112,113^. In addition, the correspondence between the two metrics was not perfect. The correlation between overall variance explained (the regression metric) and the human-model RDM similarity across network stages was R^2^=0.56 and 0.58 for NH2015 and B2021, respectively – much higher than chance, but below the noise ceiling for the two measures (Supplementary Figure S10). In addition, the best model stages for each ROI were generally weakly correlated between the two metrics (Figure 7). These discrepancies are not well understood at present, but must eventually be resolved for the modeling enterprise to declare success.

Although we found consistent evidence for correspondence between model stages and brain regions, this correspondence was coarse: the best-predicting model stages tended to be later for non-primary voxels than for primary voxels. There is at present no evidence for anything more fine-grained (e.g., with each stage of a model mapping onto a distinct stage of the auditory system). The evaluation of a more fine-grained correspondence is limited in part by fMRI data. For instance, the cortex is composed of distinct layers of neurons which might be expected to map onto distinct stages of an artificial neural network, but our fMRI data do not resolve layers within a cortical column. The many subcortical stages of auditory processing might be expected to be captured by early model stages (as would be consistent with the relatively late position of the best-predicting stages for cortical voxels), but are difficult to measure as reliably as is needed for the model comparisons used here. The position of the best-predicting stage also varied a fair bit across models. We do not find this surprising given the diversity of tasks on which the models were trained, and the diversity of model architectures, but we also lack a theory at present for why the best-predicting stages are in particular places for particular models.

Another limitation is that we did not evaluate the behavior of the models we tested. In other work we have found that models trained to perform natural tasks on natural stimuli tend to reproduce a fairly wide range of human psychophysical results^31–33^, but we did not conduct such experiments on the present set of models, in part because we currently lack psychophysical batteries for some of the training tasks (specifically, speaker and audio event recognition). It is thus possible that the models learn to perform the training tasks differently than humans despite using features that enable above-baseline predictions of brain responses^23, 25, 27, 57, 114^.

As discussed above, our study is unable to disentangle effects of model architecture, task, and training data on the brain predictions of the external models tested. We emphasize that this was not our goal – we sought to test a wide range of models as a strong test of the generality of brain-DNN similarities, cognizant that this would limit our ability to assess the reasons for model-brain discrepancies. The in-house models nonetheless help reveal some of the factors that drive differences in model predictions.

### Future directions

The finding that task-optimized neural networks generally enable improved predictions of auditory cortical brain responses motivates a broader examination of such models, as well as further model development for this purpose. For instance, the findings that different tasks best predict different brain responses suggest that models that both recognize and produce speech might help to explain differences in “ventral” and “dorsal” speech pathways^115^, particularly if paired with branching architectures^31^ that can be viewed as hypotheses for distinct functional pathways. Models trained to localize sounds^33^ in addition to recognizing them might help explain aspects of the cortical encoding of sound location and its possible segregation from representations of sound identity^97, 116, 117, 118–120^. Task-optimized models could potentially also help clarify findings that currently do not have an obvious functional interpretation, for instance the tendency for responses to broadband onsets to be anatomically segregated from responses to sustained and tonal sounds^50, 121, 122^, if such response properties emerge for some tasks and not others.

On the other hand, it remains unclear whether improvements in tasks and training datasets can entirely bridge the substantial remaining gaps between current neural network models and the brain. In principle, current model failures (aberrant behaviors such as adversarial examples and discrepant metamers, along with sub-ceiling model-brain similarity according to many metrics) might be explained simply by differences in the training task or training data relative to those that biological organisms are plausibly optimized for^123^. For instance, models are often optimized for a single task, and on data that is arguably more stereotyped (e.g., with images centered on single objects^124, 125^) than the sensory data encountered by organisms in the world, and this might be expected to limit the match to human behavior, with improvements to be expected as tasks and data are made richer. Our results from training in noise and the multi-task networks are consistent with this general idea. However, it also remains possible that because biological sensory systems result from complex evolutionary histories, they might not be well approximated by the result of a single optimization procedure for a fixed set of training objectives^30^, which might limit the extent to which pure task optimization is likely to account for biological perception.

One can also question the role of tasks in the optimization that produces biological perceptual systems. As is widely noted, the learning algorithm used in most of the models we considered (supervised learning) is not a plausible account for how biological organisms incorporate data from their environment^126^. The use of supervised learning is motivated by the possibility that one could converge on an accurate model of the brain’s representations by replicating some constraints that shape neural representations even if the way those constraints are imposed deviates from biology. It is nonetheless conceivable (and perhaps likely) that fully accurate models will require learning algorithms that more closely resemble the optimization processes of biology, in which nested loops of evolutionary selection and (largely unsupervised) learning over development combine to produce systems that can perform a wide range of tasks well, and thus successfully pass on their genes. Some initial steps in this direction can be found in recent models that are optimized without labeled training data^41, 43, 127, 128^. Our model set contained one such contrastive self-supervised model (Wav2vec2^129^), and although its brain predictions were worse than those of most of the supervised models, this direction clearly merits extensive exploration.

It will also be important to use additional means of model evaluation, such as model-matched stimuli^25, 27, 57^, stimuli optimized for a model’s predicted response^110, 130,131,132^, directly substituting brain responses into models^112^, or recently proposed alternative methods to measure representational similarity^111^. These additional types of evaluations could help address some of the limitations discussed in the previous section. And ultimately, analyses such as these need to be related to more fine-grained anatomy and brain response measurements. Model-based analyses of human intracranial data and single neuron responses from nonhuman animals both seem like promising next steps in the pursuit of complete models of biological auditory systems.

## Methods

### Voxel response modeling

The following voxel encoding model methods are adapted from those of Kell et al., (2018) and where the methods are identical, we have reproduced the analogous sections of the methods verbatim. We summarize the minor differences from the Kell et al. methods at the end of this section. All voxel response modeling and analysis code was written in Python (version 3.6.10), making heavy use of the numpy^134^ (version 1.19.0), scipy^135^ (version 1.4.1), and scikit-learn^136^ libraries (version 0.24.1).

#### General

We performed an encoding analysis in which each voxel’s time-averaged activity was predicted by a regularized linear model of the DNN activity. We operationalized each model stage within each candidate model (see Section “Candidate models”) as a hypothesis of a neural implementation of auditory processing. The fMRI hemodynamic signal to which we were comparing the candidate model blurs the temporal variation of the cortical response, thus a fair comparison of the model to the fMRI data involved predicting each voxel’s time-averaged response to each sound from time-averaged model responses. We therefore averaged the model responses over the temporal dimension after extraction. Because it seemed possible that units with real-valued activations might average out to near-zero values, we extracted unit activations after model stages that transform the output to positive values (ReLU, Tanh stages). Transformer architectures had no such stages, so we extracted the real-valued unit activations, and analyzed all model stages in this way. Pilot analyses suggested that voxel predictions from these models were similar when we time-averaged unit activations that were exclusively positive (specifically, when we used the root-mean-square instead of the mean).

#### Voxelwise modeling: Regularized linear regression and cross-validation

We modeled each voxel’s time-averaged response as a linear combination of a model stage’s time-averaged unit responses. We first generated 10 randomly selected train/test splits of the 165 sound stimuli into 83 training sounds and 82 testing sounds. For each split, we estimated a linear map from model units to voxels on the 83 training stimuli and evaluated the quality of the prediction using the remaining 82 testing sounds (described below in greater detail). For each voxel-stage pair, we took the median across the 10 splits. The linear map was estimated using regularized linear regression. Given that the number of regressors (i.e., time-averaged model units) typically exceeded the number of sounds used for estimation (83), regularization was critical. We used L2-regularized (‘‘ridge’’) regression, which can be seen as placing a zero-mean Gaussian prior on the regression coefficients. Introducing the L2-penalty on the weights results in a closed-form solution to the regression problem, which is similar to the ordinary least-squares regression normal equation:

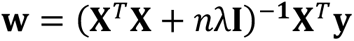

where **w** is a d-length column vector (the number of regressors – i.e., the number of time-averaged units for the given stage), **y** is an n-length column vector containing the voxel’s mean response to each sound (length 83), **X** is a matrix of regressors (n stimuli by d regressors), n is the number of stimuli used for estimation (83), and **I** is the identity matrix (d by d). We demeaned each column of the regressor matrix (i.e., each model unit’s response to each sound), but we did not normalize the columns to have unit norm. Similarly, we demeaned the target vector (i.e., the voxel’s or the component’s response to each sound). By not constraining the norm of each column to be one, we implemented ridge regression with a non-isotropic prior on each unit’s learned coefficient. Under such a prior, units with larger norm were expected a priori to contribute more to the voxel predictions. In pilot experiments, we found that this procedure led to more accurate and stable predictions in left-out data, compared with a procedure where the columns of the regressor matrices were normalized (i.e., with an isotropic prior). The demeaning was performed on the train set and the same transformation was applied on the test set. This ensured independence (no data leakage) between the train and test sets.

Performing ridge regression requires selecting a regularization parameter that trades off between the fit to the (training) data and the penalty for weights with high coefficients. To select this regularization parameter, we used leave-one-out cross validation within the set of 83 training sounds. Specifically, for each of 100 logarithmically-spaced regularization parameter values (1e-50, 1e-49, …, 1e48, 1e49), we measured the squared error in the resulting prediction of the left out sound using regression weights derived from the other sounds in the training split. We computed the average of this error (across the 83 training sounds) for each of the 100 potential regularization parameter values. We then selected the regularization parameter that minimized this mean squared error. Finally, with the regularization parameter selected, we used all 83 training sounds to estimate a single linear mapping from a stage’s features to a given voxel’s response. We then used this linear mapping to predict the response to the left-out 82 test sounds, and evaluated the Pearson correlation of the predicted voxel response with the observed voxel response. If the predicted voxel response had a standard deviation of exactly zero (no variance of the prediction across test sounds), the Pearson correlation coefficient was set to 0. Similarly, if the Pearson correlation coefficient was negative, indicating that the held-out test sounds were not meaningfully predicted by the linear map from the training set, the Pearson correlation value was similarly set to 0. We squared this Pearson correlation coefficient to yield a measure of variance explained. We found that the selected regularization parameter values rarely fell on the boundaries of the search grid, suggesting that the range of the search grid was appropriate. We emphasize that the 82 test sounds on which predictions were ultimately evaluated were not incorporated into the procedure for selecting the regularization parameter nor for estimating the linear mapping from stage features to a voxel’s response – i.e., the procedure was fully cross-validated.

Selecting regularization coefficients independently for each voxel-stage regression was computationally expensive, but seemed important for our scientific goals given that the optimal regularization parameter could vary across voxel-stage pairs. For instance, differences in the extent to which the singular value spectrum of the feature matrix is uniform or peaked (which influences the extent to which the **X***^T^***X** + *n*λ**I** matrix in the normal equation above is well-conditioned) can lead to differences in the optimal amount of regularization. Measurement noise, which varies across voxels can also influence the degree of optimal regularization. By allowing different feature sets (stages) to have different regularization parameters we are enabling each feature set to make the best possible predictions, which is appealing given that the prediction quality is the critical dependent variable that we compare across voxels and stages. Varying the regularization parameter across feature sets while predicting the same voxel response will alter the statistics of the regression coefficients across feature sets, and thus would complicate the analysis and interpretation of regression coefficients. However, we are not analyzing the regression coefficients in this work.

#### Voxelwise modeling: Correcting for reliability of the measured voxel response

The use of explained variance as a metric for model evaluation is inevitably limited by measurement noise. To correct for the effects of measurement noise we computed the reliability of both the measured voxel response and the predicted voxel response. Correcting for the reliability of the measured response is important to make comparisons across different voxels, because (as shown in e.g., Figure S2 in Kell et al., (2018)) the reliability of the BOLD response varies across voxels. This variation can occur for a variety of reasons (e.g., distance from the head coil elements). Not correcting for the reliability of the measured response will downwardly bias the estimates of variance explained and will do so differentially across voxels. This differential downward bias could lead to incorrect inferences about how well a given set of model features explains the response of voxels in different parts of auditory cortex.

#### Voxelwise modeling: Correcting for reliability of the predicted voxel response

Measurement noise corrupts the test data to which model predictions are compared (which we accounted for by correcting for the reliability of the measured voxel response, as described above), but noise is also present in the training data and thus also inevitably corrupts the estimates of the regression weights mapping from model features to a given voxel. This second influence of measurement noise is often overlooked, but can be addressed by correcting for the reliability of the predicted response. Doing so is important for two reasons. First, as with the reliability of the measured voxel response, not correcting for the predicted voxel response can yield incorrect inferences about how well a model explains different voxels. Second, the reliability of the predicted response for a given voxel can vary across feature sets, and failing to account for these differences can lead to incorrect inferences about which set of features best explains that voxel’s response. It was thus in practice important to correct for the reliability of the predicted voxel response. By correcting for both the reliability of the measured voxel response and the reliability of the predicted response, the ceiling of our measured r-squared values was 1 for all voxels and all stages, enabling comparisons of voxel predictions across all voxels and all neural network stages.

#### Voxelwise modeling: Corrected measure of variance explained

To correct for the reliability, we employ the correction for attenuation^61^. It is a standard technique in many fields, and is becoming more common in neuroscience. The correction estimates the correlation between two variables independent of measurement noise (here the measured voxel response and the model prediction of that response). The result is an unbiased estimator of the correlation coefficient that would be observed from noiseless data. Our corrected measure of variance explained was the following:

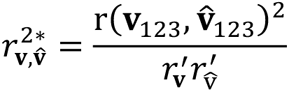

where **v**_123_ is the voxel response to the 82 left-out sounds averaged over the three scans, 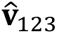 is the predicted response to the 82 left-out sounds (with regression weights learned from the other 83 sounds), r is a function that computes the correlation coefficient, 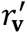 is the estimated reliability of that voxel’s response to the 83 sounds and 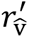 is the estimated reliability of that predicted voxel’s response. 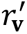 is the median of the correlation between all 3 pairs of scans (scan 0 with scan 1; scan 1 with scan 2; and scan 0 with scan 2), which is then Spearman-Brown corrected to account for the increased reliability that would be expected from tripling the amount of data ^61^. 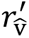 is analogously computed by taking the median of the correlations for all pairs of predicted responses (models fitted on a single scan) and Spearman-Brown correcting this measure. Note that for very noisy voxels, this division by the estimated reliability can be unstable and can cause for corrected variance explained measures that exceed one. To ameliorate this problem, we limited both the reliability of the prediction and the reliability of the voxel response to be greater than some value *k* ^58^. For *k* = 1, the denominator would be constrained to always equal one and thus the ‘‘corrected’’ variance explained measured would be identical to uncorrected value. For *k* = 0, the corrected estimated variance explained measure is unaffected by the value *k*. This k-correction can be seen through the lens of a bias-variance tradeoff: this correction reduces the amount of variance in the estimate of variance explained across different splits of stimuli, but does it at the expense of a downward bias of those variance explained metrics (by inflating the reliability measure for unreliable voxels). For 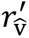, we used a k of 0.182, which is the p < 0.05 significance threshold for the correlation of two 83-dimensional Gaussian variables (i.e., with the same length as our 83-dimensional voxel response vectors used as the training set), while for 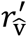 we used a k of 0.183 which is the p < 0.05 significance threshold for the correlation of two 82-dimensional Gaussian variables (i.e., same length as our 82-dimensional predicted voxel response vectors, the test set).

#### Voxelwise modeling: Summary

We repeated this procedure for each stage and voxel ten times, once each for 10 random train/test splits, and took the median explained variance across the ten splits for a given stage-voxel pair. We performed this procedure for all stages of all candidate models and all voxels (across two datasets: NH2015: 7694 voxels, B2021: 26,792 voxels). Thus, for each stage and voxel, this resulted in ten explained variance values (R^2^). We computed the median explained variance across these ten cross-validation splits for each voxel-stage pair. For comparison, we performed an identical procedure with the stages of a permuted network with the same architecture as our main networks (see Section “Candidate models with permuted weights*”*) and the SpectroTemporal baseline model. In all analyses, if a noise-corrected median explained variance value exceeded 1, we set the value to 1 to avoid an inflation of the explained variance.

In summary, the voxel prediction methods were largely the same as those in Kell et al., (2018), with the following differences. First, we imposed a different range of regularization constants to avoid hitting the bounds of the range. This difference was necessitated to accommodate a larger and more diverse set of models than in Kell et al. as well as changes to scikit learn in the years separating our study from that of Kell et al. Second, we set the r-squared values for negative r values to zero, rather than using signed r-squared values as in Kell et al. This seemed like the best choice given that negative r values indicates that a model cannot predict the data. Third, we used a different limit for the reliability used to correct the explained variance. Our limit was the minimum correlation that would be statistically significant for a sample size of 82 and 83 (which is the sample size for which the reliability is measured), whereas Kell assumed a sample size of 165. Fourth, we omitted the Fisher z transform when averaging r-squared values, as it seemed hard to motivate. We strongly suspect that none of these differences qualitatively affect any important result, but list them here for transparency.

#### Voxelwise predictions across models [Figure 2]

To compare how well each candidate model explained the variance in the fMRI data, we aggregated the explained variance across all voxels in the dataset of interest (NH2015: 7694 voxels, B2021: 26,792 voxels) for each model. We evaluated each candidate model using its best-predicting stage. Selection of the best-predicting model stage was performed in one of two ways. In the main analysis featured in Figure 2, for each voxel, we used half of the ten cross-validation test splits to select the best-predicting stage, and the remaining five test splits to obtain the median explained variance. This yielded a median explained variance per voxel. To ensure that this procedure did not depend on the random five cross-validation splits selected, we repeated this procedure ten times for each model. We then obtained the mean of the explained variance values for each voxel across these ten iterations. To mitigate concerns that this analysis might be affected by the overlap in sounds in the five splits used to select the best stage and the five splits used to measure the explained variance, we performed a second analysis in which we selected the best-predicting model stage using all the voxels for all but one participant, and then measured the explained variance in each of the voxels in the left-out participant. This analysis measures explained variance with data fully independent from that used to choose the best-stage, but is less consistent with the rest of the analyses (e.g., the maps of the best-predicting model stage, in which it was critical to choose the best-predicting stage separately for each voxel). We confirmed that the results shown in Figure 2 were qualitatively similar if this second procedure was used to choose the best-predicting stage for each model. To obtain an aggregated explained variance across voxels for each model, we first obtained the median across voxels within each participant, and then took the mean across participants. An identical procedure was used for the permuted networks.

#### Voxelwise predictions across model stages [Supplementary Figure S2]

To visualize how well each stage of each candidate model explained variance in the fMRI data, we aggregated the explained variance across all voxels in the dataset of interest (NH2015: 7694 voxels, B2021: 26,792 voxels) for each model. Given that no model stage selection procedure took place, we simply obtained the median across voxels within each participant, and then took the mean across participants for each model stage, identical to the aggregation procedure for the best stage voxelwise predictions (Figure 2).

#### Best-predicting model stage [Figure 6 / Figure 7]

We also examined which model stage best predicted each voxel’s response (an ‘‘argmax’’ analysis, i.e., the position that best predicts the response for each voxel). We assigned each model stage a position index between 0 and 1 (using minmax normalization such that the first stage was assigned a value of 0 and the last stage a value of 1, i.e.: (current_stage -min_stage) r/ (num_stages - min_stage)). For summary maps (Figure 6), we predicted responses in individuals and then aggregated results across participants (median), after they were aligned in a common coordinate system (i.e., the FsAverage surface from FreeSurfer) using K-Nearest Neighbor interpolation. For the across-model summary map we took the median of the best model stage positions across the n=15 best-performing models (rounded to the first decimal place). The plots were visualized using Freeview using default parameters (Freeview version 7.3.2). The color overlay was an inverse color wheel. The color scale limits were set to extend from 0 to the stage beyond the most common best stage (across voxels in both fMRI datasets). The permuted control networks were visualized using an identical color scale to the trained networks.

To quantify these summary maps, we compared the best-predicting model stage within different regions of the auditory cortex (Figure 7). We used four anatomical region-of-interest (ROIs): one for primary auditory cortex along with three ROIs for posterior, lateral, and anterior non-primary auditory cortex. These ROIs were combinations of subsets of ROIs in the Glasser et al. parcellation ^67^. We note that they were taken directly from a previous publication ^51^ where they were intended to capture the auditory cortical regions exhibiting reliable responses to natural sounds, and were not adapted in any way to the present analysis. For each model, we computed the relative model stage position of the best-predicting stage within each ROI (an ‘‘argmax’’ analysis, as shown on the summary maps, Figure 6), which we summarized by taking the median across voxels within each ROI for each participant followed by the mean across participants (similar to the aggregation procedure in Figure 2). This yielded an average relative best model stage position per candidate model within each ROI. An identical procedure was used for the permuted networks.

### Component modeling

We complemented the voxelwise modeling with analogous predictions of components of the fMRI response derived from all of the voxels. In previous work ^50^ we found that voxel responses to natural sounds can be explained as a linear combination of six response components (Figure 3A). The components are derived from the auditory cortical voxels (pooled across participants) that exceed a criterion of reliability. Each component is defined by a response to each of the sounds in the stimulus set, and has a weight for each voxel in the pool.

We predicted the responses for each of these six components in a manner similar to the voxelwise modeling. The only difference was that we did not perform any noise-ceiling correction for the components (the components do not have repetitions across scan sessions, unlike the voxel responses). Thus, all component predictions reported are the “raw” explained variance (squared Pearson correlation coefficient).

#### Component predictions across models [Figure 5 & Figure 9]

To compare how well each candidate model explained the component responses, we selected the best-predicting model stage for each candidate model using independent data (identical to the procedure described in “Voxelwise predictions across models” for the voxel data).

#### Component predictions across sounds [Figure 4]

We visualized the component response predictions by plotting them against the actual responses (derived from Norman-Haignere et al., (2015)) for each sound. We did this for the best-predicting model stage of the CochResNet50-MultiTask (best-performing model overall, Figure 2A) for each component. The best-predicting model stage was selected across ten iterations of the independent model stage selection procedure, as described in Section “Voxelwise predictions across models”. For each of the ten iterations, we used the median explained variance value of five random cross-validation test splits to identify the best model stage. Thus, ten iterations of this procedure yielded ten best-predicting model stages. For each component, we selected the most frequently occurring best-predicting model stage as the stage with which to visualize a given component’s predictions. Given a component and model stage, we obtained the predicted component response for a sound by taking the mean over all cross-validation splits in which that sound was included in the test set. In that way, we obtained the average prediction for each sound and each component. We note that the predicted component responses visualized in Figure 4 were demeaned during the regression procedure (using the training set mean to demean the test set, ensuring no data leakage between train/test sets, see Section: Voxelwise modeling: Regularized linear regression and cross validation) but the predicted responses were transformed back to their original scale (by adding back their mean) for visualization in Figure 4 (ordinate values). The actual component responses in Figure 4 were taken directly from Norman-Haignere et al., (2015) without any transformations (abscissa values).

### Representational Similarity Analysis

To assess the robustness of our conclusions to the evaluation metric, we also investigated the similarity of model and fMRI responses using Representational Similarity Analysis (RSA)^47, 63, 64^. We used the same set of model stages and time-averaged representations as were used in the regression-based voxelwise modeling analysis. To construct the model representational dissimilarity matrix (RDM) for each model and model stage, we computed the dissimilarity (1 minus the Pearson correlation coefficient) between the model activations evoked by each pair of sounds. Similarly, to construct the fMRI RDM, we computed the dissimilarity in voxel responses (1 minus the Pearson correlation coefficient) between all voxel responses from a participant to each pair of sounds. Before computing the RDMs from the fMRI or model responses, we z-scored the voxel or unit responses. As a measure of fMRI and model similarity, we computed the Spearman rank ordered correlation coefficient between the fMRI RDM and the model RDM.

### Representational Similarity Analysis: Across model comparison

In the analysis shown in Figure 2, we compared the RDMs computed across all voxels of a participant to the RDM computed from the time-averaged unit responses of each stage of each model. To choose the best-matching stage, we first generated 10 randomly selected train/test splits of the 165 sound stimuli into 83 training sounds and 82 testing sounds. For each split, we computed the RDMs for each model stage and for each participant’s fMRI data for the 83 training sounds. We then chose the model stage that yielded the highest Spearman ρ measured between the model stage RDM and the participant’s fMRI RDM. Using this model stage, we measured the model and fMRI RDMs from the test sounds and computed the Spearman ρ. We repeated this procedure ten times, once each for 10 random train/test splits, and took the median Spearman ρ across the ten splits. We performed this procedure for all candidate models and all participants (across two datasets: NH2015: 8 participants, B2021: 20 participants) and computed the mean Spearman ρ across participants for each model. For comparison, we performed an identical procedure with permuted versions of each neural network model, and with the SpectroTemporal baseline model.

### Representational Similarity Analysis: Noise ceiling

The representational similarity analysis is limited by measurement noise in the fMRI data. As an estimate of the RDM correlation that could be reasonably expected to be achieved between a model RDM and a single participant’s fMRI RDM, we calculated the correlation between one participant’s RDM and the average of all the other participant’s RDM. Within each dataset (NH2015 and B2021) we held out one participant and averaged the RDMs across the remaining participants. The RDMs were measured from the same 10 train/test splits of the 165 sounds described in the previous section, using the 82 test sounds for each split. We then calculated the Spearman ρ between the RDM from the held-out participant and the average participant RDM. We took the median Spearman ρ across the 10 splits of data to yield a single value for each participant. This procedure was repeated holding out each participant, and the noise ceiling shown in Figure 2E, G and Figure 8B is the mean across the measured value for each held out participant. This corresponds to the “lower bound” of the noise ceiling used in prior work^63^. We plotted the noise ceiling on the results graphs rather than noise-correcting the human-model RDM correlation to be consistent with prior modeling papers that have used this analysis^63, 64^.

### Representational Similarity Analysis: Best model stage analysis

We also examined which model stage best captured the RDM measured from each anatomical ROI (an ‘‘argmax’’ analysis). We assigned each model stage a position index between 0 and 1. Given that we only report the “argmax” for this analysis (and not the measured values), we used the full set of 165 sounds to compute the RDMs. For a given ROI, we measured each participant’s fMRI RDM computed on the voxels within the ROI. For each model, we computed the RDM for each stage and measured the Spearman ρ between the model-stage RDM and the fMRI ROI RDM. We measured the argmax across the stages for each model and each participant. We then took the mean of this position index across participants to yield an average relative best model stage per candidate model within each ROI. An identical procedure was used for the permuted networks.

### Representational Similarity Analysis: Visualization

For fMRI RDMs, we computed the RDM individually for each participant, and then averaged the RDM for visualization. Both fMRI and model RDMs are grouped and colored by the sound categories assigned in Norman-Haignere et al., (2015). fMRI RDMs for all auditory cortex voxels and for each ROI can be found in Supplementary Figure S1.

### Effective dimensionality

We investigated the effective dimensionality (ED) of the representations at each model stage as well as of the measured fMRI activity. The ED^137^ was calculated on the set of 165 sounds that were used for the brain comparisons. The ED was evaluated as:

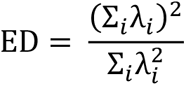

where *λ*_i_ are the square of the singular values obtained from the matrix of <activations> by <sounds>. This matrix was measured from the fMRI or model activations that were used for the regression analysis (demeaning each voxel or unit response) or for the RSA analysis (z-scoring the voxel or unit responses). In practice, these two different forms of pre-processing altered the ED measure, with ED values being about twice as large following the RSA pre-processing compared to the regression pre-processing. For instance, the ED for the fMRI data was 8.75 (for NH2015) and 5.32 (for B2021) using the regression pre-processing, but 16.9 (NH2015) and 12.9 (B2021) when using the representational similarity pre-processing. We used the two types of pre-processing to maintain consistency with the two types of model-brain similarity analysis, but the conclusions of the ED analysis would not have been qualitatively different had we exclusively used one type of pre-processing or the other.

### Statistical analysis

#### Voxel responses: Pairwise model comparisons (regression and representational similarity)

For statistical comparison of brain predictions between models trained with and without background noise, we evaluated significance non-parametrically by bootstrapping across participants (n=8 for NH2015, n=20 for B2021). For each model trained without background noise, we sampled the participant explained variance values with replacement (8 or 20 values, sampled 10,000 times) and took the average. The resulting histogram was compared to the noise-trained model’s observed value averaged across all participants. The p-value was obtained by counting the number of times the bootstrapped value was smaller than the observed value, divided by the number of bootstrap iterations (n=10,000). The test was one-tailed because it was motivated by the hypothesis that the model trained in background noise would produce better fMRI response predictions than the models trained without background noise.

#### Comparisons of best predicting model stages between ROIs (regression and representational similarity)

For statistical comparison of the mean model stage position index for pairs of anatomical ROIs (related to Figure 7), we performed a Wilcoxon signed rank test. The test compared the average values across models obtained from the primary ROI versus the average values obtained from the non-primary ROI across models. The test was two-tailed. In exploratory analyses of best predicting model stages among the three non-primary ROIs, we performed a Bonferroni correction for multiple comparisons due to the lack of a priori hypotheses.

#### Component responses: Pairwise model comparisons (regression)

For statistical comparison of component predictions between pairs of models (related to Figure 5 and 9), we evaluated significance non-parametrically via a permutation test. Based on the model stage selection procedure described in Section “Component predictions across models”, we obtained ten independently selected median explained variance values per component. For a given component, we took the average across the ten explained variance values for each model and then compute the difference between the two models. We generated a null distribution by randomly permuting the model assignment and measuring the difference between the average of these permuted model assignment lists. The p-value was obtained by counting the number of times the observed difference was smaller than the values measured from permuted data, divided by the number of permutations (n=10,000). The test was one-tailed because in each case it was motivated by a hypothesis that one model would produce better component response predictions than the other. Specifically, there were two types of comparisons. In the first, the trained models were compared to the SpectroTemporal baseline model. In the second, models trained on a task that was plausibly related to the selectivity of a component (e.g., the word task for the speech component) were compared to models trained on a task not obviously related to that component (e.g., the genre task for the speech component).

### fMRI data (NH2015)

The initial sections of the fMRI data collection methods used in Norman-Haignere et al., (2015) are very similar to the methods reported in Kell et al., (2018) and the text is replicated with minor edits.

#### fMRI cortical responses to natural sounds

The fMRI data analyzed here is a subset of the data in Norman-Haignere et al., (2015), only including the participants who completed three scanning sessions. Eight participants (four female, mean age: 22 years, range: 19-25; all right-handed) completed three scanning sessions (each ∼2 hours). Participants were non-musicians (no formal training in the five years preceding the scan), native English speakers, and had self-reported normal hearing. Two other participants only completed two scans and were excluded from these analyses, and three additional participants were excluded due to excessive head motion or inconsistent task performance. The decision to exclude these five participants was made before analyzing any of their fMRI data. All participants provided informed consent, and the Massachusetts Institute of Technology Committee on the Use of Humans as Experimental participants approved experiments (protocol number 2105000382).

#### Natural sound stimuli

The stimuli were a set of 165 two-second sounds selected to span the sorts of sounds that listeners most frequently encounter in day-to-day life (Norman-Haignere et al., 2015). All sounds were recognizable – i.e., classified correctly at least 80% of the time in a ten-way alternative forced choice task run on Amazon Mechanical Turk, with 55-60 participants per sound. See Supplementary Table S1 for names of all stimuli and category assignments. To download all 165 sounds, see the McDermott lab website: http://mcdermottlab.mit.edu/ downloads.html.

#### fMRI scanning procedure

Sounds were presented using a block design. Each block included five presentations of the identical two-second sound clip. After each two-second sound, a single fMRI volume was collected (‘‘sparse scanning’’), such that sounds were not presented simultaneously with the scanner noise. Each acquisition lasted one second and stimuli were presented during a 2.4 s interval (200 ms of silence before and after each sound to minimize forward/backward masking by scanner noise). Each block lasted 17 s (five repetitions of a 3.4 s TR). This design was selected based on pilot results showing that it gave more reliable responses than an event-related design given the same amount of overall scan time. Blocks were grouped into eleven runs, each with fifteen stimulus blocks and four blocks of silence. Silence blocks were the same duration as the stimulus blocks and were spaced randomly throughout the run. Silence blocks were included to enable estimation of the baseline response. To encourage participants to attend equally to each sound, participants performed a sound intensity discrimination task. In each block, one of the five sounds was 7 dB lower than the other four (the quieter sound was never the first sound). Participants were instructed to press a button when they heard the quieter sound.

#### fMRI data acquisition

MR data were collected on a 3T Siemens Trio scanner with a 32-channel head coil at the Athinoula A. Martinos Imaging Center of the McGovern Institute for Brain Research at MIT. Each functional volume consisted of fifteen slices oriented parallel to the superior temporal plane, covering the portion of the temporal lobe superior to and including the superior temporal sulcus. Repetition time (TR) was 3.4 s (although acquisition time was only 1 s), echo time (TE) was 30 ms, and flip angle was 90 degrees. For each run, the five initial volumes were discarded to allow homogenization of the magnetic field. In-plane resolution was 2.1 x 2.1 mm (96 x 96 matrix), and slice thickness was 4 mm with a 10% gap, yielding a voxel size of 2.1 x 2.1 x 4.4 mm. iPAT was used to minimize acquisition time. T1-weighted anatomical images were collected in each participant (1mm isotropic voxels) for alignment and surface reconstruction.

#### fMRI data preprocessing

Functional volumes were preprocessed using FSL and in-house MATLAB scripts. Volumes were corrected for motion and slice time. Volumes were skull-stripped, and voxel time courses were linearly detrended. Each run was aligned to the anatomical volume using FLIRT and BBRegister^138, 139^. These preprocessed functional volumes were then resampled to vertices on the reconstructed cortical surface computed via FreeSurfer^140^, and were smoothed on the surface with a 3mm FWHM 2D Gaussian kernel to improve SNR. All analyses were done in this surface space, but for ease of discussion we refer to vertices as ‘‘voxels’’ throughout this paper. For each of the three scan sessions, we estimated the mean response of each voxel (in the surface space) to each stimulus block by averaging the response of the second through the fifth acquisitions after the onset of each block (the first acquisition was excluded to account for the hemodynamic lag). Pilot analyses showed similar response estimates from a more traditional GLM. These signal-averaged responses were converted to percent signal change (PSC) by subtracting and dividing by each voxel’s response to the blocks of silence. These PSC values were then downsampled from the surface space to a 2mm isotropic grid on the FreeSurfer-flattened cortical sheet. For summary maps, we registered each participant’s surface to Freesurfer’s fsaverage template.

#### Voxel selection

For individual participant analyses, we used the same voxel selection criterion as Kell et al., (2018), selecting voxels with a consistent response to sounds from a large anatomical constraint region encompassing the superior temporal and posterior parietal cortex. Specifically, we used two criteria: (1) a significant response to sounds compared with silence (p < 0.001, uncorrected); and (2) a reliable response to the pattern of 165 sounds across scans. The reliability measure was as follows:

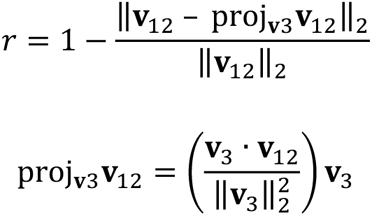

where **v**_12_ is the response of a single voxel to the 165 sounds averaged across the first two scans (a vector), and **v**_3_ is that same voxel’s response measured in the third. The numerator in the second term in the first equation is the magnitude of the residual left in **v**_12_ after projecting out the response shared with **v**_3_. This ‘‘residual magnitude’’ is divided by its maximum possible value (the magnitude of **v**_12_). The measure is bounded between 0 and 1, but differs from a correlation in assigning high values to voxels with a consistent response to the sound set, even if the response does not vary substantially across sounds. We found that using a more traditional correlation-based reliability measure excluded many voxels in primary auditory cortex because some of them exhibit only modest response variation across natural sounds. We included voxels with a value of this modified reliability measure of 0.3 or higher, which when combined with the sound responsive t test yielded a total of 7694 voxels across the eight participants (mean number of voxels per participant: 961.75; range: 637-1221).

### fMRI data (B2021)

#### fMRI cortical responses to natural sounds

The fMRI data analyzed here is from Boebinger et al., (2021)^51^. Twenty participants (fourteen female, mean age: 25 years, range: 18-34; all right-handed) completed three scanning sessions (each ∼2 hours). Half of these participants (n=10) were highly-trained musicians, with an average of 16.3 years of formal training (ranging from 11-23 years, SD = 2.5) that began before the age of seven^141^ and continued until the time of scanning. The other half of the participants (n=10) were non-musicians with less than two years of total music training, which could not have occurred either before the age of seven or within the five years preceding the time of scanning. All participants provided informed consent, and the Massachusetts Institute of Technology Committee on the Use of Humans as Experimental Subjects approved experiments (protocol number 2105000382).

#### Natural sound stimuli

The stimuli consisted of the set of 165 two-second natural sounds from Norman-Haignere et al., (2015), as well as 27 additional music and drumming clips from a variety of musical cultures, for a total of 192 sounds. For consistency with the Norman-Haignere et al., (2015) dataset, we constrained our analyses to the same set of 165 sounds.

#### fMRI scanning procedure

The fMRI scanning procedure was similar to the design of Norman-Haignere et al., (2015), except for the following minor differences. Each stimulus block consisted of three repetitions of an identical two-second sound clip, and lasted 10.2 s (three repetitions of a 3.4 s TR). Each of the three scanning sessions consisted of sixteen runs (for a total of 48 functional runs per participant), with each run containing twenty-four stimulus blocks and five silent blocks of equal duration that were evenly distributed throughout the run. The shorter stimulus blocks used in this experiment allowed each stimulus to be presented six times throughout the course of the 48 runs. To encourage participants to attend equally to each sound, participants performed a sound intensity discrimination task. In each block, either the second or third repetition was 12 dB lower, and participants were instructed to press a button when they heard the quieter sound.

#### fMRI data acquisition

The data acquisition parameters were similar to those from Norman-Haignere et al., (2015), with a few minor differences. MR data were collected on a 3T Siemens Prisma scanner with a 32-channel head coil at the Athinoula A. Martinos Imaging Center of the McGovern Institute for Brain Research at MIT. Each functional volume consisted of 48 slices oriented parallel to the superior temporal plane, covering the whole brain. However, all analyses were restricted to an anatomical mask encompassing the same portions of the temporal and parietal lobes as in Norman-Haignere et al., (2015). Repetition time (TR) was 3.4 s (TA = 1 s), echo time (TE) was 33 ms, and flip angle was 90 degrees. For each run, the four initial volumes were discarded to allow homogenization of the magnetic field. In-plane resolution was 2.1 x 2.1 mm (96 x 96 matrix), and slice thickness was 3 mm with a 10% gap, yielding a voxel size of 2.1 x 2.1 x 3.3 mm. An SMS acceleration factor of 4 was used in order to minimize acquisition time. T1-weighted anatomical images were collected in each participant (1mm isotropic voxels) for alignment and surface reconstruction.

#### fMRI data preprocessing

Preprocessing was identical to Norman-Haignere et al., (2015). However, the initial analyses of this dataset differ from Norman-Haignere et al., (2015) in that a GLM was used to estimate voxel responses rather than signal averaging, which was necessary due to the use of shorter stimulus blocks that caused more overlap between BOLD responses to different stimuli. For each of the three scan sessions, we estimated the mean response of each voxel (in the surface space) by modeling each block as a boxcar function convolved with a canonical hemodynamic response function (HRF). The model also included six motion regressors and a first-order polynomial noise regressor to account for linear drift in the baseline signal. The resulting voxel beta weights were then downsampled from the surface space to a 2mm isotropic grid on the FreeSurfer-flattened cortical sheet. For summary maps, we registered each participant’s surface to Freesurfer’s fsaverage template.

#### Voxel selection

The process of selecting voxels was identical to Norman-Haignere et al., (2015), except that the reliability of voxel responses was determined by comparing the vectors of 192 beta weights estimated separately for the two halves of the data (v1 = first three repetitions from runs 1-24, v2 = last three repetitions from runs 25-48). Voxels were selected using the following reliability measure:

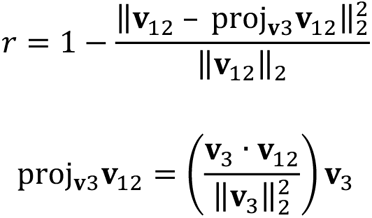

including voxels with a value of 0.3 or higher and further selecting only voxels with significant responses to sounds (p < 0.001, uncorrected). The combination of these two criteria yielded a total of 26,792 voxels across the twenty participants (mean number of voxels per participant: 1,340; range: 1,020 – 1,828).

### Candidate models

We investigated a set of n=19 candidate models. Nine of these models were trained by other labs for engineering purposes (“external”), and ten of these models were trained by us (“in-house”). Table 1 and Table 2 below show an overview of the nine external models and ten in-house models, respectively. For completeness, the SpectroTemporal baseline model is included in Table 2 along the in-house models. Details on model architectures and training can be found below the tables. For each model description, the model stage names match those in the code implementing the model. In the model-brain similarity analyses, we only included layers with learned parameters.

#### External models

Nine external models implemented in PyTorch were obtained from publicly available repositories. To our knowledge the requirement that models be available in PyTorch implementations (at the time our experiments were run) resulted in the exclusion of 3 models that we would have otherwise included: YAMNet^55, 156^ (https://github.com/tensorflow/models/tree/master/research/audioset/yamnet), DeepSpeech1^157^ ((https://github.com/mozilla/DeepSpeech), and NPC^158^ (https://github.com/Alexander-H-Liu/NPC). To accommodate the required dependencies, a separate software environment was created to run each model, and the versions of Python, PyTorch, and TorchAudio are reported separately for each model.

##### AST

Audio Spectrogram Transformer (AST) is an attention-based, convolution-free transformer architecture for audio classification.

We used the pretrained model available by Yuan Gong and colleagues as described in Gong et al., ^142^. Specifically, we used the model that was pre-trained on ImageNet^124^ using a vision transformer architecture (data-efficient image Transformer (DeiT)^159^) and afterwards trained on AudioSet^55^ (the best single model checkpoint which consisted of a model where all weights were averaged across model checkpoints from the first to last training epoch, model name: “Full AudioSet, 10 tstride, 10 fstride, with Weight Averaging (0.459 mAP)” (https://github.com/YuanGongND/ast).

AST is composed of an initial embedding layer followed by 12 multi-level encoder blocks that match the transformer architecture^160, 161^. Model activations were extracted at the output of each transformer encoder block. In addition to model activations from the transformer blocks, we extracted the initial embeddings that are fed to the model, as well as the final logits over AudioSet classes, yielding 14 layers in total.

As described in Gong et al., (2021), the audio input to AST is the raw audio waveform that is converted into a sequence of 128 log-mel filterbank features computed with 25 ms Hamming windows every 10 ms. As the model was trained on AudioSet, the input size to the model was 10.24s (1024 time frames). The model implementation zero-padded any input less than this length. Thus, the spectrogram was of size [1024, 128] ([n_temporal, n_spectral]), which in our analyses resulted from a zero-padded 2s audio clip. The spectrogram was normalized by subtracting the average value measured from the training dataset spectrograms (in this case, AudioSet), and dividing by two times the training dataset spectrogram standard deviation. The spectrogram was split into a sequence of 101 16×16 patches (see Gong et al., (2021) for details on the patch embedding procedure) with an overlap of 6 in both time and frequency (i.e., stride (10,10)). These patches were projected into an embedding of size 768 (“the patch embedding layer”) using the single convolutional layers as specified under “Architecture” in Gong et al., (2021). Two classification tokens were prepended to the embedding, which was then passed through 12 transformer encoder blocks. AST was trained on the full AudioSet dataset (consisting of the official balanced and full training set, i.e., around 2M segments) using cross-entropy. The final layer was a linear classification layer over 527 audio labels.

##### Architecture

The AST architecture is denoted below with the sizes of the tensors propagated through the network denoted in parentheses. Encoder refers to each transformer encoding block. Model stages that were used for voxel and component response modeling are denoted in bold. Here and elsewhere, the model stage names match those in the code implementing the model.

|  |  |
| --- | --- |
| Input | (1024,128) |
| **Embedding: Conv2d(1, 768, kernel_size=(16, 16), stride=(10, 10))** | **(12, 101, 768)** |
| **Encoder_1** | **(1214, 768)** |
| **Encoder_2** | **(1214, 768)** |
| **Encoder_3** | **(1214, 768)** |
| **Encoder_4** | **(1214, 768)** |
| **Encoder_5** | **(1214, 768)** |
| **Encoder_6** | **(1214, 768)** |
| **Encoder_7** | **(1214, 768)** |
| **Encoder_8** | **(1214, 768)** |
| **Encoder_9** | **(1214, 768)** |
| **Encoder_10** | **(1214, 768)** |
| **Encoder_11** | **(1214, 768)** |
| **Encoder_12** | **(1214, 768)** |
| **Linear_1(in_features=768, out_features=527, bias=True)** | **(1, 527)** |

For AST, we thus extracted model representations from the following 14 layers with the number of unit activations (regressors) for each sound denoted in parentheses: Embedding (768), Encoder_1 (768), Encoder_2 (768), Encoder_3 (768), Encoder_4 (768), Encoder_5 (768), Encoder_6 (768), Encoder_7 (768), Encoder_8 (768), Encoder_9 (768), Encoder_10 (768), Encoder_11 (768), Encoder_12 (768), Linear_1 (527).

*Extractions were performed using torch=1.8.1, torchaudio=0.8.1 in Python 3.8.11*.

##### DCASE2020 baseline

The DCASE2020 baseline model (henceforth DCASE2020) is recurrent architecture trained for automated audio captioning^143^, where the model accepts audio as input and outputs the textual description (i.e., the caption) of that signal. We used the pre-trained model implemented by Konstantinos Drossos and collaborators (https://github.com/audio-captioning/dcase-2020-baseline).

The input to the model is a log-mel spectrogram (audio was peak-normalized prior to spectrogram conversion, i.e., divided by the maximum value of the absolute value of the audio signal) with 64 frequency bins resulting from a short-time Fourier transform applying 23 ms windows (window size) every 11.5 ms (stride). This yields log spectrogram patches of 173 x 64 bins that are the inputs to the model, i.e., a 2D array [n_temporal, n_spectral]. These spectrograms are passed through a 3-layer bi-directional GRU encoder and a one bi-directional layer GRU decoder with a linear readout. There are residual connections between the second and third encoder GRUs. The linear readout is a linear projection into C classes representing the 4367 one-hot encoding of unique caption words. The decoder iterates for 22 time steps.

DCASE2020 was trained using cross-entropy loss on the development split of Clotho v1^144^ which consists of 2893 audio clips with 14465 captions. The audio samples are of 15 to 30 seconds duration, each audio sample having five captions of length 8-20 words.

##### Architecture

The DCASE2020 architecture is denoted below with the sizes of the tensors propagated through the network denoted in parentheses. Model stages that were used for voxel and component response modeling are denoted in bold. The two outputs of bidirectional recurrent stages were concatenated (i.e., treated as different features).

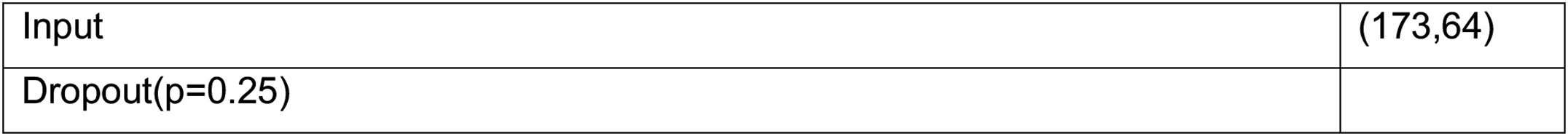
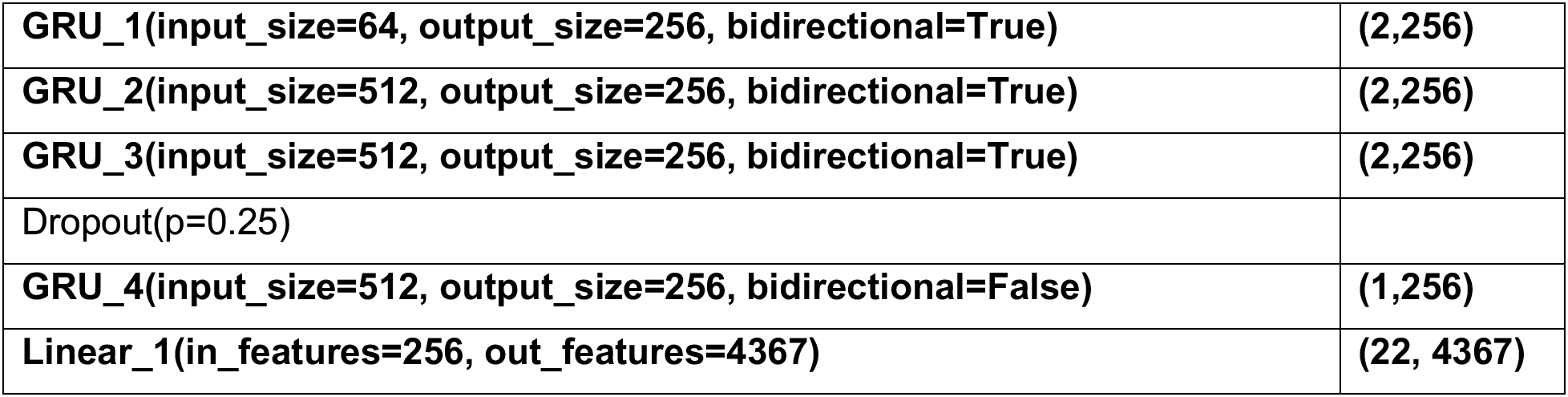

|  |  |
| --- | --- |
| Input | (173,64) |
| Dropout(p=0.25) |  |
| **GRU_1(input_size=64, output_size=256, bidirectional=True)** | **(2,256)** |
| **GRU_2(input_size=512, output_size=256, bidirectional=True)** | **(2,256)** |
| **GRU_3(input_size=512, output_size=256, bidirectional=True)** | **(2,256)** |
| Dropout(p=0.25) |  |
| **GRU_4(input_size=512, output_size=256, bidirectional=False)** | **(1,256)** |
| **Linear_1(in_features=256, out_features=4367)** | **(22, 4367)** |

Thus, for DCASE, we extracted model representations from the following 5 layers with the number of unit activations (regressors) for each sound denoted in parentheses: GRU_1 (512), GRU_2 (512), GRU_3 (512), GRU_4 (256), Linear_1 (4367).

*Extractions were performed using torch=1.3.1 in Python 3.7.10*.

##### DeepSpeech2

DeepSpeech2 is a recurrent architecture for automatic speech recognition^145^. We used the pre-trained PyTorch model by Sean Naren and collaborators (https://github.com/SeanNaren/deepspeech.pytorch).

As described by Amodei et al., ^145^, the input to the model is a log-spectrogram with 161 frequency bins resulting from a short-time Fourier transform applying 20 ms windows (window size) every 10 ms (stride). This yields log spectrogram patches of 201 x 161 bins that are the inputs to the model, i.e., a 2D array [n_temporal, n_spectral]. Each spectrogram was normalized by subtracting the mean spectrogram value and dividing by the standard deviation. These spectrograms were transformed by two 2D convolutional layers followed by five bidirectional recurrent Long Short-Term Memory (LSTM) layers and ending in a fully connected layer. The fully connected layer is a linear projection into C classes representing the vocabulary of the task. The vocabulary consists of 29 classes (output features), corresponding to English characters and space, apostrophe, blank. DeepSpeech2 was trained using a CTC loss on the Librispeech corpus^146^ (960hrs).

##### Architecture

The DeepSpeech2 architecture is denoted below with the sizes of the tensors propagated through the network denoted in parentheses. Model stages that were used for voxel and component response modeling are denoted in bold. The two outputs of bidirectional recurrent stages (using the LSTM output cell states) were concatenated (i.e., treated as different features).

|  |  |
| --- | --- |
| Input | (201, 161) |
| Conv2d_1(in_channels=1, out_channels=32, kernel_size=(41,11), stride=(2,2), padding=(20,5)) | (32, 81, 101) |
| BatchNorm2d_1(num_features=32) | (32, 81, 101) |
| **HardTanh_1(min_val=0, max_val=20)** | **(32, 81, 101)** |
| Conv2d_2(in_channels=32, out_channels=32, kernel_size=(21,11), stride=(2,1), padding=(10,5)) | (32, 41, 101) |
| BatchNorm2d_2(num_features=32) | (32, 41, 101) |
| **HardTanh_2(min_val=0, max_val=20)** | **(32, 41, 101)** |
| **LSTM_1(input_size=1312, hidden_size=1024, bidirectional=True)** | **(2, 1024)** |
| SequenceWise BatchNorm1d_1(num_features=1024) | (101, 1024) |
| **LSTM_2(input_size=1024, hidden_size=1024, bidirectional=True)** | **(2, 1024)** |
| SequenceWise BatchNorm1d_2(num_features=1024) | (101, 1024) |
| **LSTM_3(input_size=1024, hidden_size=1024, bidirectional=True)** | **(2, 1024)** |
| SequenceWise BatchNorm1d_3(num_features=1024) | (101, 1024) |
| **LSTM_4(input_size=1024, hidden_size=1024, bidirectional=True)** | **(2, 1024)** |
| SequenceWise BatchNorm1d_4(num_features=1024) | (101, 1024) |
| **LSTM_5(input_size=1024, hidden_size=1024, bidirectional=True)** | **(2, 1024)** |
| BatchNorm1d_5(num_features=1024) | (101, 1024) |
| **Linear_1(in_features=1024, out_features=29)** | **(101, 29)** |

Thus, for DeepSpeech2, we extracted model representations from the following 8 layers with the number of unit activations (regressors) for each sound denoted in parentheses: HardTanh_1 (2,592), HardTanh_2 (1,312), LSTM_1 (2,048), LSTM_2 (2,048), LSTM_3 (2,048), LSTM_4 (2,048), LSTM_5 (2,048), Linear_1 (29).

*Extractions were performed using torch=1.7.1, torchaudio=0.7.2 in Python 3.6.13*.

##### MetricGAN

MetricGAN+ (henceforth MetricGAN) is a generative adversarial network (GAN) for speech enhancement. We used the pretrained model available by SpeechBrain^162^ (hosted by HuggingFace) as described in Fu et al., ^147^. Specifically, we used a model that was pre-trained on the Voicebank-DEMAND dataset^148^ (training files: 20,000 (58.03hr) + validation files: 5,000 (14.65hr)) (https://huggingface.co/speechbrain/metricgan-plus-voicebank).

The generator of MetricGAN is a Bidrectional Long Short-Term Memory (BLSTM) with two bidirectional LSTM layers followed by two fully-connected layers. The objective of the generator is to estimate a mask consisting of the noise in the signal in order to generate clean speech. The discriminator of MetricGAN consists of a convolutional architecture (not investigated here).

As described in Fu et al., (2021), the audio input to the model is the magnitude spectrogram resulting from a short-time Fourier transform applying 32 ms (window size) windows every 16 ms (stride) resulting in 256 power frequency bins. This yields magnitude spectrogram patches of 126 x 256 bins that are the inputs to the model, i.e., a 2D array [n_temporal, n_spectral] that are passed through the BLSTM and linear layers of the generator model.

##### Architecture

The MetricGAN architecture is denoted below with the sizes of the tensors propagated through the network denoted in parentheses. Model stages that were used for voxel and component response modeling are denoted in bold. The two outputs of bidirectional recurrent stages (using the LSTM output cell states) were concatenated (i.e., treated as different features).

|  |  |
| --- | --- |
| **Input** | **(126, 257)** |
| **LSTM_1(input_size=257, hidden_size=200, bidirectional=True)** | **(2, 200)** |
| **LSTM_2(input_size=257, hidden_size=200, bidirectional=True)** | **(2, 200)** |
| Linear_1(in_features=400, out_features=300, bias=True) | (126, 300) |
| **LeakyReLU_1** | **(126, 300)** |
| **Linear_2(in_features=300, out_features=257, bias=True)** | **(126, 257)** |

For MetricGAN, we thus extracted model representations from the following 4 layers with the number of unit activations (regressors) for each sound denoted in parentheses: LSTM_1 (400), LSTM_2 (400), LeakyReLU_1 (300), Linear_2 (257).

*Extractions were performed using torch=1.9.1, speechbrain=0.5.10, huggingface-hub=0.0.17 in Python 3.8.11*.

##### S2T

S2T (also known as Speech-to-Text) is an attention-based transformer architecture for automatic speech recognition (ASR) and speech-to-text translation (ST). We used the pre-trained model available by HuggingFace^163^ as described in Wang et al., ^149^. Specifically, we used the large model trained on Librispeech corpus^146^ (960hrs) (https://huggingface.co/facebook/s2t-large-librispeech-asr).

S2T is an encoder-decoder model. The encoder part is composed of two convolutional layers followed by 12 multi-level encoder blocks that match the transformer architecture^160, 161^. Model activations were extracted at the output of each transformer encoder block. In addition to model activations from the transformer blocks, we extracted the initial embeddings that were fed to the model, yielding 13 layers in total. We did not investigate the decoder part of the model.

As described by Wang et al., (2020), the audio input to S2T is a log-mel spectrogram with 80 mel-spaced frequency bins resulting from a short-time Fourier transform applying 25 ms windows every 10 ms. Each spectrogram was normalized by subtracting the mean value of the spectrogram and dividing by the standard deviation. This yields the log-mel spectrogram of 198 x 80 bins that are the inputs to the model, i.e., a 2D array [n_temporal, n_spectral]. The spectrogram is passed through two convolutional layers before it is then passed through the 12 transformer encoder blocks. S2T was trained using cross-entropy loss, and the output consists of the 10K unigram vocabulary from SentencePiece^164^.

##### Architecture

The S2T architecture is denoted below with the sizes of the tensors propagated through the network denoted in parentheses (which is determined by the total stride in the initial feature encoder part of the architecture; not investigated here). Encoder refers to each transformer encoding block. Model stages that were used for voxel and component response modeling are denoted in bold.

|  |  |
| --- | --- |
| Input | (198, 80) |
| **Embedding / input post feature encoder** | **(50, 1024)** |
| **Encoder_1** | **(50, 1024)** |
| **Encoder_2** | **(50, 1024)** |
| **Encoder_3** | **(50, 1024)** |
| **Encoder_4** | **(50, 1024)** |
| **Encoder_5** | **(50, 1024)** |
| **Encoder_6** | **(50, 1024)** |
| **Encoder_7** | **(50, 1024)** |
| **Encoder_8** | **(50, 1024)** |
| **Encoder_9** | **(50, 1024)** |
| **Encoder_10** | **(50, 1024)** |
| **Encoder_11** | **(50, 1024)** |
| **Encoder_12** | **(50, 1024)** |

Thus, for S2T, we extracted model representations from the following 13 layers with the number of unit activations (regressors) for each sound denoted in parentheses: Embedding (1024), Encoder_1 (1024), Encoder_2 (1024), Encoder_3 (1024), Encoder_4 (1024), Encoder_5 (1024),

Encoder_6 (1024), Encoder_7 (1024), Encoder_8 (1024), Encoder_9 (1024), Encoder_10 (1024), Encoder_11 (1024), Encoder_12 (1024).

*Extractions were performed using transformers=4.10.0, torch=1.9.0, huggingface-hub=0.0.16 in Python 3.8.11*.

##### SepFormer

SepFormer (also known as Separation Transformer) is an attention-based transformer architecture for speech separation. We used the pretrained model available by SpeechBrain^162^ (hosted by HuggingFace) as described in Subakan et al., ^150^. Specifically, we used a model that was pre-trained on the WHAMR! dataset^151^ (training files: 20,000 (58.03hr) + validation files: 5,000 (14.65hr)) (https://huggingface.co/speechbrain/sepformer-whamr).

SepFormer is composed of an initial encoder followed by 32 multi-level dual-path encoder blocks similar to the transformer architecture^160, 161^. followed by a decoder. The transformer blocks follow a dual-path framework consisting of transformer blocks that model the short-term dependencies (IntraTransformer, IntraT), and Transformer blocks that model longer-term dependencies (InterTransformer, InterT). There are respectively 8 such IntraT and InterT blocks, yielding 16 transformer blocks, which is then repeated twice, yielding 32 transformer blocks in total. The objective of the dual-path transformer architecture is to estimate optimal masks to separate the audio sources present in the audio mixtures. The model was trained using scale-invariant source-to-noise ratio (SI-SNR) loss.

As described in Subakan et al., (2021), the audio input to AST is the raw audio waveform that is transformed by a single convolutional layer (encoder) followed by chunking the temporal dimension into patches of size 250. These chunks were then passed through the 32 transformer encoder blocks.

Model activations were extracted at the output of each transformer encoder block. In addition to model activations from the transformer blocks, we extracted the initial encoder embeddings that are fed to the model, yielding 33 layers in total.

##### Architecture

The SepFormer architecture is denoted below with the sizes of the tensors propagated through the network denoted in parentheses. Encoder refers to each transformer encoding block. Model stages that were used for voxel and component response modeling are denoted in bold.

|  |  |
| --- | --- |
| Input | (1, 16000) |
| Embedding: Conv1d(1, 256, kernel_size=(16,), stride=(8,), bias=False) | (256, 1999) |
| **ReLU_1()** | **(256, 1999)** |
| Conv1d(256, 256, kernel_size=(1,), stride=(1,), bias=False) | (256, 1999) |
| **Encoder_1** | **(18, 250, 256)** |
| **Encoder_2** | **(18, 250, 256)** |
| ... | ... |
| **Encoder_31** | **(18, 250, 256)** |
| **Encoder_32** | **(18, 250, 256)** |

For SepFormer, we thus extracted model representations from the following 33 layers with the number of unit activations (regressors) for each sound denoted in parentheses: Embedding (after ReLU) (256), Encoder_1 (256), Encoder_2 (256), …, Encoder_31 (256), Encoder_32 (256).

*Extractions were performed using torch=1.9.1, speechbrain=0.5.10, huggingface-hub=0.0.17 in Python 3.8.11*.

##### VGGish

VGGish is a convolutional architecture for audio classification inspired by the VGG model for image recognition^165^. VGGish converts audio input features into a semantically meaningful, 128-dimensional embedding. We used the pretrained VGGish by Hershey et al., (2017)^152^ (https://github.com/tensorflow/models/tree/master/research/audioset, specifically the PyTorch-compatible port by Harri Taylor and collaborators as found here: https://github.com/harritaylor/torchvggish).

VGGish was trained on the YouTube-100M corpus (70M training videos, 5.24 million hours with 30,871 labels)^152^. The videos average 4.6 minutes and are (machine) labeled with 5 labels on average per video from the set of 30,871 labels. The model was trained to predict the video-level labels based on audio information using a cross-entropy loss function. As described by Hershey et al., (2017), the audio input consists of 960 ms audio frames that are decomposed with a short-time Fourier transform applying 25 ms (window size) windows every 10 ms (stride) resulting in 64 log mel-spaced frequency bins. This yields log-mel spectrogram patches of 96 x 64 bins that are the inputs to the model, i.e., a 3D array [n_frames, n_temporal, n_spectral]. Given that VGGish contained an additional temporal dimension (the n_frames dimension), we averaged over both temporal dimensions (the n_temporal dimension which corresponds to the time dimension of the spectrogram as well as the n_frames dimension which corresponds to the batch dimension) to obtain a time-averaged model representation.

##### Architecture

The VGGish architecture is denoted below with the sizes of the tensors propagated through the network denoted in parentheses. Model stages that were used for voxel and component response modeling are denoted in bold.

|  |  |
| --- | --- |
| Input | (2, 96, 64) |
| Conv2d_1(in_channels=1, out_channels=64, kernel_size=(3,3), stride=(1,1), padding=(1,1)) | (2, 64, 96, 64) |
| **ReLU_1()** | **(2, 64, 96, 64)** |
| **MaxPool2d_1(kernel_size=2, stride=2, padding=0)** | **(2, 64, 48, 32)** |
| Conv2d_2(in_channels=64, out_channels=128, kernel_size=(3,3), stride=(1,1), padding=(1,1)) | (2, 128, 48, 32) |
| **ReLU_2()** | **(2, 128, 48, 32)** |
| **MaxPool2d_2(kernel_size=2, stride=2, padding=0)** | **(2, 128, 24, 16)** |
| Conv2d_3(in_channels=128, out_channels=256, kernel_size=(3,3), stride=(1,1), padding=(1,1)) | (2, 256, 24, 16) |
| **ReLU_3()** | **(2, 256, 24, 16)** |
| Conv2d_4(in_channels=256, out_channels=256, kernel_size=(3,3), stride=(1,1), padding=(1,1)) | (2, 256, 24, 16) |
| **ReLU_4()** | **(2, 256, 24, 16)** |
| **MaxPool2d_4(kernel_size=2, stride=2, padding=0)** | **(2, 256, 12, 8)** |
| Conv2d_5(in_channels=256, out_channels=512, kernel_size=(3,3), stride=(1,1), padding=(1,1)) | (2, 512, 12, 8) |
| **ReLU_5()** | **(2, 512, 12, 8)** |
| Conv2d_6(in_channels=512, out_channels=512, kernel_size=(3,3), stride=(1,1), padding=(1,1)) | (2, 512, 12, 8) |
| **ReLU_6()** | **(2, 512, 12, 8)** |
| **MaxPool2d_6(kernel_size=2, stride=2, padding=0)** | **(2, 512, 6, 4)** |
| Linear_1(in_features=12288, out_features=4096) | (2, 4096) |
| **ReLU_7()** | **(2, 4096)** |
| Linear_2(in_features=4096, out_features=4096) | (2, 4096) |
| **ReLU_8()** | **(2, 4096)** |
| Linear_3(in_features=4096, out_features=128) | (2, 128) |
| **ReLU_9()** | **(2, 128)** |

Thus, for VGGish, we extracted model representations from the following 13 layers with the number of unit activations (regressors) for each sound denoted in parentheses: ReLU_1 (4,096), MaxPool2d_1 (2,048), ReLU_2 (4,096), MaxPool2d_2 (2,048), ReLU_3 (4,096), ReLU_4 (4,096), MaxPool2d_3 (2,048), ReLU_5 (4,096), ReLU_6 (4,096), MaxPool2d_4 (2,048), ReLU_7 (4,096), ReLU_8 (4,096), ReLU_9 (128).

*Extractions were performed using torch=1.8.0 and torchaudio=0.8.1 in Python 3.8.5*.

##### VQ-VAE (ZeroSpeech2020)

Vector-quantized variational autoencoder (henceforth VQ-VAE) is an encoder-decoder architecture trained to synthesize speech in a target speaker’s voice. The model was trained for the ZeroSpeech 2020 challenge^166^. We used the pretrained model by Benjamin van Niekerk and colleagues as described in Niekerk et al., ^153^ (https://github.com/bshall/ZeroSpeech).

VQ-VAE consists of a CNN-based encoder and an RNN-based decoder. The encoder encodes the audio spectrogram and the decoder produces the new sound waveform. The model maps speech into a discrete latent space before reconstructing the original waveform.

As described by Niekerk et al., (2020), the input to VQ-VAE is the log-mel spectrogram (audio was peak-normalized prior to spectrogram conversion by dividing by the maximum of the absolute value of the audio signal, and this signal was multiplied by 0.999) with 80 mel-spaced frequency bins resulting from a short-time Fourier transform applying 25 ms windows every 10 ms. This yields the log-mel spectrogram of 201 x 80 bins that are the inputs to the model, i.e., a 2D array [n_temporal, n_spectral]. These spectrograms were transformed by five 1D convolutional layers. The model was trained to maximize the log-likelihood of the waveform given the discretized latent space bottleneck (details in Niekerk et al., (2020)). The model was trained on the ZeroSpeech 2019 English dataset consisting of the Train Voice Dataset (4h40min) and the Train Unit Dataset (15h40min)^154^. To extract model activations, the audio samples were converted to the first encountered speaker ID on the available speaker ID list (“S015”).

##### Architecture

The VQ-VAE encoder architecture is denoted below with the sizes of the tensors propagated through the network denoted in parentheses. We did not investigate the decoder part of VQ-VAE. Model stages that were used for voxel and component response modeling are denoted in bold.

|  |  |
| --- | --- |
| Input | (80,201) |
| Conv1d_1(80, 768, kernel_size=(3,), stride=(1,), bias=False) | (768,199) |
| BatchNorm1d_1(768, eps=1e-05, momentum=0.1, affine=True, track_running_stats=True) | (768,199) |
| **ReLU_1()** | **(768,199)** |
| Conv1d(768, 768, kernel_size=(3,), stride=(1,), padding=(1,), bias=False) | (768,199) |
| BatchNorm1d(768, eps=1e-05, momentum=0.1, affine=True, track_running_stats=True) | (768,199) |
| **ReLU_2()** | **(768,199)** |
| Conv1d(768, 768, kernel_size=(4,), stride=(2,), padding=(1,), bias=False) | (768,99) |
| BatchNorm1d(768, eps=1e-05, momentum=0.1, affine=True, track_running_stats=True) | (768,99) |
| **ReLU_3()** | **(768,99)** |
| Conv1d(768, 768, kernel_size=(3,), stride=(1,), padding=(1,), bias=False) | (768,99) |
| BatchNorm1d(768, eps=1e-05, momentum=0.1, affine=True, track_running_stats=True) | (768,99) |
| **ReLU_4()** | **(768,99)** |
| Conv1d(768, 768, kernel_size=(3,), stride=(1,), padding=(1,), bias=False) | (768,99) |
| BatchNorm1d(768, eps=1e-05, momentum=0.1, affine=True, track_running_stats=True) | (768,99) |
| **ReLU_5()** | **(768,99)** |

Thus, for VQ-VAE, we extracted model representations from the following 5 layers with the number of unit activations (regressors) for each sound denoted in parentheses: ReLU_1 (768), ReLU_2 (768), ReLU_3 (768), ReLU_4 (768), ReLU_5 (768).

*Extractions were performed using torch=1.9.0 in Python 3.8.11*.

##### Wav2vec 2.0

Wav2vec 2.0 (henceforth Wav2vec2) is a self-supervised transformer architecture for automatic speech recognition that learns representations of speech from masked parts of raw audio. We used the pretrained model from Huggingface Transformers^163^; original model can be found here: https://github.com/facebookresearch/fairseq/tree/main/examples/wav2vec#wav2vec-20). Specifically, we used the base version trained and fine-tuned on the Librispeech corpus^146^ (960hrs) (https://huggingface.co/facebook/wav2vec2-base-960h).

Wav2vec2 is composed of an initial multi-layer convolutional feature encoder followed by 12 multi-level encoder blocks that match the transformer architecture^160, 161^. Model activations were extracted at the output of each transformer encoder block. In addition to model activations from the transformer blocks, we extracted the initial embeddings that are fed to the model, as well as the final logits over character tokens, yielding 14 layers in total.

As described by Baevski et al., ^129^ the audio input to Wav2vec2 is a sound waveform of zero mean and unit variance. Wav2vec2 is trained via a contrastive task where the true speech input is masked in a latent space and has to be distinguished from distractors. The contrastive loss is augmented by a diversity loss to encourage the model to use samples equally often. The pre-trained model is fine-tuned for speech recognition by adding a linear projection on top of the network into C classes representing the vocabulary of the task by minimizing a CTC loss^167^. The vocabulary consists of 32 classes (output features), corresponding to English characters + bos_token=‘<s>’, eos_token=’</s>’, unk_token=‘<unk>’, pad_token=‘<pad>’, word_delimiter_token=’|’.

##### Architecture

The Wav2vec2 architecture is denoted below with the sizes of the tensors propagated through the network denoted in parentheses (which is determined by the total stride in the initial feature encoder part of the architecture; not investigated here). Encoder refers to each transformer encoding block. Model stages that were used for voxel and component response modeling are denoted in bold.

|  |  |
| --- | --- |
| Input | (32000) |
| **Embedding / input post feature encoder** | **(99, 768)** |
| **Encoder_1** | **(99, 768)** |
| **Encoder_2** | **(99, 768)** |
| **Encoder_3** | **(99, 768)** |
| **Encoder_4** | **(99, 768)** |
| **Encoder_5** | **(99, 768)** |
| **Encoder_6** | **(99, 768)** |
| **Encoder_7** | **(99, 768)** |
| **Encoder_8** | **(99, 768)** |
| **Encoder_9** | **(99, 768)** |
| **Encoder_10** | **(99, 768)** |
| **Encoder_11** | **(99, 768)** |
| **Encoder_12** | **(99, 768)** |
| **Linear_1** | **(99, 768)** |

For Wav2vec2, we thus extracted model representations from the following 14 layers with the number of unit activations (regressors) for each sound denoted in parentheses: Embedding (768), Encoder_1 (768), Encoder_2 (768), Encoder_3 (768), Encoder_4 (768), Encoder_5 (768),

Encoder_6 (768), Encoder_7 (768), Encoder_8 (768), Encoder_9 (768), Encoder_10 (768), Encoder_11 (768), Encoder_12 (768), Linear_1 (32).

*Extractions were performed using transformers=4.10.0, torch=1.9.0, huggingface-hub=0.0.16 in Python 3.8.11*.

#### In-house models

The in-house models consisted of a fixed cochleagram stage followed by either a convolutional architecture similar to that used in Kell et al., (2018) or a ResNet50 architecture. We refer to the full model architectures as CochCNN9 (indicating the 9 stages of this model) and CochResNet50. The models were trained either on the Word-Speaker-Noise dataset^25^, which supports three different tasks (word, speaker and audio event recognition), or the musical genre dataset compiled in Kell et al., (2018). In house models were trained and evaluated with Python 3.8.2 and PyTorch 1.5.0.

##### Cochleagram inputs

The SpectroTemporal model and all CochResNet50 and CochCNN9 architectures had a cochleagram representation as the input to the model. A cochleagram is a time-frequency representation of the audio with frequency bandwidth and spacing that mimics the human ear, followed by a compressive nonlinearity^105, 168^. The audio waveform passes through a bank of 211 bandpass filters ranging from 50Hz to 10kHz. Audio was sampled at 20kHz for the SpectroTemporal and the Word, Speaker, and AudioSet task models, and was sampled at 16kHz for the genre task models. Filters are zero-phase with frequency response equal to the positive portion of a single period of a cosine function. Filter spacing was set by the Equivalent Rectangular Bandwidth (ERB _N_) scale. Filters perfectly tile the spectrum such that the summed square response across all frequencies is flat, which includes four low-pass and four high-pass filters. The envelope was extracted from each filter subband using the magnitude of the analytic signal (Hilbert transform), and the envelopes were raised to the power of 0.3 to simulate basilar membrane compression. The resulting envelopes were lowpass filtered and downsampled to 200Hz, without any zero padding, resulting in a cochleagram representation of 211 frequency channels by 390 time points. This representation was the input to the auditory models. Cochleagram generation was implemented in PyTorch (code available: https://github.com/jenellefeather/chcochleagram).

##### SpectroTemporal model

For comparison to previous hand-engineered models of the auditory system we included a single layer SpectroTemporal model based on Chi et al., (2005). The main difference was that spectral filters were specified in cycles/erb (rather than cycles/octave) as the input signal to the model is a cochleagram with ERB-spaced filters. The model consists of a linear filter bank tuned to spectrotemporal modulations at different frequencies, spectral scales, and temporal rates. The different frequencies were implemented via applying the spectrotemporal filters as a 2D convolution with zero padding in frequency (800 samples) and time (211 samples). Spectrotemporal filters were constructed with center frequencies for the spectral modulations of [0.0625, 0.125, 0.25, 0.5, 1, 2] cycles/erb. Center frequencies for the temporal modulations consisted of [0.5, 1, 2, 4, 8, 16, 32, 64] and both upward and downward frequency modulations were included (resulting in 96 filters). An additional 6 purely spectral and 8 purely temporal modulation filters were included for a total of 110 modulation filters. To extract the power in each frequency band for each filter, we squared the output of each filter response at each time step and took the average across time for each frequency channel, similar to previous studies ^31, 45, 57^. These power measurements were used as the regressors for voxel and component modeling (23421 activations).

##### CochCNN9 architecture

The CochCNN9 architecture is based on the architecture in Kell et al., (2018) that emerged from a neural network architecture search. The architecture used here differed in that the input to the first layer of the network is maintained as the 211×390 size cochleagram rather than being reshaped to 256×256. The convolutional layer filters and pooling regions were adjusted from those of the Kell et al. architecture to maintain the same receptive field size in frequency and time given the altered input dimensions. The other difference was that the network here was trained with batch normalization rather than the local response normalization used in Kell et al., (2018). Along with the CochResNet50, this architecture was used for task-optimization comparisons throughout the paper.

The CochCNN9 architecture is denoted below with the sizes of the tensors propagated through the network denoted in parentheses. Model stages that were used for voxel and component response modeling are denoted in bold.

|  |  |
| --- | --- |
| Input (cochleagram) | (1,211,390) |
| BatchNorm2d_1 (1) | (1,211,390) |
| Conv2d_1(1, 96, kernel_size = [7, 14], stride = [3, 3], padding = 'same') | (96, 71, 130) |
| **ReLU_1** | **(96, 71, 130)** |
| **MaxPool2d_1(kernel_size = [2,5] , stride = [2,2], padding = 'same')** | **(96, 36, 65)** |
| BatchNorm2d_2(96) | (96, 36, 65) |
| Conv2d_2(96, 256, kernel_size = [4,8], stride = [2,2], padding = 'same') | (256, 18, 33) |
| **ReLU_2** | **(256, 18, 33)** |
| **MaxPool2d_2(kernel_size = [2,5] , stride = [2,2], padding = 'same')** | **(256, 9, 17)** |
| BatchNorm2d_3 (256) | (256, 9, 17) |
| Conv2d_3 (512, kernel_size = [2,5], stride = [1,1], padding = 'same') | (512, 9, 17) |
| **ReLU_3** | **(512, 9, 17)** |
| Conv2d_4 (1024, kernel_size = [2,5], stride = [1,1], padding = 'same') | (1024, 9, 17) |
| **ReLU_4** | **(1024, 9, 17)** |
| Conv2d_5 (512, kernel_size = [2,5], stride = [1,1], padding = 'same') | (512, 9, 17) |
| **ReLU_5** | **(512, 9, 17)** |
| **AvgPool_1 (kernel_size = [2,5] , stride = [2,2], padding = 'same')** | **(512, 5, 9)** |
| Linear_1 | (4096) |
| **ReLU_6** | **(4096)** |
| **Dropout_1 (p=0.5)** | **(4096)** |
| Linear_2 | (num_classes) |

where num_classes corresponds to the number of logits used for training each task (Table 1).

Thus, for CochCNN9, we extracted model representations from the following 10 layers with the number of unit activations (regressors) for each sound denoted in parentheses:

Cochleagram (211), ReLU_1 (6816), MaxPool2d_1 (3456), ReLU_2 (4608), MaxPool2d_2 (2304), ReLU_3 (4608), ReLU_4 (9216), ReLU_5 (4608), AvgPool_1 (2560), ReLU_6 (4096).

##### CochResNet50 architecture

The CochResNet50 model is composed of a ResNet50 backbone architecture applied to a cochleagram representation (with 2D convolutions applied to the cochleagram). Along with CochCNN9, this architecture was used for task-optimization comparisons throughout the paper.

The CochResNet50 architecture is denoted below with the sizes of the tensors propagated through the network denoted in parentheses. Model stages that were used for voxel and component response modeling are denoted in bold.

The model architecture follows:

|  |  |
| --- | --- |
| Input (cochleagram) | (1,211,390) |
| Conv2d_1(1, 64, kernel_size=7, stride=2, padding=3, bias=False) | (64, 106, 195) |
| BatchNorm2d_1(64) | (64, 106, 195) |
| **ReLU_1** | **(64, 106, 195)** |
| **MaxPool2d_1(kernel_size=3, stride=2, padding=1)** | **(64, 53, 98)** |
| **ResNetBlock_1(inplanes=64, planes=64, num_blocks=3, stride=1)** | **(256, 53, 98)** |
| **ResNetBlock_2(inplanes=256, planes=128, num_blocks=4, stride=2)** | **(512, 27, 49)** |
| **ResNetBlock_3(inplanes=512, planes=256, num_blocks=6, stride=2)** | **(1024, 14, 25)** |
| **ResNetBlock_4(inplanes=1024, planes=512, num_blocks=3, stride=2)** | **(2048, 7, 13)** |
| AvgPool_1 | (2048, 1, 1) |
| Linear_1 | (num_classes) |

where num_classes corresponds to the number of logits used for training each task (Table 1) and the ResNetBlock components of the architecture have the following structure:

|  |
| --- |
| 1. input (x) |
| 2. 1x1 Conv2d(inplanes, planes, stride=1) |
| 3. BatchNorm2d(planes) |
| 4. ReLU |
| 5. 3x3 Conv2d (planes, planes, stride=1) |
| 6. BatchNorm2d(planes) |
| 7. ReLU |
| 8. 1x1 Conv2d (planes, planes * expansion, stride=1) |
| 9. BatchNorm2d(planes) |
| 10. Residual connection on x (if inplanes !=planes * expansion): 1x1 Conv2D (inplanes, planes * expansion, stride) |
| 11. Residual connection on x (if inplanes !=planes * expansion): BatchNorm2d(planes * expansion) |
| 12. Add output from (9) to output from (11) |
| 13. (Output) ReLU |

Multiple of these residual blocks (num\_blocks) are stacked together to form a single ResNetBlock. The expansion factor was set to four for all layers (expansion=4).

Thus, for CochResNet50, we extracted model representations from the following 8 layers with the number of unit activations (regressors) for each sound denoted in parentheses:

Cochleagram (211), ReLU_1 (6784), MaxPool_1 (3392), ResNetBlock_1 (13568), ResNetBlock_2 (13824), ResNetBlock_3 (14336), ResNetBlock_4 (14336), AvgPool_1 (2048).

##### Training dataset for CochResNet50 and CochCNN9 models – Word, Speaker, and AudioSet tasks

Eight in-house models were trained on the Word-Speaker-Noise (WSN) dataset. This dataset was first presented in Feather et al., (2019)^25^ and was constructed from existing speech recognition and audio event classification datasets. The dataset description that follows is reproduced from Feather et al., (2022)^27^ with some additions to further detail the speaker and audio event recognition tasks.

The dataset was approximately balanced to enable performance of three tasks on the same training exemplar: (1) recognition of the word at the center of a two second speech clip (2) recognition of the speaker and (3) recognition of audio events, that were superimposed with the speech clips (serving as “background noise” for the speech tasks while enabling an audio event recognition task).

The speech clips used in the dataset were excerpted from the Wall Street Journal^169^ (WSJ) and Spoken Wikipedia Corpora^170^ (SWC). To choose speech clips, we screened WSJ, TIMIT^171^ and a subset of articles from SWC for appropriate audio clips (specifically, clips that contained a word at least four characters long and that had one second of audio before the beginning of the word and after the end of the word, to enable the temporal jittering augmentation described below). Some SWC articles were left out of the screen due to a) potentially offensive content for human listening experiments; (29/1340 clips), b) missing data; (35/1340 clips), or c) bad audio quality (for example, due to computer generated voices of speakers reading the article or the talker changing mid-way through the clip; 33/1340 clips). Each segment was assigned the word class label of the word overlapping the segment midpoint and a speaker class label determined by the speaker. With the goal of constructing a dataset with speaker and word class labels that were approximately independent, we selected words and speaker classes such that the exemplars from each class spanned at least 50 unique cross-class labels (e.g., 50 unique speakers for each of the word classes). This exclusion fully removed TIMIT from the training dataset. We then selected words and speaker classes that each contained at least 200 unique utterances, and such that each class could contain a maximum of 25% of a single cross-class label (e.g., for a given word class, a maximum of 25% of utterances could come from the same speaker). These exemplars were subsampled so that the maximum number in any word or speaker class was less than 2000. The resulting training dataset contained 230,356 unique clips in 793 word classes and 432 speaker classes, with 40,650 unique clips in the test set. Each word class had between 200 and 2000 unique exemplars. A “null” class was used as a label for the word and speaker when a background clip was presented without the added speech.

The audio event clips that were superimposed on the speech clips were a subset of examples from the “Unbalanced Train” split of the AudioSet dataset (a set of annotated YouTube video soundtracks)^55^ To minimize ambiguity for the two speech tasks, we removed any sounds under the “Speech” or “Whispering” branch of the AudioSet ontology. Since a high proportion of AudioSet clips contain music, we achieved a more balanced set by excluding any clips that were only labeled with the root label of “Music”, with no specific branch labels. We also removed silent clips by first discarding everything tagged with a “Silence” label and then culling clips containing more than 10% zeros. This screening resulted in a training set of 718,625 unique natural sound clips spanning 516 categories. Each AudioSet clip was a maximum of 10 seconds long, from which a 2-second excerpt was randomly cropped during training (see below). A “null” audio event label was used as a label when speech clips were presented without added background sound.

During training, the speech clips from the Word-Speaker-Noise dataset were randomly cropped in time and superimposed on random crops of the AudioSet clips. Data augmentations during training consisted of 1) randomly selecting a clip from the pre-screened AudioSet clips to pair with each labeled speech clip, 2) randomly cropping 2 seconds of the AudioSet clip and 2 seconds of the speech clip, cropped such that the labeled word remained in the center of the clip (due to training pipeline technicalities, we used a pre-selected set of 5,810,600 paired speech and natural sound crops which spanned 25 epochs of the full set of speech clips and 8 passes through the full set of AudioSet clips), 3) superimposing the speech and the noise (i.e., the AudioSet crop) with a Signal-to-Noise-Ratio (SNR) sampled from a uniform distribution between −10dB SNR and 10dB SNR, augmented with additional samples of speech without an AudioSet background (i.e., with infinite SNR, 2464 examples in each epoch) and samples of AudioSet without speech (i.e., with negative infinite SNR, 2068 examples in each epoch) and 4) setting the root-mean-square (RMS) amplitude of the resulting signal to 0.1. By constructing the dataset in this way, we could train networks on different tasks while using the same dataset and training and test augmentations.

Evaluation performance for the word and speaker recognition tasks was measured from one pass through the speech test set (i.e., one crop from each of the 40,650 unique test set speech clips) constructed with the same augmentations used during training (specifically, variable SNR and temporal crops, paired with a set of AudioSet test clips from the “Balanced Train” split, same random seed used to test each model such that test sets were identical across models).

The representation from the AudioSet-trained models were evaluated with a support vector machine (SVM) fit to the ESC-50 dataset^172^, composed of 50 types of environmental sounds. After the model was trained, and for each of the five folds in ESC-50, an SVM was fit to the output representation of the top of the layer immediately before the final linear layer (AvgPool_1 for CochResNet50 and ReLU_6 for CochCNN9). Each fold had 400 sounds, resulting in 1600 sounds used for training when holding out each fold. As the networks were trained with two-second-long sound clips, we took random two-second crops of the ESC-50 sounds. For each sound in the training and test data, we took 5 two-second-long crops at random from the five second sound (randomly selecting a new crop if the chosen crop was all zeros). The five crops of the training data were all used in fitting the SVM, treated as separate training data points. After the predictions were measured for the five crops for each test sound, we chose the label that was predicted most often as the prediction for the test sound.

The SVM was implemented with sklearn’s LinearSVC, with cross validation over five regularization parameters (C=[0.01, 0.1, 1.0, 10.0, 100.0]). For cross validation, a random selection of 25% of the training sounds were held out and the SVM was fit on the other 75% of the sounds, and this was repeated three times (the five crops from a given sound were never split up between cross validation training and test splits, such that the cross validation tested for generalization to held-out sounds). This cross-validation strategy is independent of the held-out test fold, as it only relies on the training dataset. A best regularization parameter was determined by choosing the parameter that resulted in the maximum percent correct averaged across the three splits, and we refit the SVM using the selected regularization parameter on the entire training dataset of 1600 sounds to measure the performance on the held-out fold (400 sounds). The reported performance is the average across the 5 folds of the ESC-50 dataset.

##### Training CochResNet50 and CochCNN9 models – Word, Speaker, and AudioSet tasks

Each audio model was trained for 150 epochs of the speech dataset (corresponding to 48 epochs of the AudioSet training data). The learning rate was decreased by a factor of 10 after every 50 speech epochs (16 AudioSet epochs). All models were trained on the OpenMind computing cluster at MIT using NVIDIA GPUs.

The Word and Speaker networks were trained with a cross entropy loss on the target labels. Because the AudioSet dataset has multiple labels per clip, the logits are passed through a sigmoid and the Binary Cross Entropy is used as the loss function. Models had weight decay of 1e-4, except for models trained on the AudioSet task (including the multi-task models) which had weight decay of 0.

Both of the CochResNet50 and CochCNN9 architectures were trained simultaneously on all three tasks by including three fully connected layers as the final readout. These models were optimized by adding together a weighted loss from each individual task, and minimizing this summed loss. The weights used for the loss function were 1.0 (Word), 0.25 (Speaker), and 300 (AudioSet).

Additional training details are given in the table below.

| Model Name | Batch Size | Initial Learning Rate | Num Classes (includes “null”) | Accuracy on Training Task |
| --- | --- | --- | --- | --- |
| CochCNN9 Word | 128 | 0.01 | 794 | (Top 1) 66.640%<br>(Top 5) 83.102% |
| CochCNN9 Speaker | 128 | 0.01 | 433 | (Top 1) 96.216%<br>(Top 5) 99.058% |
| CochCNN9 AudioSet | 128 | 0.00001* | 517 | (ESC-50 SVM) 83.60% |
| CochCNN9 MultiTask | 128 | 0.00001* | Three tasks:<br>(Word) 794,<br>(Speaker) 433,<br>(AudioSet) 517 | (Top 1 Word) 64.954%<br>(Top 5 Word) 81.998%<br>(Top 1 Speaker) 86.686%<br>(Top 5 Speaker) 96.039%<br>(ESC-50 SVM) 82.60% |
| CochResNet50 Word | 256 | 0.1 | 794 | (Top 1) 86.792%<br>(Top 5) 95.149% |
| CochResNet50 Speaker | 256 | 0.1 | 433 | (Top 1) 99.114%<br>(Top 5) 99.835% |
| CochResNet50 AudioSet | 256 | 0.001* | 517 | (ESC-50 SVM) 91.6% |
| CochResNet50 MultiTask | 256 | 0.001* | Three tasks:<br>(Word) 794,<br>(Speaker) 433,<br>(AudioSet) 517 | (Top 1 Word) 83.459%<br>(Top 5 Word) 93.422%<br>(Top 1 Speaker) 94.354%<br>(Top 5 Speaker) 98.785%<br>(ESC-50 SVM) 87.450% |

\* Models trained with the AudioSet loss had additional gradient clipping ( $\max \| \cdot \|_2 = 1.0$ ) and learning rate warm-up for the first 500 batches of training (learning rate = $\langle \text{initial learning rate} \rangle / (500 - i)$ , where $i$ is the batch number).

##### Training dataset for CochResNet50 and CochCNN9 models - musical genre task

The genre task was the 41-way classification task introduced by Kell et al., (2018). The sounds and labels were derived from The Million Song Dataset^155^. Genre labels were obtained from user-generated ‘‘tags’’ from the MusicBrainz open-source music encyclopedia (https://musicbrainz.org/). Tags were first culled to eliminate those that did not apply to at least ten different artists or that did not obviously correspond to a genre. These tags were then grouped into genre classes using hierarchical clustering applied to the tag co-occurrence matrix, grouping together tags that overlapped substantially. See Table S2 from the Kell et al., (2018) for a list of genres and the tags associated with each genre.

Training exemplars for the genre task were obtained by randomly excerpting two-second clips from the tracks that had tags for the genre labels selected for the task. The music excerpts were superimposed with two-second excerpts of one of four different background noises: (1) auditory scenes, (2) two-speaker speech babble, (3) eight-speaker speech babble, or (4) music-shaped noise. Music-shaped noise consisted of a two-second clip of noise that was matched to the average spectrum of its corresponding two-second clip of music. SNRs were selected to yield performance in human listeners that was below ceiling (but above chance). The mean SNR for each of the four background types was 12 dB, with the SNR for each training example drawn randomly from a Gaussian with a standard deviation of 2 dB. All waveforms were downsampled to 16 kHz.

##### Training CochResNet50 and CochCNN9 models - musical genre task

The genre networks were trained with a cross entropy loss, and. A stochastic gradient descent optimizer was used for training with weight decay of 1e-4, momentum of 0.9, and an initial learning rate of 0.01. The models were trained for 125 epochs of the genre dataset, and the learning rate was dropped by a factor of 10 after every 50 epochs. A batch size of 64 was used for training. The CochCNN9 architecture achieved Top 1 accuracy of 83.21% and Top 5 of 96.19% on the musical genre task, and the CochResNet50 model achieved Top 1 accuracy of 87.99% and Top 5 accuracy of 97.56%.

### Models trained on clean speech

Models trained on clean speech used the speech clips from the Word-Speaker-Noise dataset^25^ without using the associated AudioSet backgrounds. Data augmentations during training consisted of 1) pseudorandomly selecting a labeled speech clip as in the original dataset, 2) randomly cropping 2 seconds of the speech clip, cropped such that the labeled word remained in the center of the clip, 3) setting the root-mean-square (RMS) amplitude of the resulting signal to 0.1. Each clean speech model was trained for 150 epochs of the speech dataset. The learning rate was decreased by a factor of 10 after every 50 speech epochs. All models were trained on the OpenMind computing cluster at MIT using NVIDIA GPUs. The Word and Speaker networks were trained with a cross entropy loss on the target labels.

Evaluation performance for the word and speaker recognition tasks was measured from one pass through the speech test set (i.e., one crop from each of the 40,650 unique test set speech clips) constructed with the 2 second temporal crop and RMS normalization used during training (same random seed used to test each model such that test sets were identical across models). Additional details of model training and performance are given in the table below.

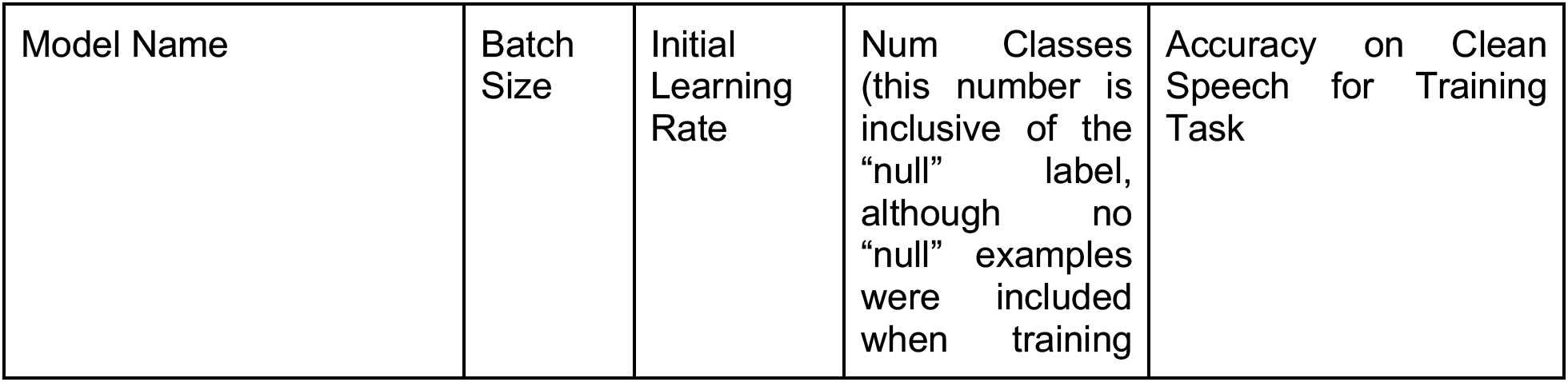
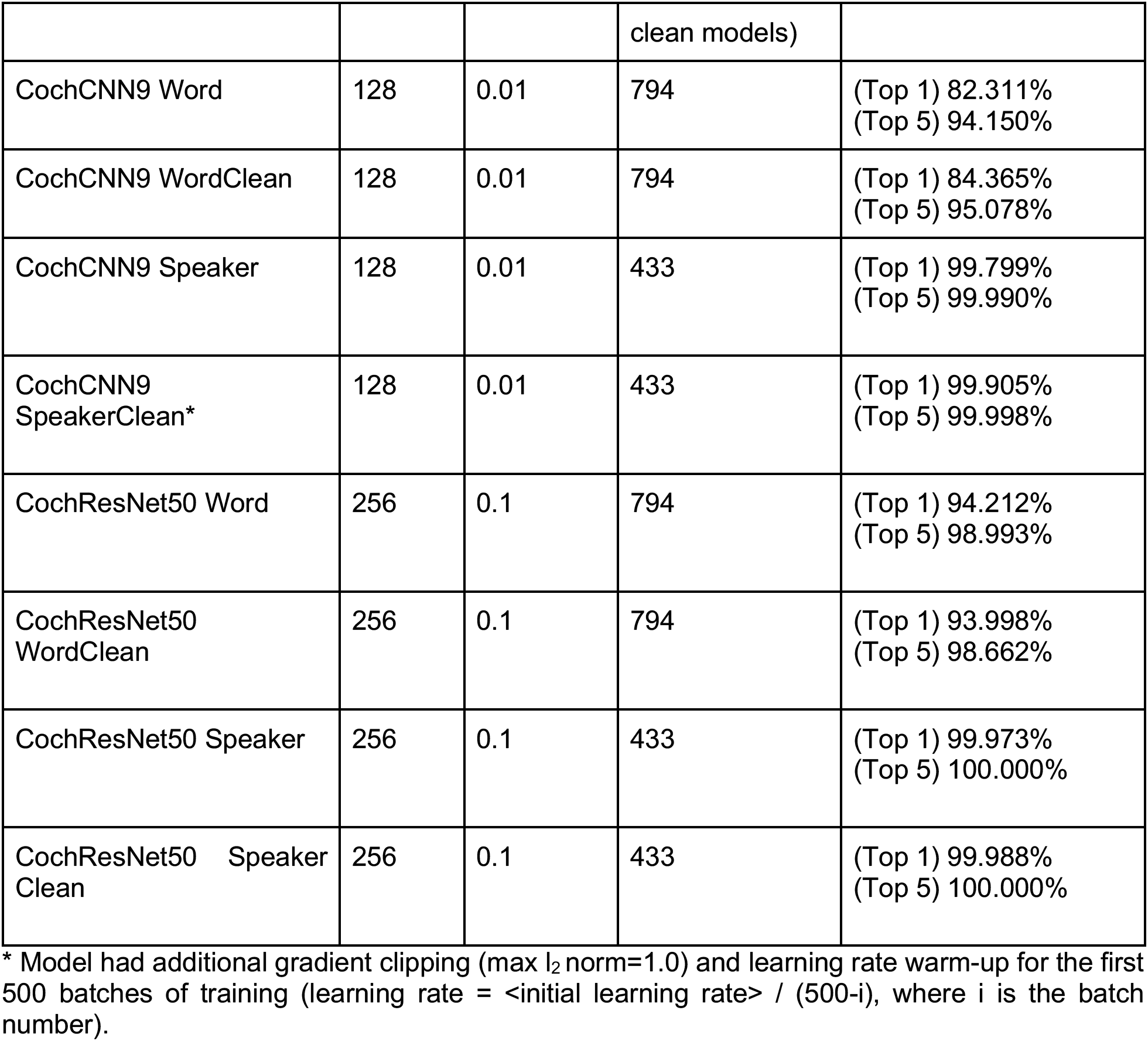

### Candidate models with permuted weights

In addition to the trained networks, we also analyzed ‘permuted’ versions of the models with the exact same architecture as the trained models. We created these models by replacing all parameters making up the trained model in each network by random permutations across all tensor dimensions within a given parameter block (e.g., a weight or bias matrix) for each model stage. This model manipulation destroyed the parameter structure learned during task-optimization, while preserving the marginal statistics of the parameters. All analyses procedures were identical for trained and permuted networks.

## Acknowledgements

We thank Ian Griffith for training the music genre classification models, Alex Kell for helpful discussions, Nancy Kanwisher for sharing fMRI data, developers for making their trained models available for public use, and the McDermott lab for comments on an earlier draft of the paper. Work supported by NIH grant R01DC017970. G.T. was supported by the Amazon Fellowship from the Science Hub (administered by the MIT Schwarzman College of Computing) and the International Doctoral Fellowship from American Association of University Women (AAUW). J.F was supported by an MIT Friends of McGovern Institute Fellowship and a DOE Computational Science Graduate Fellowship (grant DE-FG02-97ER25308).

