## Supplementary Information (SI) for "Many but not all deep neural network audio models capture brain responses and exhibit correspondence between model stages and brain regions"

##### Overview

**Figure S1:** Representational Dissimilarity Matrices for fMRI
**Figure S2:** Median variance explained across model stages for each model
**Figure S3:** Model evaluation consistency for components between different random seeds
**Figure S4:** Surface maps of best-predicting model stage for trained models
**Figure S5:** Surface maps of best-predicting model stage for permuted control models
**Figure S6:** Median variance explained across model stages for each model separated into ROIs
**Figure S7:** Stage-region correspondence of permuted control networks
**Figure S8:** Component response variance explained by models trained in background noise versus no background noise
**Figure S9:** Effective dimensionality (ED) in relation to model-brain similarity metrics
**Figure S10:** Consistency between regression and representational similarity model-brain similarity metrics

**Table S1:** Natural sound stimulus set

### Supplementary Figure S1

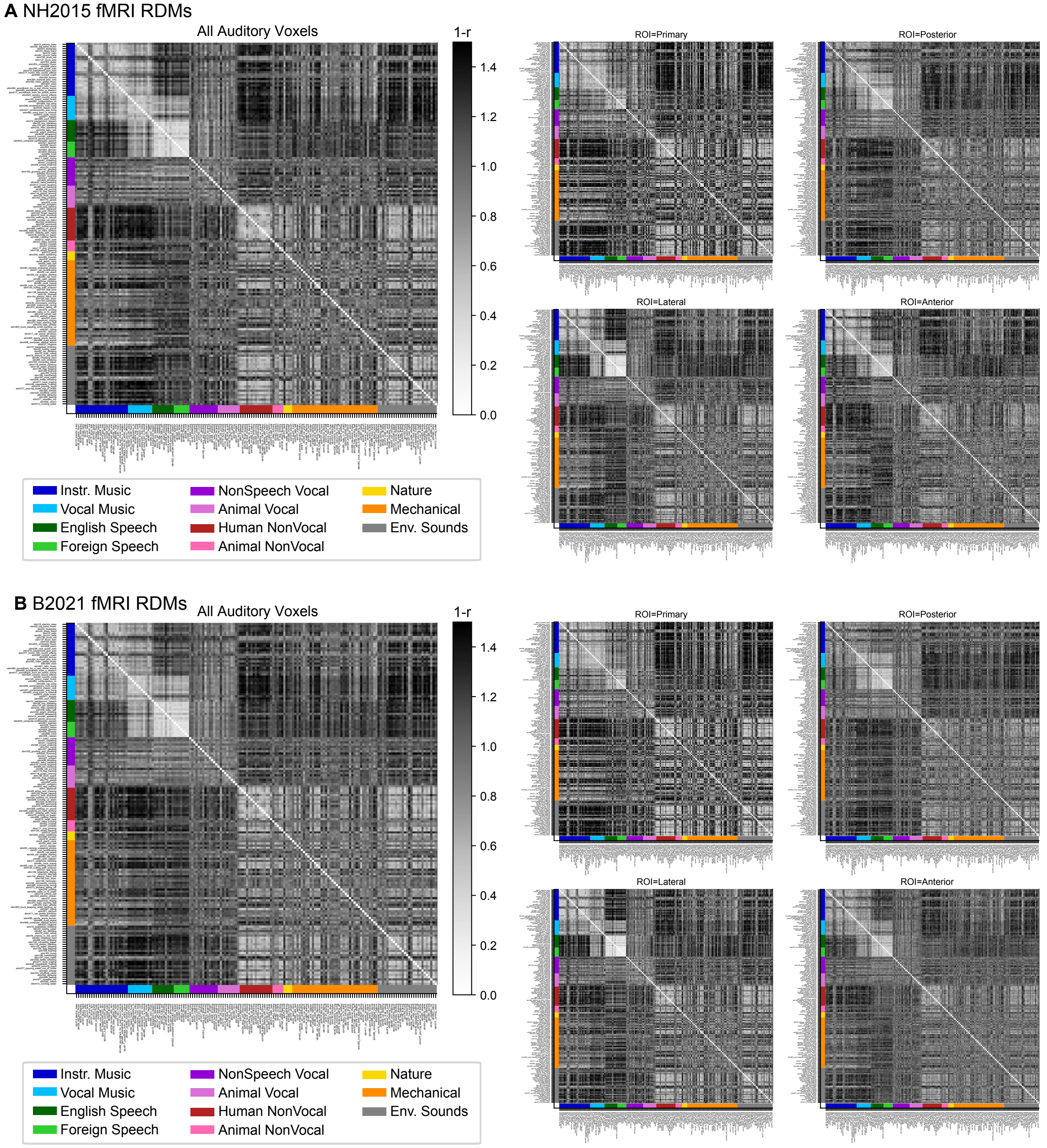


**Figure S1. Representational Dissimilarity Matrices for fMRI voxels in (A) NH2015 and (B) B2021.** For visualization purposes, the RDMs are computed as 1-Spearman Correlation between the 3-scan average BOLD responses for pairs of sounds. RDMs are computed for all sound-responsive voxels (left) and using only a subset of voxels for each of the anatomical ROIs (right). Sounds are grouped by sound categories (included in colors on the axis). Data and code with which to reproduce results are available at https://github.com/gretatuckute/auditory_brain_dnn.

### Supplementary Figure S2

**A** Predictivity across Model Stages

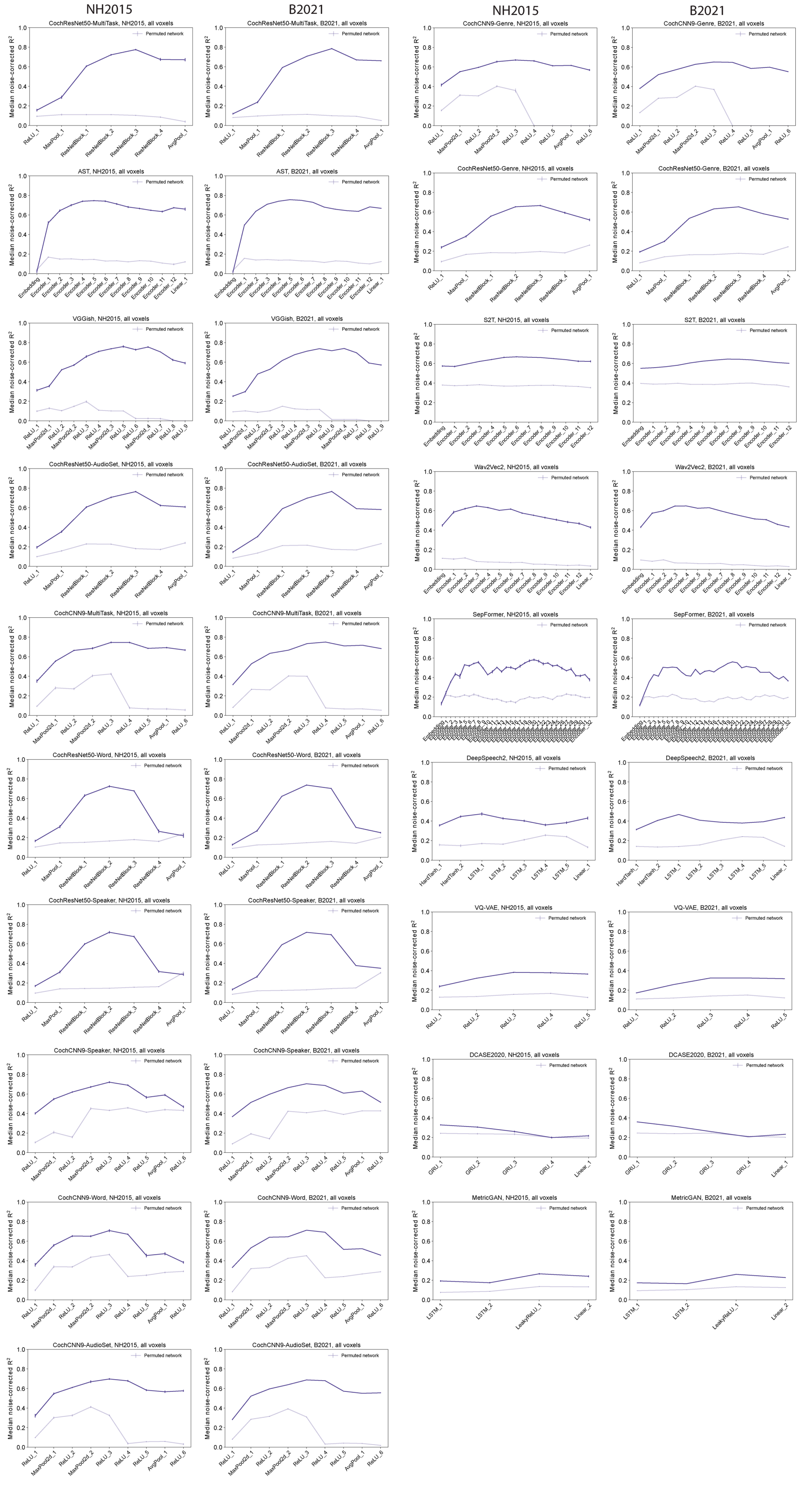

**Figure S2. Median variance explained across model stages for each model.** Explained variance was measured for each voxel and the aggregated median variance explained across all voxels in auditory cortex was obtained. This aggregated median variance explained is plotted for all candidate models (n=19) for both fMRI datasets. The model plots are sorted according to overall model performance (median noise-corrected $R^2$ for NH2015 in Figure 2A in the main text), meaning that the first subplot shows the best-performing model, CochResNet50-MultiTask, and the last subplot shows the worst-performing model, MetricGAN. Dark lines show the trained networks, and lighter lines show the control networks with permuted weights. Error bars are within-participant SEM. Error bars are smaller for the B2021 dataset because of the larger number of participants (20 vs. 8). We note that some of the variation in predictivity across model stages in the models with permuted weights could be driven by the receptive field sizes at different stages, which are partly a function of the model architecture. Data and code with which to reproduce results are available at https://github.com/gretatuckute/auditory_brain_dnn.

#### Supplementary Figure S3

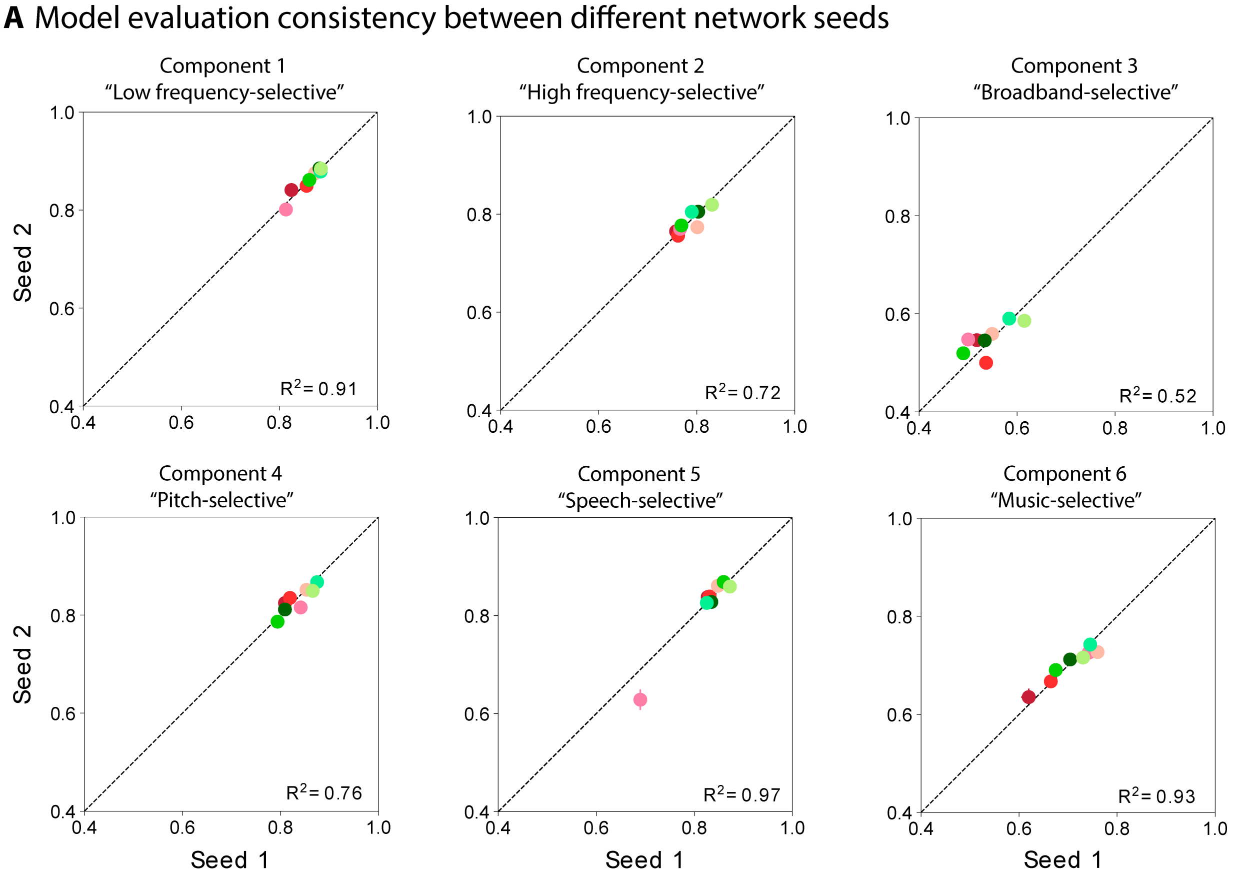


**Figure S3. Comparison of component variance explained by in-house models trained from different random seeds.** We trained the in-house models from two different random seeds. The variance explained for the first seed is plotted on the x-axis and for the second seed on the y-axis. Each data point represents a model using with the same color correspondence as in Figure 2 in the main text. Variance explained was obtained from the best-predicting stage of each model for each component, selected using independent data. Error bars are SEM over iterations of the model stage selection procedure (see Methods; Component modeling). Data and code with which to reproduce results are available at https://github.com/gretatuckute/auditory_brain_dnn.

#### **Supplementary Figure S4** (extension of Figure 6)

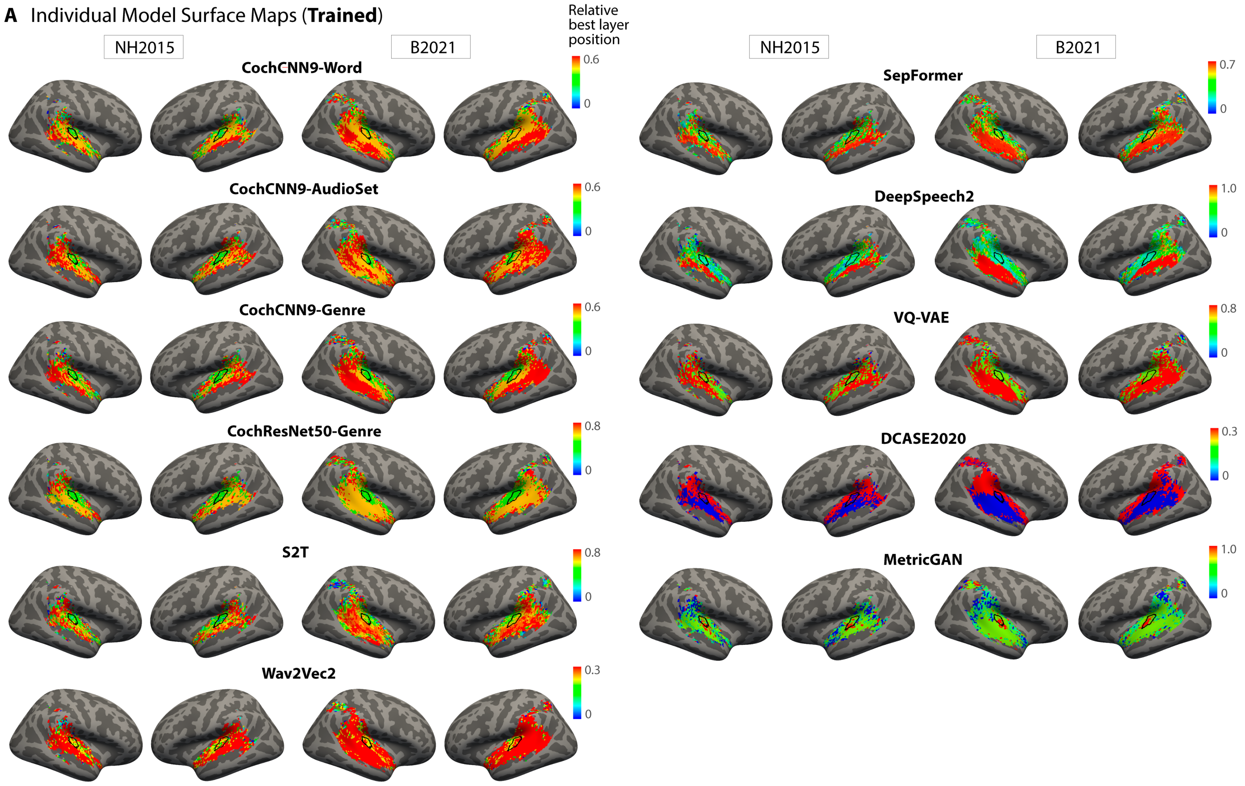


**Figure S4. Surface maps of best-predicting model stage for trained models.** The figure shows surface maps for trained models that are not included in Figure 6A in the main text (which featured the n=8 best-predicting models, leaving the n=11 models shown here). The plots are sorted according to overall model predictivity (the quantity plotted in Figure 2A in the main text). As in Figure 6A in the main text, the plots show the model stage that best predicts each voxel as a surface map (FsAverage) (median best stage across participants). We assigned each model stage a position index between 0 and 1. The color scale limits were set to extend from 0 to the stage beyond the most common best stage (across voxels). Data and code with which to reproduce results are available at https://github.com/gretatuckute/auditory_brain_dnn.

#### Supplementary Figure S5 (extension of Figure 6)

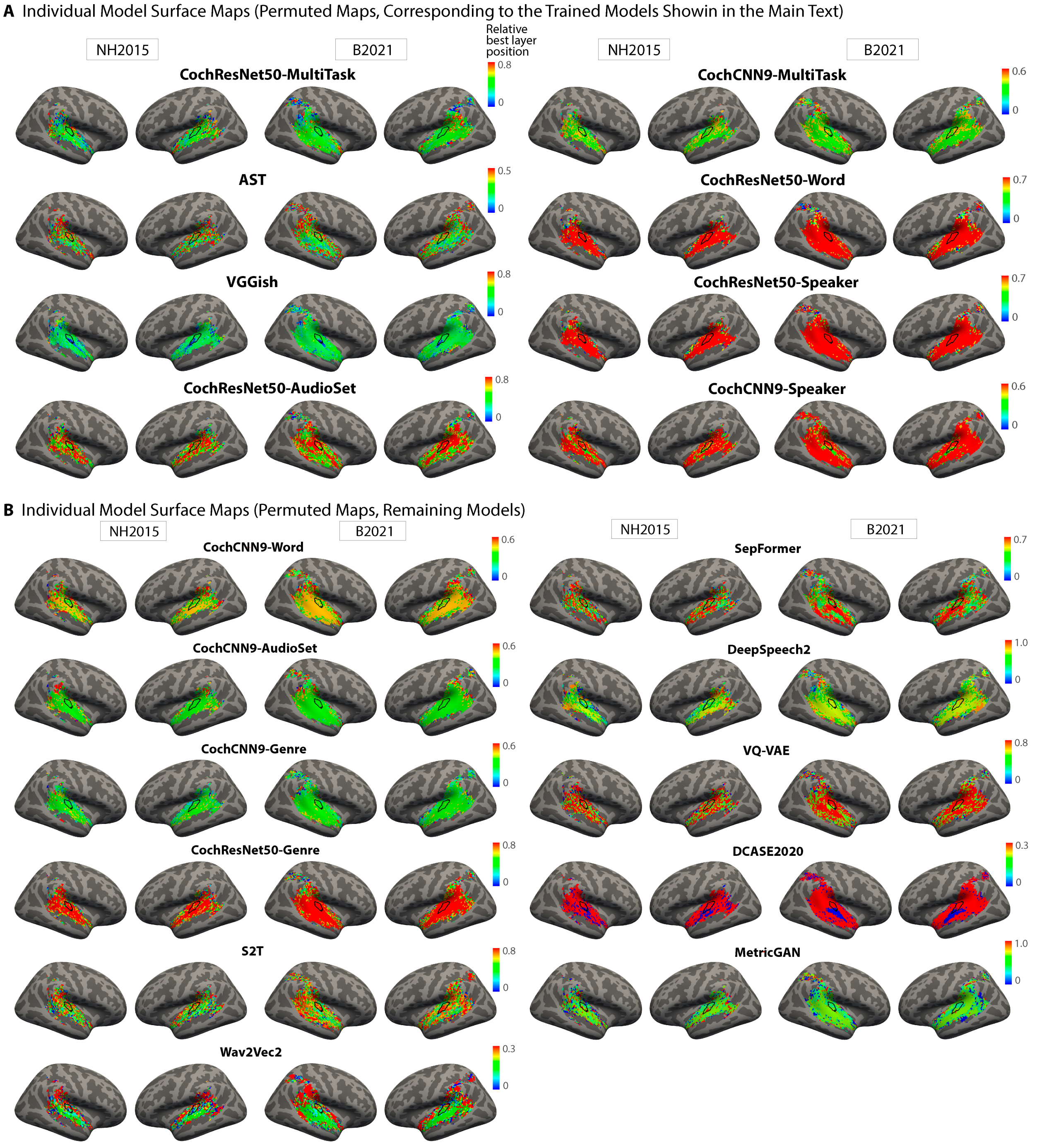


**Figure S5. Surface maps of best-predicting model stage for permuted control models.** Panel A shows the surface maps for the eight models shown in Figure 6A in the main text, but with permuted weights. Panel B shows surface maps for models with permuted weights that are not included in Figure 6A in the main text. Identical analyses procedures and color scale limits were used for the permuted models as for the trained ones. Data and code with which to reproduce results are available at https://github.com/gretatuckute/auditory_brain_dnn.

### Supplementary Figure S6

#### A Predictivity across Model Stages for Anatomical ROIs

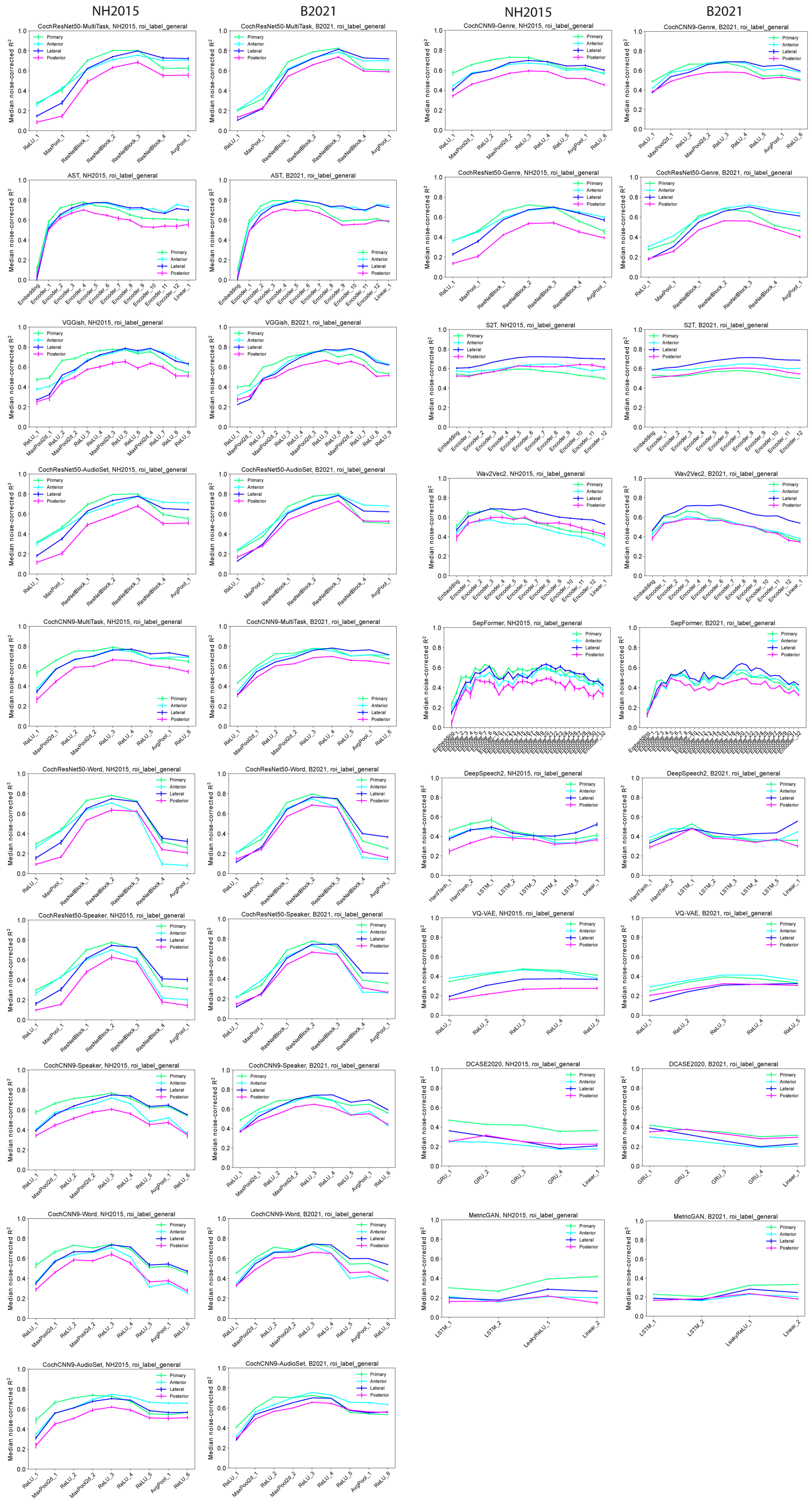

**Figure S6**. **Median variance explained by each model stage of each model for different auditory ROIs.** Explained variance was measured for each voxel and the aggregated median variance explained across each of the four anatomical ROIs (primary, anterior, lateral, posterior) was obtained. This aggregated median variance explained is plotted for all stages of all candidate models (n=19) for both fMRI datasets. The model plots are sorted according to overall model predictivity (median noise-corrected $R^2$ for NH2015 in Figure 2A in the main text; same model order as in Figure S2). Error bars are within-participant SEM. Error bars are smaller for the B2021 dataset because of the larger number of participants (20 vs. 8). Data and code with which to reproduce results are available at https://github.com/gretatuckute/auditory_brain_dnn.

#### **Supplementary Figure S7** (extension of Figure 7)

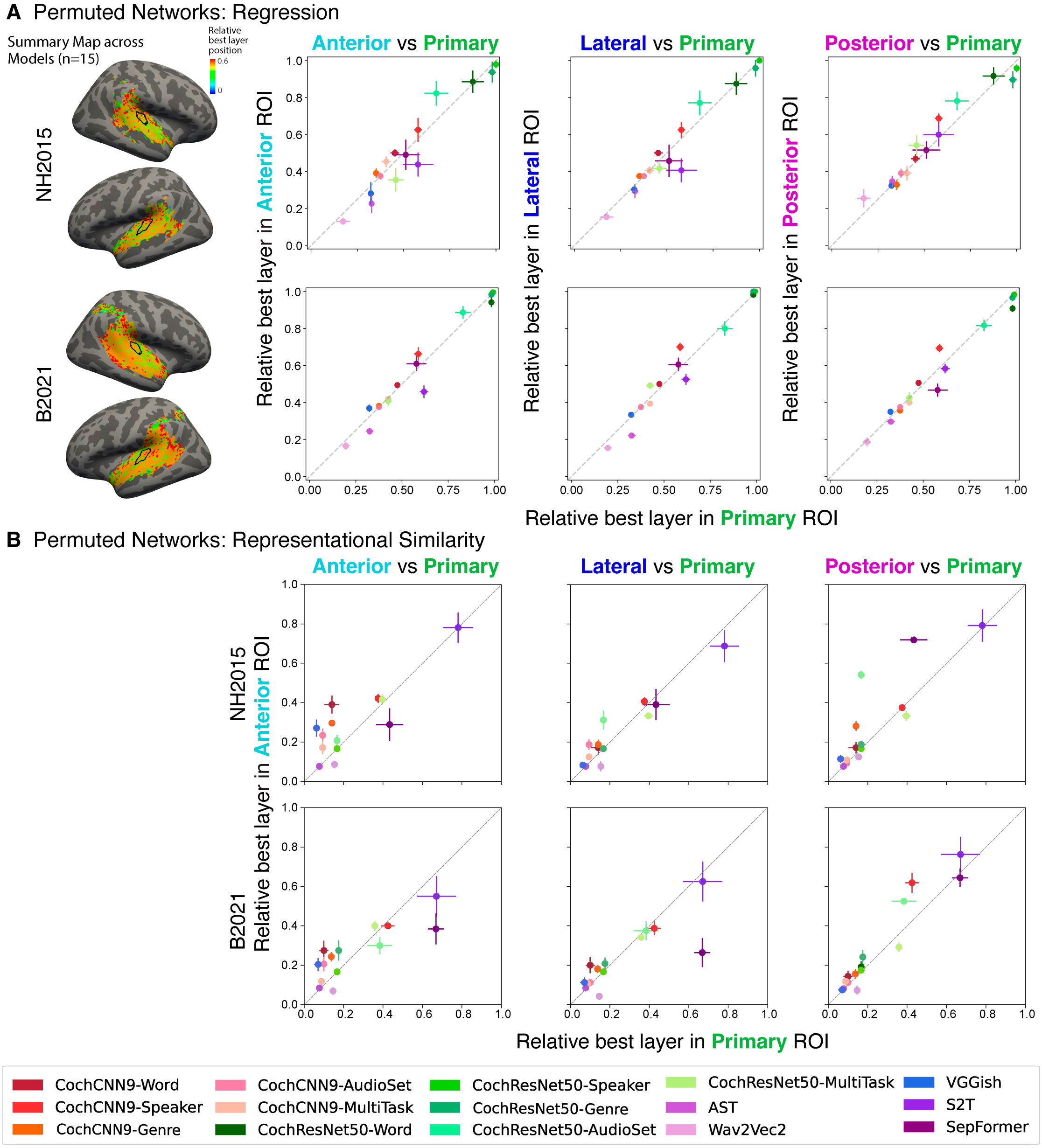


**Figure S7. Stage-region correspondence of permuted models.** This figure mirrors Figure 7 in the main text which shows the quantification of model-stage-region correspondence across trained models. **(A)** As in Figure 7 in the main text, we obtained the median best-predicting stage for each model within four anatomical ROIs (illustrated in Figure 7A, main text): primary auditory cortex (x axis in each plot in A and B) and anterior, lateral, and posterior non-primary regions (y axes in A and B). We performed the analysis on each of the two fMRI data sets, including each model that out-predicted the baseline model in Figure 2A in the main text (n=15 models). Each data point corresponds to a model with permuted weights, with the same color correspondence as in Figure 2 in the main text. None of the six possible comparisons (two datasets x three non-primary ROIs) were statistically significant even without correction for multiple comparisons, p>0.16 in all cases (Wilcoxon signed rank tests, two-tailed). **(B)** Same analysis as A but with the best-matching model stage determined by correlations between the model and ROI representational dissimilarity matrices. None of the six possible comparisons were statistically significant even without correction for multiple comparisons, p>0.07 in all cases (Wilcoxon signed rank tests, two-tailed). Data and code with which to reproduce results are available at https://github.com/gretatuckute/auditory_brain_dnn.

### Supplementary Figure S8

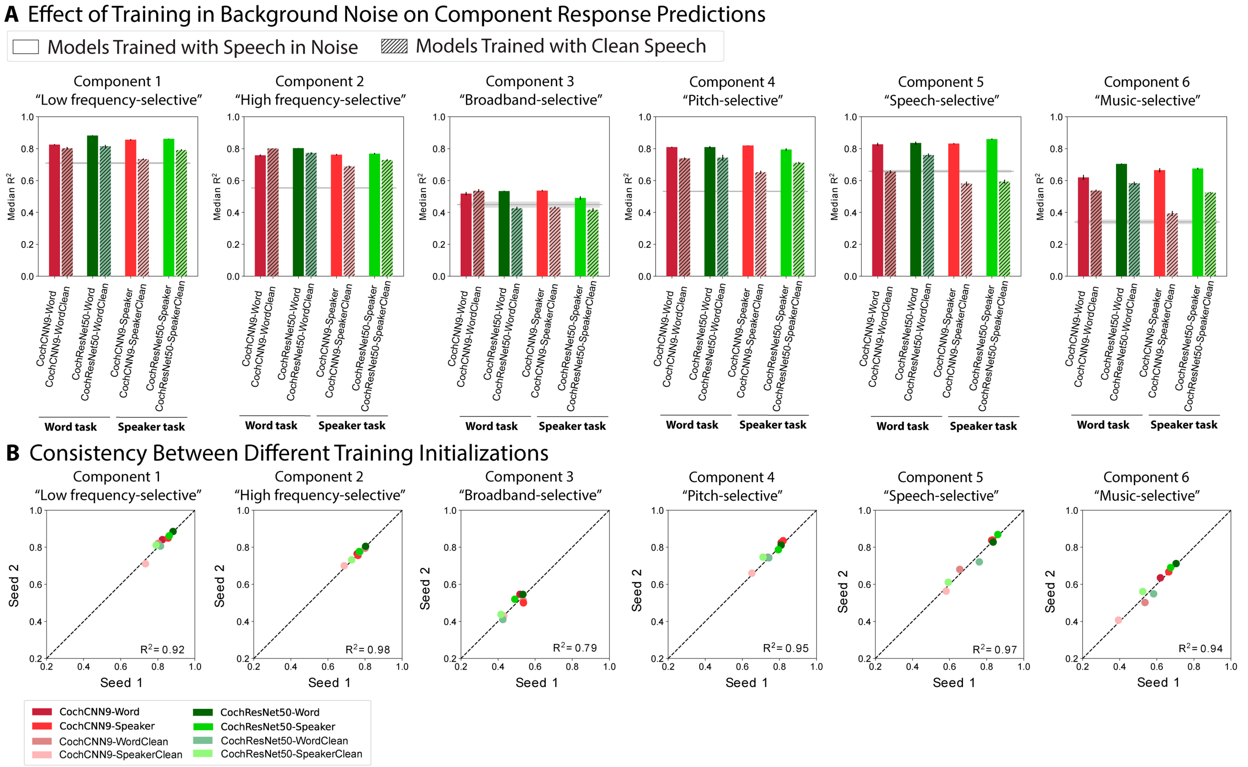


**Figure S8. Component response variance explained by models trained with and without background noise. (A)** Variance explained was obtained from the best-predicting stage of each model for each component, selected using independent data. Models trained in the presence of background noise are shown in the same color scheme as in Figure 2 in the main text; models trained with clean speech are shown with hashing. Grey line shows variance explained by the SpectroTemporal baseline model. Error bars are SEM over iterations of the model stage selection procedure (see Methods; Component modeling). **(B)** We trained the models from two different random seeds. The variance explained for the first seed is plotted on the x-axis and for the second seed on the y-axis. Each data point represents a model. Data and code with which to reproduce results are available at https://github.com/gretatuckute/auditory_brain_dnn.

### Supplementary Figure S9

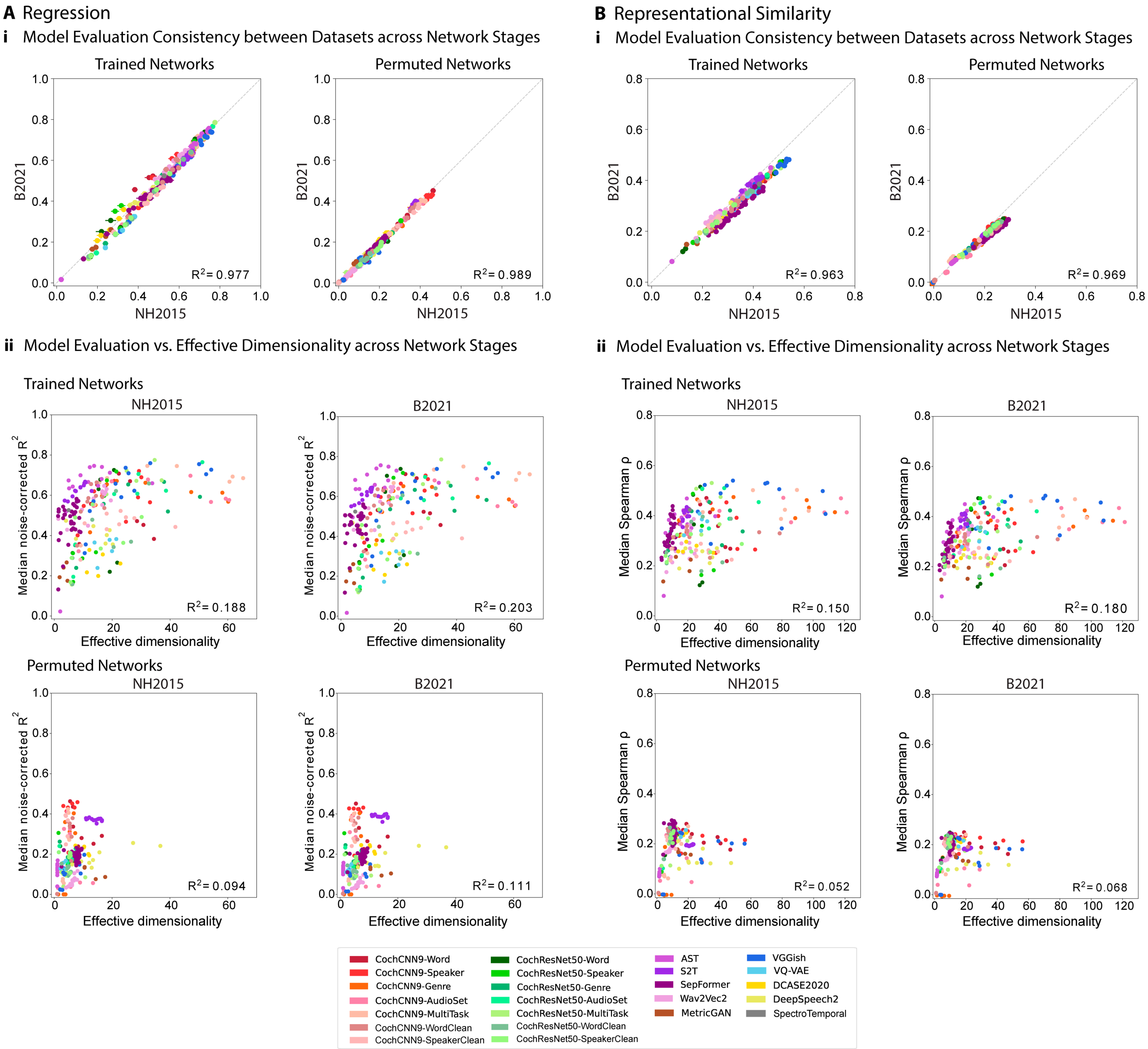


**Figure S9. Effective dimensionality in relation to model-brain similarity metrics. (A)** Effective dimensionality and regression-based model-brain similarity metric (voxelwise modeling). Panel **i** shows the consistency of the model evaluation metric (median noise-corrected $R^2$) between the two datasets analyzed in the paper (NH2015 and B2021). The consistency between datasets provides a ceiling for the strength of the relationship shown in panel **ii.** Panel **ii** shows the relationship between the model evaluation metric (median noise-corrected $R^2$) and effective dimensionality (computed as described in Methods; Effective dimensionality). Each data point corresponds to a model stage, with the same color correspondence as in Figure 2 in the main text. **(B)** Same analysis as A but with the representational similarity analysis evaluation metric (median Spearman correlation between the model and fMRI representational dissimilarity matrices). All unique models in the study were included in this analysis (n=20 models in Figure 2 in the main text plus n=4 models trained on the word and speaker tasks without background noise from Figure 8 in the main text, i.e., n=24 models in total). Data and code with which to reproduce results are available at https://github.com/gretatuckute/auditory_brain_dnn.

#### Supplementary Figure S10

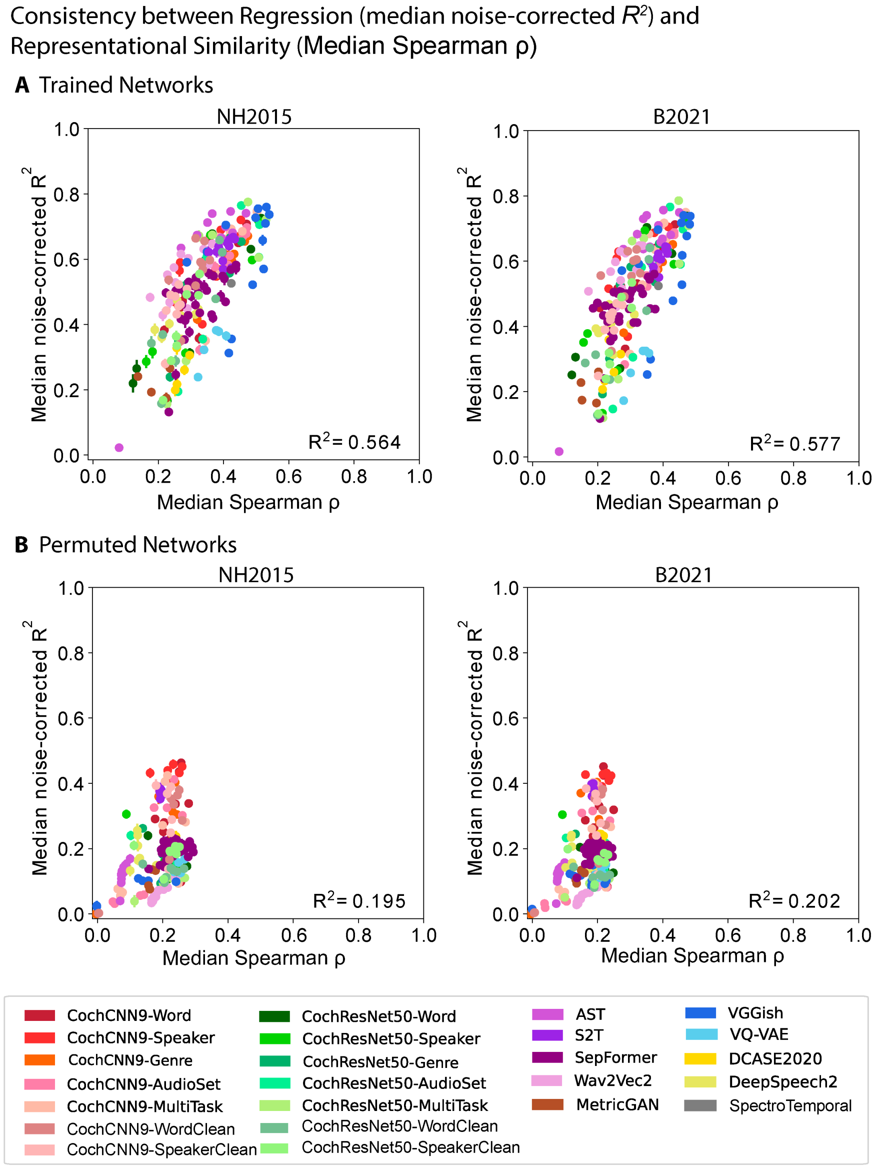


**Figure S10. Consistency between regression and representational similarity model-brain similarity metrics. (A)** Correlation between the regression-based metric (median noise-corrected noise-corrected $R^2$) and the representational similarity metric (median Spearman correlation) across trained network stages for the NH2015 and B2021 datasets. Each data point corresponds to a network stage, with the same color correspondence as in Figure 2 in the main text. **(B)** Same as panel A, but for permuted network stages. All unique models in the study were included in this analysis (n=20 models in Figure 2 in the main text plus n=4 models trained on the word and speaker tasks without background noise from Figure 8 in the main text, i.e., n=24 models in total). Data and code with which to reproduce results are available at https://github.com/gretatuckute/auditory_brain_dnn.

#### Supplementary Table S1

##### Mechanical
alarm clock
bike bell
blender
camera snapping photos
car accelerating
car alarm
car engine starting
car horn
cash register
phone vibrating
clock ticking
coin in vending machine
cutting with scissors
dial tone
telephone dialing
doorbell
electric hand drill
hair dryer
helicopter
microwave running
motorcycle revving
airplane taking off
printing
radio or tv static
school bell
electric shaver
siren
telephone ringing
train warning bell
train whistle
truck beeping while backing up
typing
vacuum
car power windows
zipper
sports arena buzzer
busy signal
computer startup sound
ringtone

##### Nature
wind
water splashing
thunder
stream

##### Environmental
basketball dribbling
boiling water
car skidding
chair rolling
chimes in the wind
chopping food
coin dropping
crumpling paper
dishes clanking
water dripping
flag flapping
flushing
frying
hammering
road traffic
kettle whistling
keys jingling
newspaper rustling
pouring liquid
pouring water out of bottle
whistle
shuffling cards
spraying
tearing
squeaky toy
velcro
running water

##### Animal non-vocalization
bees buzzing
cicadas
crickets
dog drinking
wings flapping

##### Animal vocalization
cat meowing
cat purring
dog barking
puppy whining
duck quack
frog croaking
geese
crow
songbird
dog panting

##### Human non-vocal
applause
biting & chewing
finger tapping
door knocking
walking on leaves
running up stairs
scratching
swimming
toothbrushing
walking on gravel
walking on hard surface
walking with heels
writing on paper
rubbing hands
heart beat

##### Human non-speech vocalization
person screaming
baby crying
breathing
coughing
crowd cheering
baby crying
gargling
grunting & groaning
humming
laughing
whistling
baby babbling
crowd laughing

##### English speech
background speech
boy speaking
girl speaking
man speaking
baby talk
angry shouting
whispering
woman speaking
sports announcer
computer-synthesized speech

**Table S1. Natural sound stimulus set.** List of all 165 sounds presented to human listeners while in the fMRI machine. Category assignments were based on judgments of human subjects on Amazon Mechanical Turk. Source data originally published in: Norman-Haignere, S. V., Kanwisher, N. G. & McDermott, J. H. (2015) Distinct cortical pathways for music and speech revealed by hypothesis-free voxel decomposition. *Neuron* **88**, 1281–1296.
